# Personality traits vary in their association with brain activity across situations

**DOI:** 10.1101/2024.04.18.590056

**Authors:** Samyogita Hardikar, Brontë McKeown, Adam Turnbull, Ting Xu, Sofie L. Valk, Boris C. Bernhardt, Daniel S. Margulies, Michael P. Milham, Elizabeth Jefferies, Robert Leech, Arno Villringer, Jonathan Smallwood

## Abstract

Human cognition supports complex behaviour across a range of situations, and traits (such as personality) influence how we react in these different contexts. Although viewing traits as situationally grounded is common in social sciences it is often overlooked in neuroscience. Often studies focus on linking brain activity to trait descriptions of humans examine brain-trait associations in a single task, or, under passive conditions like wakeful rest. These studies, often referred to as brain wide association studies (BWAS) have recently become the subject of controversy because results are often unreliable even with large sample sizes. Although there are important statistical reasons why BWAS yield inconsistent results, we hypothesised that results are inconsistent because the situation in which brain activity is measured will impact the power in detecting a reliable link to a specific trait. To examine this possibility, we performed a state-space analysis in which tasks from the Human Connectome Project (HCP) were organized into a low-dimensional space based on how they activated different large-scale neural systems. We examined how individuals’ observed brain activity across these different contexts related to their personality. Our analysis found that for multiple personality traits (including Agreeableness, Openness to Experience and Conscientiousness) stronger associations with brain activity emerge in some tasks than others. These data establish that for specific personality traits there are situations in which reliable associations with brain activity can be identified with greater accuracy, highlighting the importance of context-bound views of understanding how brain activity links to trait variation in human behaviour.

**Significance statement:** As a species humans act efficiently in many contexts, however, as individuals our personality makes us more specialised in some situations than others. This “if-then” view of personality is widely accepted in the social sciences but is often overlooked in neuroscience. Here we show adopting a situationally bound view of human traits provides more meaningful descriptions of a brain-trait associations than are possible in traditional brain wide association studies (BWAS) that measure brain activity in a single situation. Our results demonstrate multiple personality traits (including Agreeableness, Openness to Experience and Conscientiousness) show stronger associations with brain activity in some tasks than others, explaining why studies focusing on changes in brain activity at rest can lead to weak or contradictory results.

## Introduction

Adaptive behaviour depends on efficiently meeting the demands imposed by specific environmental conditions, and humans function successfully in a wide range of situations. For example, situations can vary on the need for sustained attention (1), skilled performance acquired through learning (2), or on our knowledge of the world (3). In any specific situation, therefore, optimal performance corresponds to a specific balance of input from different cognitive systems. Consistent with this perspective, contemporary work in psychology has established that how individuals respond to environmental demands provides a useful way to understand trait variation within our species (4). For example, personality dimensions can be conceptualised as “if-then” rules where a given trait is most likely to lead to a type of behaviour when the individual is in a situation with a specific set of features (5).

Although this context-dependent view of human behaviour has made important contributions to the social sciences (6) it has played a less important role in neuroscience (7). For example, studies that link brain activity to traits, often referred to as Brain Wide Association Studies (BWAS), focus on differences in brain activity that emerge during tasks (REF) or often at rest. However, the BWAS paradigm has recently become the subject of controversy due to concerns that without sample sizes in excess of several thousand individuals, the results may be prone to false positives (i.e. Type I error (8), although see (9) for an alternative perspective). Our study set out to explore whether BWAS focusing on an “if-then” view of personality provides an alternative, more useful, way of estimating the brain basis of different human traits.

## Results

In order to determine how brain responses under different situations relate to personality traits, we leveraged the task and resting state functional Magnetic Resonance Imaging (fMRI) data and the self-reported personality measures from the Human Connectome Project (HCP, 10). As descriptions of personality traits we focused on the so called Big 5 personality traits (11) since these traits are replicable (12) and show well described links to real world behaviour (13), (14). In order to compare how different personality traits vary with brain activity across multiple task situations we constructed a state space using the first three dimensions of brain variation from a previous decomposition of the resting state data (15). These dimensions of brain variation, often referred to as “gradients” (16) describe functional differences between activity in different brain systems. We focused on the first three dimensions, which correspond to differences between primary and association cortex (Dimension 1, D1), visual and somato-motor cortex (Dimension 2, D2) and variation between the two large scale systems embedded within association cortex, namely the default mode network, (DMN) and fronto-parietal networks (FPN) (Dimension 3, D3). Note in our study we only use resting-state data to describe brain organisation (e.g. Glasser et al. (17)), and so does not incur the hypothesised problems in using this method for ascertaining brain-trait associations(8). We used this ‘state space’ to organise the macro-scale patterns of each individual’s brain activity in the seven tasks (13 conditions) measured in the HCP by correlating each spatial map for each task condition (contrasted with the implicit baseline of each condition) with each of the dimensions of brain variation (see Methods). This process yielded a set of x, y, z co-ordinates for describing the observed brain activity for each individual in each task context (Figure 1. We also calculated the pair-wise similarities in the whole brain maps; see Supporting Information, SI Figure 2). Our analytic approach, uses no group averaging and preserves the unique functional topography of each individual as this is argued to be important for accurately describing brain organisation ^17, 18^. At the same time, our state-space approach allows trait-related variation in brain activity, contextual differences in brain activity, and their interaction, to be differentiated along one or more of the dimensions of brain variation focused on in our study (See (20–22) for prior demonstrations of this approach). One advantage of our state-space method is that it provides a simple low dimensional manifold in which the impact of traits and situations can both be assessed. In other words, it provides a simple way to assess the possibility that brain-trait relationships are situationally dependent in way that would be analytically complex using more concrete regional approaches to understanding brain function.

**Figure 1.**
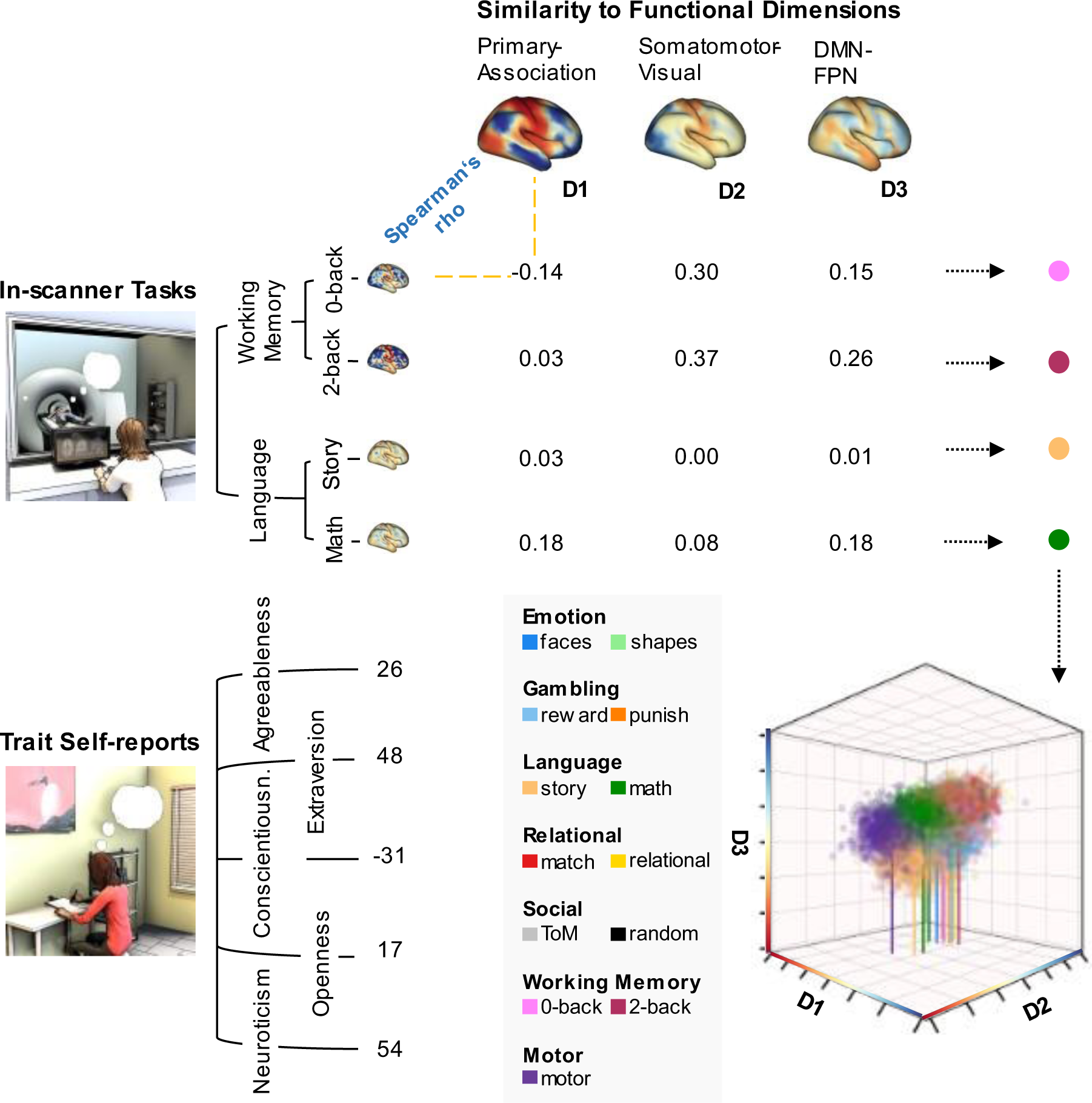
Creation of a state space to understand how the neural correlates of personality traits are differentially expressed across task situations. To simultaneously map neural activity across individuals and situations we utilised a state-space approach (20–22) in which we calculated the correlation between the whole brain map of an individual’s brain activity in a specific task condition (contrasted with the respective baseline) with each of the three dimensions of brain variation generated by the decomposition of brain activity at rest (15). This results in a series of values which can be considered to be co-ordinates in a 3-dimensional space upon which we can conduct inferential statistics to understand how neural activity changes across situations and individuals and how these two influences on brain activity interact. (See SI Figure 1 for the distribution of all individuals’ maps within the space and SI Figure 2 for the group average of each task condition within the state space).

**Figure 2.**
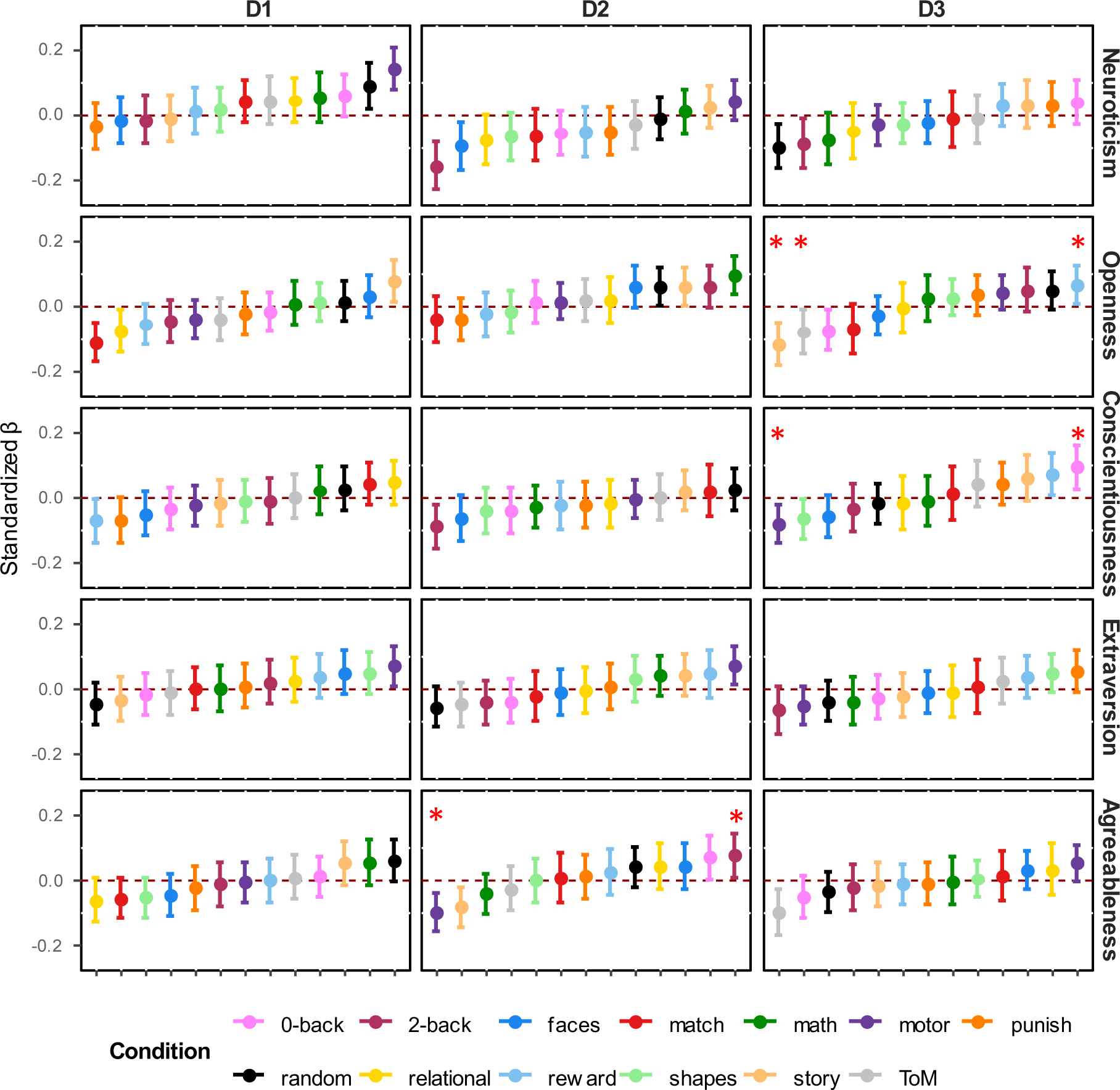
Associations between personality traits and state space location across all dimensions and task conditions. In this figure, each point reflects the estimate of the association between a trait of the “Big 5” and a single dimension of brain variation, under a single task condition, controlling for all other variables in the model. Error bars indicate the 95% Confidence Intervals around this estimate. For ease of interpretation, tasks are ordered from the most negative to the most positive. Significant interaction effects between conditions and traits are marked with asterisk (controlling for multiple comparisons, p < 0.00004). D1 reflects the dissociation from primary cortex (negative) to association cortex (positive). D2 reflects the dissociation between somato-motor (negative) and visual cortex (positive). D3 reflects the dissociation between the default mode network (negative) and fronto-parietal network (positive).

We used linear mixed models, as used the lmerTest package (23) in R (24), to perform inferential statistics on the co-ordinates of each brain map to understand whether they supported a situationally bound account of how brain activity maps onto dimensions of personality (see Methods for details of the models). We performed these analyses once for each dimension of brain variation, and in each analysis modelled (i) the main effect of task conditions, (ii) the main effect of each dimension of personality (Neuroticism, Openness to Experience, Conscientiousness Extraversion and Agreeableness) and (iii) the interactions between each trait dimension and task condition. Subject ID and family ID were modelled as random effects, and age, gender, and mean framewise displacement were added as covariates of no interest. We included family ID as a random effect to control for the fact that the HCP data set includes individuals who are biological siblings.

In these analyses, a main effect of condition indicates a difference between the location of task conditions on the dimension of brain variation of interest. A main effect of a personality trait indicates a similar association between that trait and brain activity across each task context. Finally, a trait-by-condition interaction indicates that the association between trait and brain activity follows an “if-then” rule, because the strength and/or direction of the association between trait and brain activity is variable across the task conditions sampled in our study. To control for family-wise error in these analyses we controlled for the 78 pairwise comparisons between tasks, the five traits that make up the Big 5 and the three dimensions of brain variation (78 * 5 * 3 = 1170). Using the Bonferroni correction method this led to an alpha value of 0.05/1170 = 0.00004.

In each of these three models we identified a significant main effect of task condition (D1: F(_12, 11964_) = 5.92, p<0.00001; D2: F(_12, 972_) = 51.05, p<0.00001; D3: F(_12, 11992_) = 10.83, p<0.00001). This indicates that brain activity recorded in the specific task conditions measured in the HCP significantly varies along each of the three dimensions of brain variation that make up our state space (see SI Table 2 for complete results and SI Figure 2 for mean locations of all conditions in the state space), establishing that across the set of tasks included in the HCP, there was significant variation in the average balance of different neural systems engaged during task completion.

In addition, for two out of three dimensions of brain variation studied we identified at least one example where the association between a trait and brain activity was explained by an “if-then” relationship (Figure 3). For the second dimension of brain variation (D2, differentiation between visual and somato-motor cortex), a significant interaction was observed for Agreeableness (F(_12, 11958_) = 3.82, p< 0.00001). The condition with the most positive association (i.e., towards visual cortex) was the “2-back” working memory (t = 2.17) while the task with the strongest negative association (towards somato-motor cortex) was “motor” (t = −3.35).

**Figure 3.**
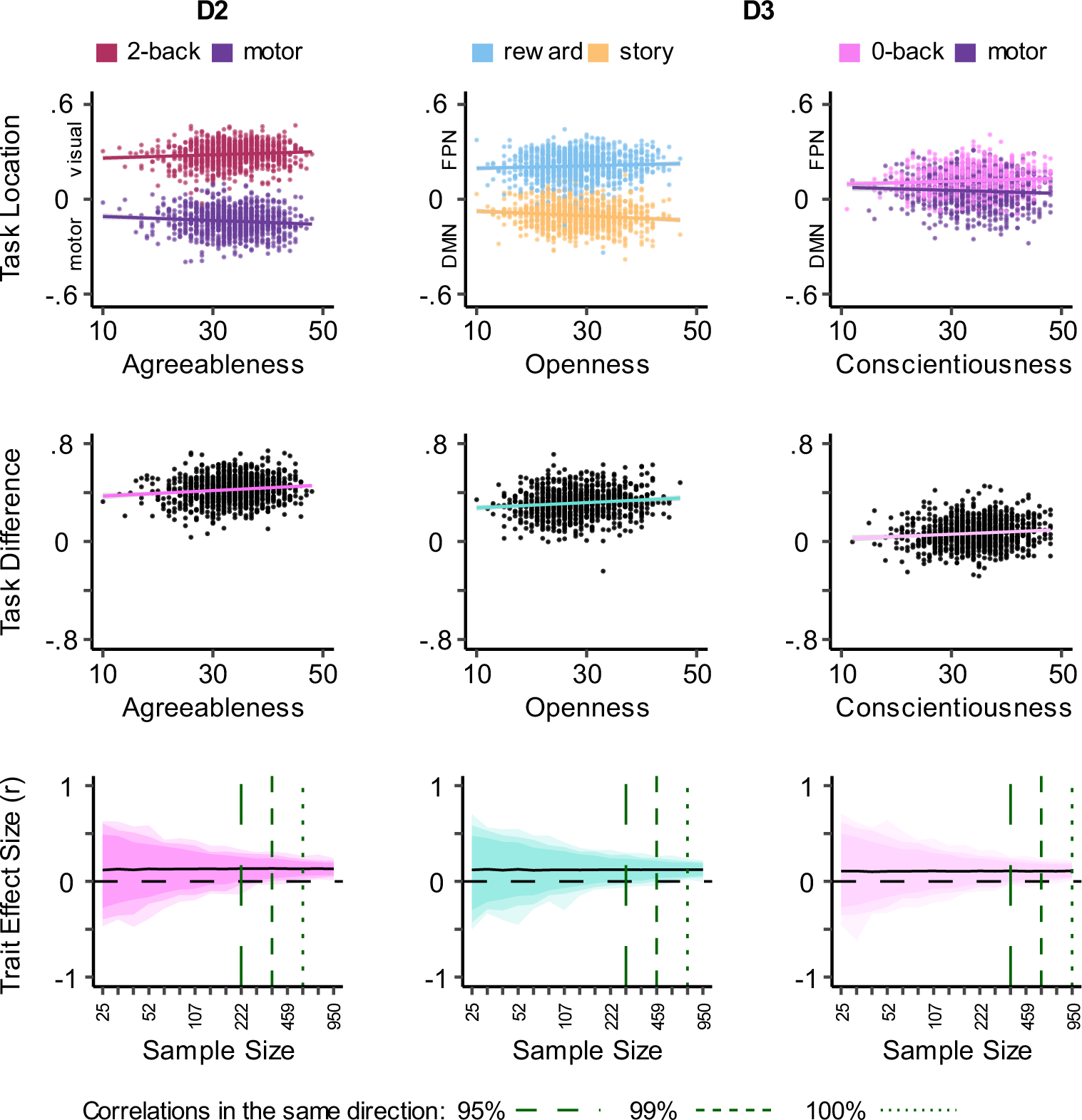
Trait associations with respect to variation between visual and sensorimotor systems (D2) and default mode and fronto-parietal networks (D3). This figure illustrates the strongest relationship between brain activity and trait for the three personality dimensions for which significant interactions were identified: Agreeableness, Openness to Experience and Conscientiousness. Scatter plots in the top row show the relationships between two conditions and the specific trait along the dimension of brain variation of interest. In these plots, x axis shows the trait score and y axis shows the location of a specific task condition on the dimension of brain variation of interest. Each point represents one individual. Scatter plots in the middle row show the correlation between the trait, and the divergence of two task-condition maps shown above on the dimension of brain variation interest. In these plots, y axis shows the pair-wise difference (e.g. 2back - motor) in the location of the two task maps. Plots in the bottom row summarize the results of a bootstrapping analysis showing the distribution of the same correlations as a function of sample size. In these plots the shaded regions show the distribution of 100%, 99%, and 95% of the effects (Pearson’s R) derived from the bootstrapping and vertical dashed lines indicate the sample sizes required to consistently find effects in the same direction within the 95% and 99% confidence intervals, and in the whole range (100%).

For D3 (the dissociation between the default mode network and the fronto-parietal network) we found significant interactions for both Openness to Experience (F(_12, 11979_) = 4.40, p< 0.00001) and Conscientiousness (F(_12, 11984_) = 3.57, p< 0.00003). For Openness to Experience the task with the most negative association (i.e., towards the default mode network) was “story” (t = −3.60), while the most positive was the “reward” condition in the gambling task (t = 2.12). For Conscientiousness, the task with the most negative association was with the “motor” task (t = −2.78) and the “0-back” working memory condition had the strongest positive association (t = 2.66).

It is important to note that in all three models none of the main effects of traits passed our correction for family-wise error; the strongest association was in D2 for Neuroticism (p = .033 uncorrected, complete results of all linear models can be found under Supporting Information Tables 3-9). Together, therefore, our analysis indicates only weak support for the hypothesis that traits will show a general association with brain activity across tasks, and substantial support for the view that these associations are modulated by the task context. Overall, therefore, our data is consistent with the view that traits lead to situationally specific changes in brain activity and inconsistent with the implicit assumptions behind many BWAS that attempt to link traits to a single condition.

Next, we examined the sample size needed to infer associations between traits and brain activity across situations in our analysis. One criticism of BWAS is that the magnitude of associations between activity within regions or sets of regions and traits are often higher with smaller numbers of participants and decline with larger sample sizes: a pattern that is indicative of false positives with underpowered designs (8). For each of the significant interaction effects, therefore, we repeatedly sampled individuals from our population to create samples-sizes ranging from 25 to 950 in 16 log-spaced steps, creating 1000 examples of each sample size, and examined how these relationships changed with increasing power (bottom row, Figures 3). It can be seen that with smaller sample sizes the task-trait relationships begin to stabilize, i.e., generate > 95% estimates that all have the same direction, for samples that are between 222 and 459. Relative to those observed in BWAS that focus on brain activity at rest, these estimates tend to stabilize with equivalent, if not smaller, samples. Lastly, as the full HCP dataset contains pairs of individuals who are siblings, we repeated the reproducibility analysis, generating bootstrapped samples which only contained singletons in each resampling iteration, yielding broadly similar results (See SI Figure 4).

## Discussion

Using a state-space created from dimensions of brain variation observed at rest, we established that different tasks employed in the HCP vary in the whole brain patterns of brain activity they engender, confirming that these situations provoke different challenges to the brain. Consistent with psychological models of cognition and behaviour that emphasise trait variation as a set of “if-then” rules, different tasks systematically varied in their utility to capture the brain activity associated with different traits. Openness to Experience, was most strongly associated with increased activity within the default mode network during the “story” task condition and least in the “reward” condition. Agreeableness was linked to relatively greater activity in somato-motor cortex in the “motor” task and with relatively greater visual activity during “2-back” working memory. In contrast, Conscientiousness was linked to greater engagement of the fronto-parietal system than the default mode network during the “0-back” working memory than the “motor” task.

Together these data illustrate that associations between brain activity and trait variation cannot be mapped equally in a single situation. Consistent with psychological perspectives that traits can be conceived of as stable responses to specific environmental challenges (“if-then” rules) our analysis establishes that brain activity correlates of traits vary substantially across situations. This suggests that regardless of sample size (8), or analytic approach (9), BWAS will likely have greater success in detecting accurate correlates of specific traits by tailoring the situations in which brain activity is measured to challenge the brain in an appropriate manner. Our study, therefore, establishes that one reason why BWAS yield inconsistent results is that human traits are inextricably linked to how the individual responds to specific situations. While it is important to employ well powered designs, and better analytic approaches, BWAS will become more useful following the development of better theoretical models which include the situations in which traits influence behaviour and appropriate paradigms to capture them e.g. (25). We note that a trend for estimating trait associations in appropriate contexts is also emerging in population studies of genetics (26).

At the same time as illustrating that trait associations with brain activity are situationally bound, our approach highlights important new avenues of inquiry for understanding how human variation is linked to patterns of brain activity. For example, what are the best situations for mapping the neural correlates of specific traits, and how many situations are needed to efficiently map the bulk of human traits? We utilised the HCP task data because it is the largest existing dataset to sample a wide range of tasks along with trait measures. However, the selection of tasks for the HCP was not designed to test “if-then” relationships between personality and brain activity. It is likely, therefore, that there are better batteries of tasks with which to distinguish the neural correlates of different traits. Fortunately, techniques such as neuroadaptive Bayesian optimisation, which uses machine learning to efficiently identify correlations between brain activity and behaviour, can be used to tailor situations to map traits to brain activity efficiently (27). Iteratively, this process will enable the situations in which we measure brain activity to be optimized, and in turn improve the efficiency with which the neural correlates of different features of human behaviour can be understood in BWAS.

Our analyses also highlight specific analytic questions that will need to be resolved to properly determine how the correlations between neural activity and traits vary across situations. For example, our study used a state space based on patterns of brain variation at rest as a low dimensional space to preserve individual topography while simultaneously modelling task and individual differences in brain function. It has been shown that using broad-scale activations as a measure of brain function offers more statistical power over regional effects, with a relatively moderate decrease in specificity (25). We used functional gradients because they are a convenient tool for organising brain-wide activity (20–22). However, there are likely better ways to characterise the dimensions of brain variation and organisation in order to perform state-space analyses like the one we report here. For example, contemporary work in neuroscience has identified dimensions of brain variation that combine information related to brain structure with functional behaviour (e.g. (28)). Fortunately, machine learning can be used to perform multiverse analyses that optimise how different features of brain organisation can be best analysed to maximise their links with phenotypes (29). It is likely that optimising the dimensions of brain variation will generate state spaces that allow BWAS to estimate trait related patterns of brain activity in a more effective manner. Finally, it is important to note that our study focused on trait descriptions of human behaviour that have well established features (i.e., the “Big 5”). Our analytic choice was motivated by the idea that BWAS should focus on traits that have real world significance (30). In this context, the “Big 5” have well established reliability (12) and are predictive of behaviour in real-world situations including academia (13) and the workplace (14) and psychopathology(31). However, it remains to be seen whether the same tasks which establish the situationally specific neural correlates of the “Big 5” can also discriminate the brain mechanisms which impact mental health, or physical illness, both of which are probably the most important outcomes from BWAS studies (30). In the future, therefore, it is important to take seriously the goal of understanding how to tailor task conditions for acquiring brain activity that have better capacity to discriminate phenotypes linked to health, wellbeing, productivity and disease.

In conclusion, our state space analysis of the tasks in the HCP suggest that brain-trait associations are inextricably linked to the context in which brain activity is measured and this observation leads to two concrete suggestions for future studies. First, when examining specific brain-trait relationships it would be helpful to consider the most appropriate situations in which these association will emerge as out study shows that this intimately related to the sample size needed for these associations to stabilise. Second, if in the future it is deemed important to generate large data sets similar to the HCP (10) or the Adolescent Brain Cognitive Development (ABCD, 32) project, then it is likely that the statistical power for detecting robust brain correlates for a range of different traits can be derived from a combination of both the amount of time spent acquiring data in a specific situation (enabling stable measurement) and the range of different cognitive and emotional features that data acquisition encompasses (which enables the testing battery to discriminate multiple different traits).

## Materials and Methods

### Data

We used task and resting state fMRI, and self-reported questionnaire data from the human Connectome Project (10) 1200 subjects release. The HCP dataset (N= 1206) includes multimodal MRI, behavioural, genetic, physiological and demographic data from adult twins and their non-twin siblings between 22-37 years of age. From these, our analysis made use of the minimally preprocessed (2mm smoothing) (33) task-fMRI maps and NEO-FFI personality measures (11) and summaries of the group-averaged functional connectivity matrix from the 900 subjects release (15). Additionally, we used demographic information of subjects (age in years, gender), and head-movement parameters of each task-fMRI session as covariates. The final sample size of all HCP subjects with preprocessed task-fMRI data available for download is 1088 (590 women, mean age = 29.52 ± 3.59 years; 498 men, mean age = 27.92 ± 3.61 years). SI Table 1 shows the number of subjects available in each task condition.

### Neural State Space

To create a neural state space, which describes maximal functional covariation of different neural systems, we used previously established low dimensional summaries of the group-level whole-brain functional connectivity matrix (15). These dimensions of brain variation, often referred to as “gradients”, describe functional differences between brain systems. In our analysis, we created a three-dimensional “state space” from the first three gradients which correspond to differences between (i) primary and association cortex, (ii) visual and sensorimotor cortex and (iii) variation between the two large scale systems embedded within association cortex (default mode network, DMN, and fronto-parietal networks, FPN).

To project each individual’s brain activity across different contexts into the state space, we calculated spearman rank correlation between each of the three gradient maps and each individual’s un-thresholded z-map from each task condition in grayordinate space. (Figure 1). Only the main contrasts (each condition against the respective implicit baseline) were used for this purpose, resulting in 13 maps for each individual, namely, Motor: 1)average of all movements; Emotion: 1) faces 2) shapes; Language: 1) maths, 2) story; Social: 1) random interactions, 2) theory of mind (ToM); Working Memory: 1) 0-back, 2) 2-back; Gambling: 1) reward, 2) punish; and Relational: 1) match 2) relational). The correlation coefficient of each map with the three gradients served as the location of that map along the respective dimension of brain variation in the state space (Figure 1).

### Linear Mixed Models

To understand and quantify how locations of tasks in the state space varied with dimensions of personality, we performed regression using linear mixed models once for each dimension of brain variation as the outcome variable, and the task context, each dimension of personality (Neuroticism, Openness to Experience, Conscientiousness, Extraversion, and Agreeableness) and the interactions between each dimension and each task condition as predictors. Subject ID and family ID were added as random effects, and age, gender, and mean framewise displacement were used as covariates of no interest.

Example model for one dimension of variation:

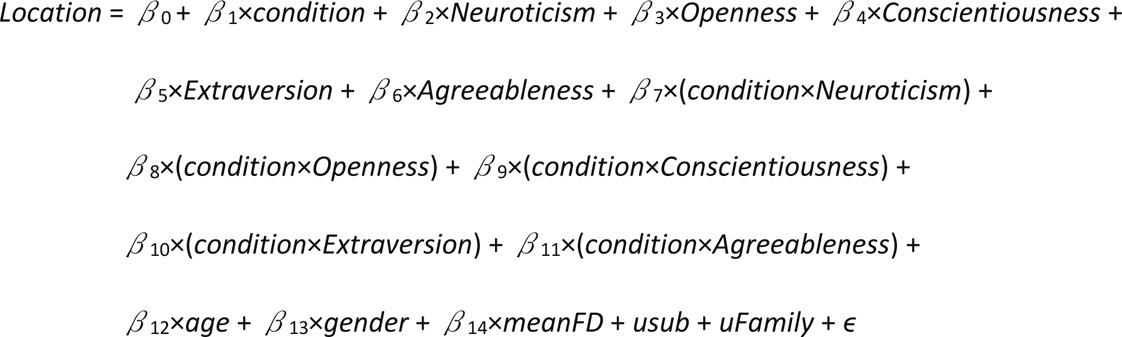

In these analyses, a main effect of task indicates a difference between the location of tasks on the dimension of interest. A main effect of personality trait indicates a similar association with brain activity with a trait across each task context. Finally, a trait-by-task interaction indicates that associations between traits and brain activity varied in their strength, direction, or both across different tasks. To control for family wise error in these analyses we controlled for the 78 pairwise comparisons between tasks, the five traits which make up the “Big 5” and the three dimensions of brain variation (78 * 5 * 3 = 1170). Using the Bonferroni correction method this led to an alpha value of 0.05/1170 = 0.00004. To illustrate the change in trait-brain associations depending on context, we followed up each significant trait*condition interaction, by comparing the strongest positive and negative associations of the respective trait and with task locations along dimensions of brain variation (Figures 3). Linear models were fitted using the lmerTest(34) package in R(24). We used the emmeans(35) package to derive the slope for each trait in the model at each level of the factor “condition”, resulting in an estimate for the association of each combination of trait and task condition and state-space location shown in Figure 2.

### Reproducibility of context-specific brain-wide associations

To examine the distribution of effects found in our analysis as a function of sample size, and to estimate the sample size required to reliably identify such effects, we calculated the bootstrapped (with 1000 iterations) bivariate correlation estimates and confidence intervals for all significant task-brain interactions (Figures 3). Following Marek and colleagues (8) we focused on the strongest associations identified in our initial analyses. For each of these effects, we created 16 logarithmically spaced samples sizes from 25 to 950 subjects, by resampling subjects with replacement 1000 times at each sample size. Similar to the follow-up analysis for task*trait interactions in the original sample, in each resampled dataset, we calculated the bivariate correlation between trait scores and the distance between two brain maps that show the strongest diverging associations with that trait along a dimension. (e.g., correlation between Openness and (D3 location of reward – D3 location of story). Figures 3 shows the distribution of these bootstrapped estimates with increasing sample sizes and indicates the 95% and 99% confidence intervals as well as full range of effect sizes derived from bootstrapping.

Finally, given that the HCP dataset is made up of sibling pairs and groups, to avoid inflated estimates resulting from resampling of closely related individuals, we repeated the bootstrapping analysis in a smaller subsample (n=442) of “singletons” where, in each iteration, no more than one member of each family could be included at a time. For this analysis, we used 13 log-spaced sample sizes between 25 and 442. Results of this analysis are shown in SI Figure 4.

## Data availability

All data included in the present analyses were acquired with informed consent and are available at https://db.humanconnectome.org/.

## Code Availability

All code used for analysis and visualization can be found at https://github.com/samyogita-hardikar/hcp-task-trait

## Corresponding author

Correspondence and requests for materials should be sent to Samyogita Hardikar or Jonathan Smallwood.

