## Supplemental Information for "Personality traits vary in their association with brain activity across situations"

**Supplemental Information (SI)**

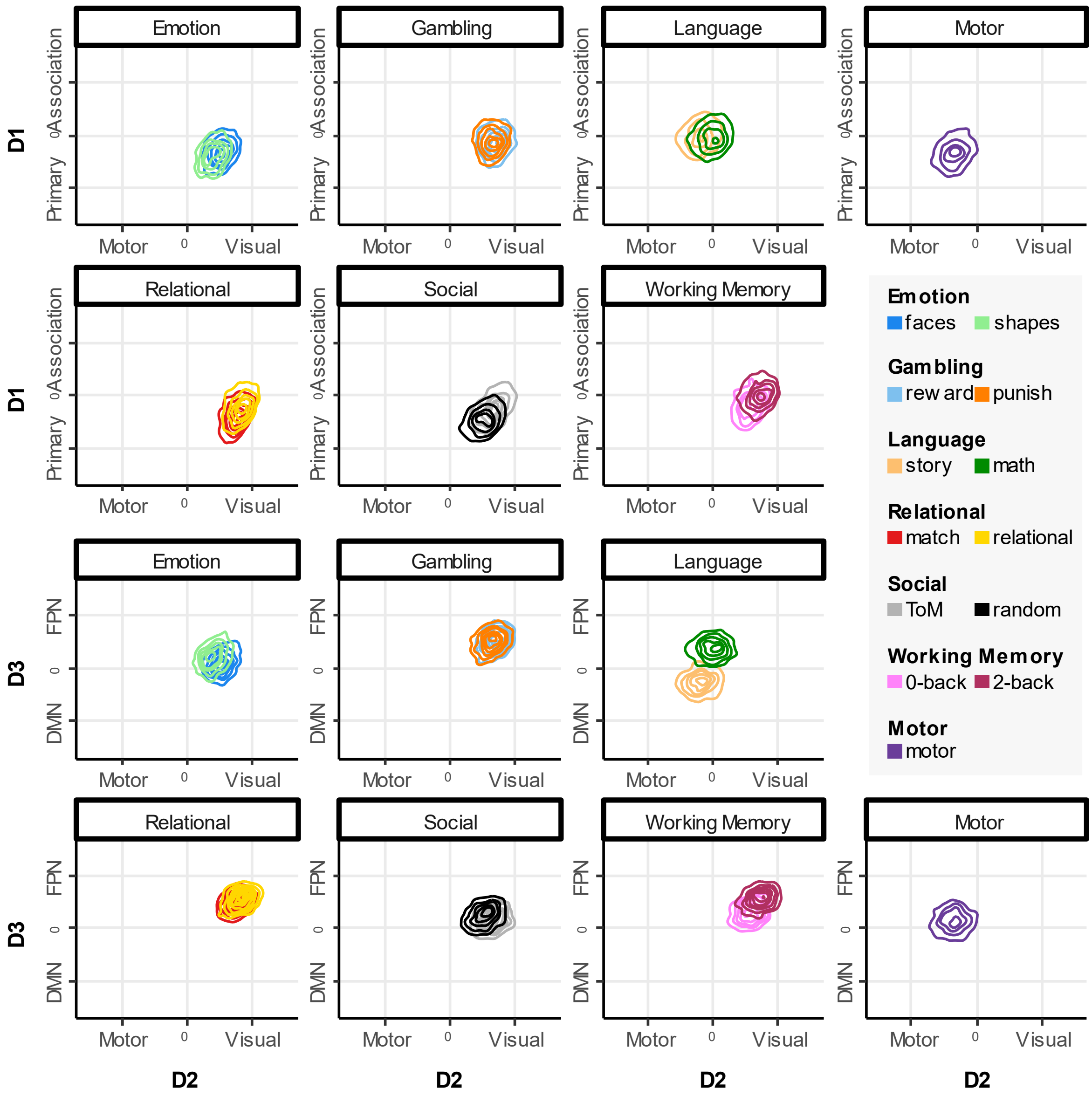


**SI Figure 1. Contour plots illustrating the distribution of all individuals' data for each task condition within the state space.** Each contour plots shows the distribution of brain maps from the HCP data along two dimensions of brain variation. The location along each dimension is determined by the correlation between the task-maps of each individual and the respective functional connectivity gradient identified at rest (see Figure 1). Conditions from the same task are plotted in the same panel.

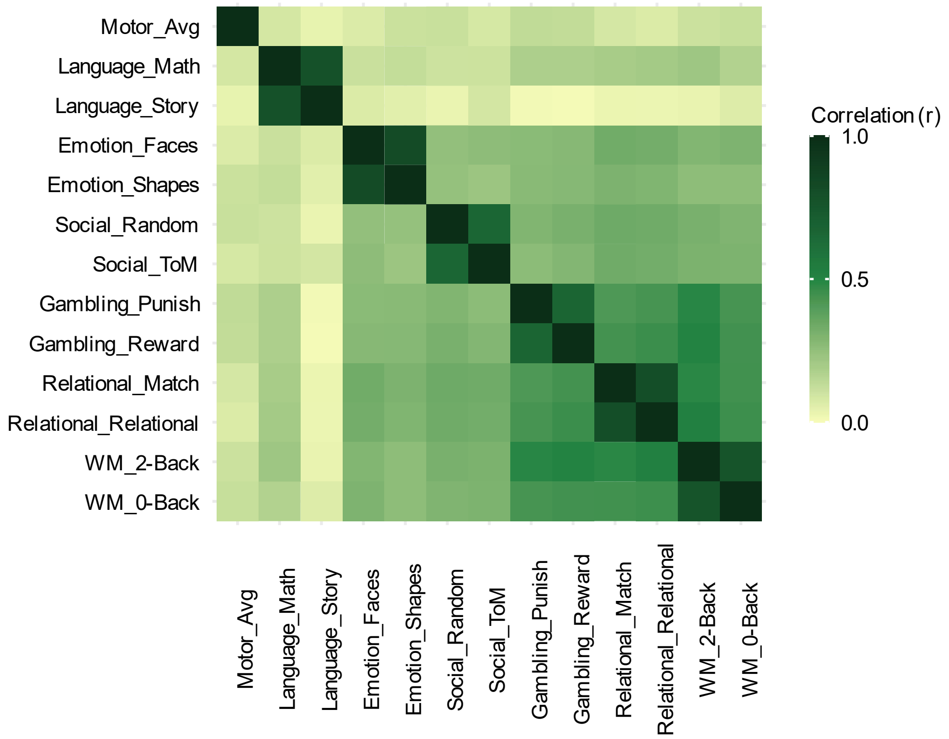


**SI Figure 2. Heatmap showing the group-averaged spearman rank correlations between brain-wide activity patterns across different task conditions.** The group level correlation matrix was derived by first calculating all pair-wise spearman rank correlations between task maps for each individual, and then calculating the mean score across all individuals for each pair at the group level.

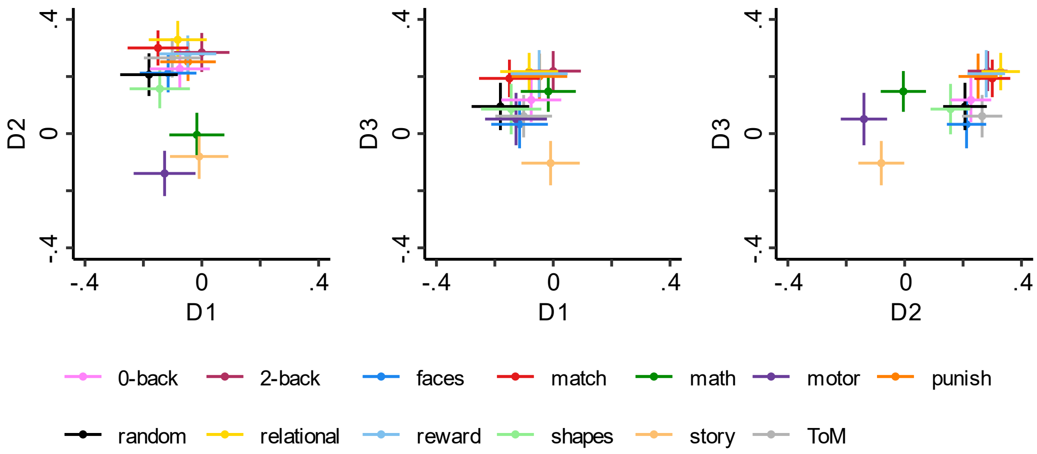


**SI Figure 3. Location of whole brain maps in the state space, shown as group means and standard deviations for each task condition.**

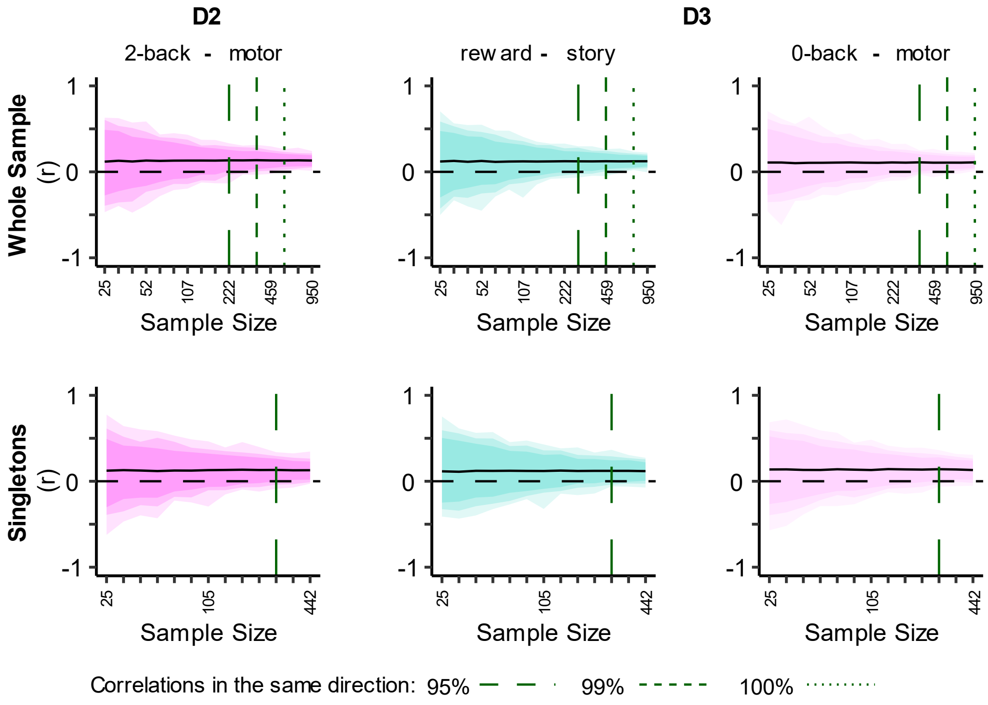


**SI Figure 4. Replication of strongest trait-brain associations in a "singletons only" subsample of HCP.** These plots summarize the results of a bootstrapping analysis showing the distribution of the same correlations as a function of sample size. Vertical lines indicate the sample sizes required to consistently find effects in the same direction within the 95%, and 99% confidence intervals, and in the whole range (100%).

**SI Table 1.** Number of subjects with minimally pre-processed contrast-maps available for each task in the HCP task battery.

| **variable** | **(available) n** |
|---|---:|
| Subject | 1088 |
| Neuroticism | 1083 |
| Openness | 1083 |
| Conscientiousness | 1083 |
| Extraversion | 1083 |
| Agreeableness | 1083 |
| Age | 1088 |
| Gender | 1088 |
| Emotion | 1017 |
| Gambling | 1060 |
| Language | 941 |
| Motor | 1032 |
| Relational | 994 |
| Theory of Mind (ToM) | 1032 |
| Working Memory | 1061 |

**SI Table 2.** Results of Linear Mixed Models showing the associations between personality traits, task context, and dimensions of brain variation.

| DV | IV | Sum Sq | Mean Sq | NumDF | DenDF | F value | Pr(>F) |
|---|---|---|---|---|---|---|---|
| D1 | cond | 0.48 | 0.04 | 12.0000 | 11963.65 | 5.92 | 0.000000 |
| | Age | 0.15 | 0.15 | 1.0000 | 1034.21 | 22.05 | 0.000003 |
| | Gender | 0.38 | 0.38 | 1.0000 | 1046.55 | 56.48 | 0.000000 |
| | motion | 0.30 | 0.30 | 1.0000 | 4486.02 | 44.59 | 0.000000 |
| | Neuroticism | 0.02 | 0.02 | 1.0000 | 1031.87 | 2.59 | 0.107586 |
| | Openness | 0.01 | 0.01 | 1.0000 | 1043.52 | 1.37 | 0.241949 |
| | Conscientiousness | 0.00 | 0.00 | 1.0000 | 1042.08 | 0.41 | 0.521734 |
| | Extraversion | 0.00 | 0.00 | 1.0000 | 1035.67 | 0.32 | 0.571320 |
| | Agreeableness | 0.00 | 0.00 | 1.0000 | 1012.51 | 0.13 | 0.716277 |
| | cond:Neuroticism | 0.23 | 0.02 | 12.0000 | 11952.38 | 2.86 | 0.000614 |
| | cond:Openness | 0.27 | 0.02 | 12.0000 | 11951.15 | 3.36 | 0.000066 |
| | cond:Conscientiousness | 0.14 | 0.01 | 12.0000 | 11955.93 | 1.67 | 0.066138 |
| | cond:Extraversion | 0.12 | 0.01 | 12.0000 | 11947.43 | 1.52 | 0.109945 |
| | cond:Agreeableness | 0.18 | 0.02 | 12.0000 | 11949.54 | 2.24 | 0.008204 |
| D2 | cond | 2.24 | 0.19 | 12.0000 | 11972.46 | 51.05 | 0.000000 |
| | Age | 0.00 | 0.00 | 1.0000 | 1026.75 | 1.35 | 0.244854 |
| | Gender | 0.00 | 0.00 | 1.0000 | 1051.29 | 0.67 | 0.411903 |
| | motion | 0.12 | 0.12 | 1.0000 | 3964.27 | 33.09 | 0.000000 |
| | Neuroticism | 0.02 | 0.02 | 1.0000 | 1042.05 | 4.54 | 0.033380 |
| | Openness | 0.01 | 0.01 | 1.0000 | 1041.96 | 1.63 | 0.202610 |
| | Conscientiousness | 0.00 | 0.00 | 1.0000 | 1052.46 | 1.17 | 0.279410 |
| | Extraversion | 0.00 | 0.00 | 1.0000 | 1043.06 | 0.01 | 0.929634 |
| | Agreeableness | 0.00 | 0.00 | 1.0000 | 1027.54 | 0.00 | 0.954479 |
| | cond:Neuroticism | 0.12 | 0.01 | 12.0000 | 11960.58 | 2.77 | 0.000892 |
| | cond:Openness | 0.11 | 0.01 | 12.0000 | 11959.69 | 2.48 | 0.003106 |
| | cond:Conscientiousness | 0.05 | 0.00 | 12.0000 | 11964.77 | 1.20 | 0.276057 |
| | cond:Extraversion | 0.09 | 0.01 | 12.0000 | 11955.57 | 2.16 | 0.011168 |
| | cond:Agreeableness | 0.17 | 0.01 | 12.0000 | 11957.55 | 3.82 | 0.000008 |
| D3 | cond | 0.61 | 0.05 | 12.0000 | 11991.89 | 10.83 | 0.000000 |
| | Age | 0.00 | 0.00 | 1.0000 | 1048.42 | 0.35 | 0.554338 |
| | Gender | 0.09 | 0.09 | 1.0000 | 1061.92 | 19.40 | 0.000012 |
| | motion | 0.65 | 0.65 | 1.0000 | 3459.66 | 138.19 | 0.000000 |
| | Neuroticism | 0.01 | 0.01 | 1.0000 | 1048.19 | 1.38 | 0.240672 |
| | Openness | 0.00 | 0.00 | 1.0000 | 1058.04 | 0.11 | 0.735335 |

| | | | | | | |
|---|---|---|---|---|---|---|
| Conscientiousness | 0.00 | 0.00 | 1.0000 | 1060.99 | 0.00 | 0.996172 |
| Extraversion | 0.00 | 0.00 | 1.0000 | 1051.58 | 0.18 | 0.670017 |
| Agreeableness | 0.00 | 0.00 | 1.0000 | 1029.67 | 0.30 | 0.585759 |
| cond:Neuroticism | 0.12 | 0.01 | 12.0000 | 11979.31 | 2.04 | 0.017536 |
| cond:Openness | 0.25 | 0.02 | 12.0000 | 11978.93 | 4.40 | 0.000001 |
| cond:Conscientiousness | 0.20 | 0.02 | 12.0000 | 11984.30 | 3.57 | 0.000025 |
| cond:Extraversion | 0.10 | 0.01 | 12.0000 | 11974.14 | 1.70 | 0.059102 |
| cond:Agreeableness | 0.10 | 0.01 | 12.0000 | 11976.01 | 1.78 | 0.045507 |

**DV**= Dependent Variable; **IV**= Independent Variable; **cond**=condition

**SI Table 3.** Estimated marginal means of all task contrast maps along state-space dimensions.

| Dimension | condition | emmean | SE | df | lower CL | upper CL |
|---|---|---|---|---|---|---|
| D1 | 0back | -0.074 | 0.0032 | 2053.36 | -0.080 | -0.068 |
| | 2back | 0.002 | 0.0032 | 2053.36 | -0.005 | 0.008 |
| | faces | -0.113 | 0.0033 | 2150.67 | -0.120 | -0.107 |
| | match | -0.149 | 0.0033 | 2216.20 | -0.156 | -0.143 |
| | math | -0.015 | 0.0034 | 2382.98 | -0.022 | -0.009 |
| | motor | -0.128 | 0.0033 | 2134.89 | -0.135 | -0.122 |
| | punish | -0.045 | 0.0032 | 2060.85 | -0.051 | -0.038 |
| | random | -0.180 | 0.0033 | 2117.63 | -0.186 | -0.173 |
| | relational | -0.082 | 0.0033 | 2216.20 | -0.088 | -0.075 |
| | reward | -0.046 | 0.0032 | 2060.85 | -0.052 | -0.039 |
| | shapes | -0.142 | 0.0033 | 2150.67 | -0.148 | -0.135 |
| | story | -0.008 | 0.0034 | 2382.98 | -0.014 | -0.001 |
| | ToM | -0.100 | 0.0033 | 2117.63 | -0.106 | -0.094 |
| D2 | 0back | 0.227 | 0.0023 | 2533.41 | 0.222 | 0.231 |
| | 2back | 0.284 | 0.0023 | 2533.41 | 0.280 | 0.289 |
| | faces | 0.211 | 0.0023 | 2655.56 | 0.207 | 0.216 |
| | match | 0.300 | 0.0023 | 2739.28 | 0.295 | 0.304 |
| | math | -0.005 | 0.0024 | 2953.96 | -0.010 | 0.000 |
| | motor | -0.139 | 0.0023 | 2635.63 | -0.144 | -0.135 |
| | punish | 0.251 | 0.0023 | 2542.29 | 0.247 | 0.256 |
| | random | 0.206 | 0.0023 | 2613.97 | 0.201 | 0.210 |
| | relational | 0.328 | 0.0023 | 2739.28 | 0.324 | 0.333 |
| | reward | 0.279 | 0.0023 | 2542.29 | 0.275 | 0.284 |
| | shapes | 0.157 | 0.0023 | 2655.56 | 0.152 | 0.161 |
| | story | -0.080 | 0.0024 | 2953.96 | -0.085 | -0.076 |
| | ToM | 0.264 | 0.0023 | 2613.97 | 0.260 | 0.269 |
| D3 | 0back | 0.117 | 0.0025 | 3108.27 | 0.112 | 0.122 |
| | 2back | 0.218 | 0.0025 | 3108.27 | 0.213 | 0.223 |
| | faces | 0.032 | 0.0026 | 3254.83 | 0.027 | 0.037 |
| | match | 0.194 | 0.0026 | 3355.92 | 0.189 | 0.199 |
| | math | 0.147 | 0.0026 | 3616.68 | 0.142 | 0.153 |
| | motor | 0.053 | 0.0026 | 3229.93 | 0.048 | 0.058 |
| | punish | 0.199 | 0.0025 | 3118.63 | 0.194 | 0.204 |

| | | | | | |
|---|---|---|---|---|---|
| random | 0.094 | 0.0026 | 3204.54 | 0.089 | 0.099 |
| relational | 0.218 | 0.0026 | 3355.92 | 0.213 | 0.223 |
| reward | 0.209 | 0.0025 | 3118.63 | 0.204 | 0.214 |
| shapes | 0.085 | 0.0026 | 3254.83 | 0.080 | 0.090 |
| story | -0.104 | 0.0026 | 3616.68 | -0.109 | -0.098 |
| ToM | 0.060 | 0.0026 | 3204.54 | 0.055 | 0.065 |

**SI Table 4.** Estimated marginal trends (slopes) of personality traits for all task conditions along D1 (primary- association)

| **condition** | **$\beta_{Neuroticism}$** | **SE** | **df** | **$CL_{lower}$** | **$CL_{upper}$** | **t.ratio** | **p.value** |
|---|---|---|---|---|---|---|---|
| 0back | 0.0008 | 0.0005 | 6739.25 | -0.0001 | 0.0017 | 1.731 | 0.08358 |
| 2back | -0.0002 | 0.0005 | 6739.24 | -0.0011 | 0.0007 | -0.419 | 0.67548 |
| faces | -0.0002 | 0.0005 | 6929.58 | -0.0011 | 0.0007 | -0.457 | 0.64775 |
| match | 0.0006 | 0.0005 | 7041.75 | -0.0004 | 0.0015 | 1.208 | 0.22696 |
| math | 0.0007 | 0.0005 | 7415.50 | -0.0003 | 0.0016 | 1.366 | 0.17203 |
| motor | 0.0020 | 0.0005 | 6840.75 | 0.0011 | 0.0030 | 4.322 | 0.00002 |
| punish | -0.0005 | 0.0005 | 6740.19 | -0.0014 | 0.0004 | -1.032 | 0.30188 |
| random | 0.0012 | 0.0005 | 6872.28 | 0.0002 | 0.0021 | 2.479 | 0.01321 |
| relational | 0.0006 | 0.0005 | 7041.75 | -0.0004 | 0.0015 | 1.226 | 0.22031 |
| reward | 0.0001 | 0.0005 | 6740.19 | -0.0008 | 0.0011 | 0.280 | 0.77982 |
| shapes | 0.0002 | 0.0005 | 6929.58 | -0.0007 | 0.0012 | 0.464 | 0.64235 |
| story | -0.0001 | 0.0005 | 7415.50 | -0.0011 | 0.0008 | -0.306 | 0.75964 |
| ToM | 0.0006 | 0.0005 | 6872.28 | -0.0004 | 0.0015 | 1.206 | 0.22787 |
| **condition** | **$\beta_{Openness}$** | **SE** | **df** | **$CL_{lower}$** | **$CL_{upper}$** | **t.ratio** | **p.value** |
| 0back | -0.0003 | 0.0005 | 6615.40 | -0.0013 | 0.0007 | -0.602 | 0.54746 |
| 2back | -0.0007 | 0.0005 | 6615.39 | -0.0017 | 0.0002 | -1.460 | 0.14434 |
| faces | 0.0005 | 0.0005 | 6820.61 | -0.0005 | 0.0015 | 0.932 | 0.35147 |
| match | -0.0019 | 0.0005 | 6943.89 | -0.0029 | -0.0008 | -3.623 | 0.00029 |
| math | 0.0001 | 0.0005 | 7361.77 | -0.0009 | 0.0012 | 0.244 | 0.80740 |
| motor | -0.0007 | 0.0005 | 6746.85 | -0.0017 | 0.0003 | -1.413 | 0.15776 |
| punish | -0.0003 | 0.0005 | 6549.14 | -0.0013 | 0.0006 | -0.672 | 0.50138 |
| random | 0.0002 | 0.0005 | 6748.70 | -0.0008 | 0.0012 | 0.418 | 0.67572 |
| relational | -0.0012 | 0.0005 | 6943.89 | -0.0022 | -0.0002 | -2.425 | 0.01534 |
| reward | -0.0009 | 0.0005 | 6549.14 | -0.0019 | 0.0001 | -1.768 | 0.07705 |
| shapes | 0.0002 | 0.0005 | 6820.61 | -0.0008 | 0.0012 | 0.311 | 0.75557 |
| story | 0.0012 | 0.0005 | 7361.77 | 0.0002 | 0.0023 | 2.360 | 0.01830 |
| ToM | -0.0006 | 0.0005 | 6748.70 | -0.0016 | 0.0003 | -1.282 | 0.19999 |
| **condition** | **$\beta_{Conscientiousness}$** | **SE** | **df** | **$CL_{lower}$** | **$CL_{upper}$** | **t.ratio** | **p.value** |
| 0back | -0.0006 | 0.0006 | 6917.68 | -0.0017 | 0.0005 | -1.049 | 0.29401 |
| 2back | -0.0002 | 0.0006 | 6917.67 | -0.0013 | 0.0009 | -0.307 | 0.75855 |
| faces | -0.0008 | 0.0006 | 7103.86 | -0.0020 | 0.0003 | -1.461 | 0.14399 |
| match | 0.0007 | 0.0006 | 7247.03 | -0.0004 | 0.0018 | 1.185 | 0.23615 |
| math | 0.0003 | 0.0006 | 7625.04 | -0.0008 | 0.0015 | 0.549 | 0.58297 |
| motor | -0.0005 | 0.0006 | 7054.14 | -0.0016 | 0.0007 | -0.797 | 0.42538 |

| | | | | | | | |
|---|---|---|---|---|---|---|---|
| punish | -0.0011 | 0.0006 | 6840.30 | -0.0022 | 0.0000 | -1.997 | 0.04588 |
| random | 0.0004 | 0.0006 | 7112.79 | -0.0007 | 0.0015 | 0.685 | 0.49334 |
| relational | 0.0007 | 0.0006 | 7247.03 | -0.0004 | 0.0019 | 1.294 | 0.19569 |
| reward | -0.0012 | 0.0006 | 6840.30 | -0.0023 | -0.0001 | -2.107 | 0.03516 |
| shapes | -0.0003 | 0.0006 | 7103.86 | -0.0014 | 0.0009 | -0.439 | 0.66033 |
| story | -0.0003 | 0.0006 | 7625.04 | -0.0014 | 0.0009 | -0.481 | 0.63086 |
| ToM | 0.0000 | 0.0006 | 7112.79 | -0.0011 | 0.0011 | 0.042 | 0.96611 |
| **condition** | **$\beta_{Extraversion}$** | **SE** | **df** | **$CL_{lower}$** | **$CL_{upper}$** | **t.ratio** | **p.value** |
| 0back | -0.0003 | 0.0006 | 6898.90 | -0.0014 | 0.0008 | -0.538 | 0.59030 |
| 2back | 0.0003 | 0.0006 | 6898.89 | -0.0008 | 0.0014 | 0.561 | 0.57472 |
| faces | 0.0008 | 0.0006 | 6993.66 | -0.0003 | 0.0019 | 1.380 | 0.16762 |
| match | 0.0000 | 0.0006 | 7178.98 | -0.0011 | 0.0011 | -0.046 | 0.96293 |
| math | 0.0000 | 0.0006 | 7455.71 | -0.0011 | 0.0011 | 0.000 | 0.99963 |
| motor | 0.0012 | 0.0006 | 7066.46 | 0.0001 | 0.0023 | 2.201 | 0.02778 |
| punish | 0.0001 | 0.0006 | 6873.23 | -0.0010 | 0.0012 | 0.181 | 0.85642 |
| random | -0.0008 | 0.0006 | 6959.69 | -0.0019 | 0.0003 | -1.421 | 0.15527 |
| relational | 0.0004 | 0.0006 | 7178.98 | -0.0007 | 0.0015 | 0.757 | 0.44892 |
| reward | 0.0006 | 0.0006 | 6873.23 | -0.0005 | 0.0017 | 1.104 | 0.26967 |
| shapes | 0.0008 | 0.0006 | 6993.66 | -0.0003 | 0.0019 | 1.509 | 0.13142 |
| story | -0.0006 | 0.0006 | 7455.71 | -0.0017 | 0.0005 | -1.011 | 0.31212 |
| ToM | -0.0002 | 0.0006 | 6959.69 | -0.0013 | 0.0009 | -0.421 | 0.67371 |
| **condition** | **$\beta_{Agreeableness}$** | **SE** | **df** | **$CL_{lower}$** | **$CL_{upper}$** | **t.ratio** | **p.value** |
| 0back | 0.0002 | 0.0006 | 6534.58 | -0.0009 | 0.0013 | 0.303 | 0.76220 |
| 2back | -0.0002 | 0.0006 | 6534.57 | -0.0013 | 0.0009 | -0.369 | 0.71241 |
| faces | -0.0008 | 0.0006 | 6658.90 | -0.0019 | 0.0003 | -1.357 | 0.17494 |
| match | -0.0010 | 0.0006 | 6774.67 | -0.0021 | 0.0001 | -1.814 | 0.06973 |
| math | 0.0009 | 0.0006 | 7336.52 | -0.0003 | 0.0020 | 1.444 | 0.14888 |
| motor | -0.0001 | 0.0006 | 6683.68 | -0.0012 | 0.0010 | -0.220 | 0.82589 |
| punish | -0.0004 | 0.0006 | 6544.37 | -0.0015 | 0.0007 | -0.748 | 0.45459 |
| random | 0.0010 | 0.0006 | 6585.19 | -0.0001 | 0.0021 | 1.809 | 0.07056 |
| relational | -0.0011 | 0.0006 | 6774.67 | -0.0022 | 0.0001 | -1.861 | 0.06278 |
| reward | 0.0000 | 0.0006 | 6544.37 | -0.0011 | 0.0011 | -0.076 | 0.93924 |
| shapes | -0.0010 | 0.0006 | 6658.90 | -0.0021 | 0.0001 | -1.733 | 0.08309 |
| story | 0.0009 | 0.0006 | 7336.52 | -0.0003 | 0.0020 | 1.512 | 0.13070 |
| ToM | 0.0001 | 0.0006 | 6585.19 | -0.0010 | 0.0013 | 0.263 | 0.79247 |

**SI Table 5.** Estimated marginal trends (slopes) of personality traits for all task conditions along D2 (motor- visual)

| condition | $\beta_{Neuroticism}$ | SE | df | $CL_{lower}$ | $CL_{upper}$ | t.ratio | p.value |
|---|---|---|---|---|---|---|---|
| 0back | -0.0005 | 0.0003 | 7616.59 | -0.0012 | 0.0001 | -1.570 | 0.11656 |
| 2back | -0.0014 | 0.0003 | 7616.58 | -0.0021 | -0.0008 | -4.291 | 0.00002 |
| faces | -0.0009 | 0.0003 | 7807.57 | -0.0015 | -0.0002 | -2.546 | 0.01092 |
| match | -0.0005 | 0.0003 | 7923.02 | -0.0012 | 0.0002 | -1.503 | 0.13281 |
| math | 0.0001 | 0.0004 | 8300.29 | -0.0006 | 0.0008 | 0.273 | 0.78508 |
| motor | 0.0004 | 0.0003 | 7722.01 | -0.0002 | 0.0011 | 1.305 | 0.19178 |
| punish | -0.0005 | 0.0003 | 7617.66 | -0.0011 | 0.0002 | -1.384 | 0.16644 |
| random | -0.0001 | 0.0003 | 7749.94 | -0.0008 | 0.0005 | -0.368 | 0.71276 |
| relational | -0.0007 | 0.0003 | 7923.02 | -0.0013 | 0.0000 | -1.942 | 0.05211 |
| reward | -0.0005 | 0.0003 | 7617.66 | -0.0011 | 0.0002 | -1.365 | 0.17226 |
| shapes | -0.0006 | 0.0003 | 7807.57 | -0.0013 | 0.0001 | -1.775 | 0.07596 |
| story | 0.0003 | 0.0004 | 8300.29 | -0.0004 | 0.0010 | 0.733 | 0.46384 |
| ToM | -0.0003 | 0.0003 | 7749.94 | -0.0010 | 0.0004 | -0.876 | 0.38107 |

| condition | $\beta_{Openness}$ | SE | df | $CL_{lower}$ | $CL_{upper}$ | t.ratio | p.value |
|---|---|---|---|---|---|---|---|
| 0back | 0.0001 | 0.0004 | 7459.87 | -0.0006 | 0.0008 | 0.382 | 0.70238 |
| 2back | 0.0007 | 0.0004 | 7459.86 | 0.0000 | 0.0014 | 1.846 | 0.06487 |
| faces | 0.0006 | 0.0004 | 7672.47 | -0.0001 | 0.0013 | 1.723 | 0.08488 |
| match | -0.0004 | 0.0004 | 7798.44 | -0.0011 | 0.0003 | -1.135 | 0.25624 |
| math | 0.0012 | 0.0004 | 8225.09 | 0.0004 | 0.0019 | 3.152 | 0.00163 |
| motor | 0.0002 | 0.0004 | 7599.32 | -0.0005 | 0.0009 | 0.490 | 0.62435 |
| punish | -0.0005 | 0.0004 | 7388.06 | -0.0012 | 0.0002 | -1.265 | 0.20591 |
| random | 0.0007 | 0.0004 | 7600.45 | 0.0000 | 0.0014 | 1.941 | 0.05234 |
| relational | 0.0002 | 0.0004 | 7798.44 | -0.0005 | 0.0009 | 0.504 | 0.61404 |
| reward | -0.0002 | 0.0004 | 7388.06 | -0.0009 | 0.0005 | -0.689 | 0.49091 |
| shapes | -0.0002 | 0.0004 | 7672.47 | -0.0009 | 0.0005 | -0.457 | 0.64736 |
| story | 0.0007 | 0.0004 | 8225.09 | 0.0000 | 0.0015 | 1.983 | 0.04741 |
| ToM | 0.0002 | 0.0004 | 7600.45 | -0.0005 | 0.0009 | 0.525 | 0.59938 |

| condition | $\beta_{Conscientiousness}$ | SE | df | $CL_{lower}$ | $CL_{upper}$ | t.ratio | p.value |
|---|---|---|---|---|---|---|---|
| 0back | -0.0005 | 0.0004 | 7805.43 | -0.0013 | 0.0003 | -1.178 | 0.23899 |
| 2back | -0.0010 | 0.0004 | 7805.40 | -0.0018 | -0.0002 | -2.568 | 0.01024 |
| faces | -0.0007 | 0.0004 | 7988.24 | -0.0015 | 0.0001 | -1.760 | 0.07842 |
| match | 0.0002 | 0.0004 | 8132.40 | -0.0006 | 0.0010 | 0.504 | 0.61448 |
| math | -0.0004 | 0.0004 | 8510.75 | -0.0012 | 0.0004 | -0.926 | 0.35463 |
| motor | -0.0001 | 0.0004 | 7943.36 | -0.0009 | 0.0007 | -0.206 | 0.83699 |

| | | | | | | | |
|---|---|---|---|---|---|---|---|
| punish | -0.0002 | 0.0004 | 7727.19 | -0.0010 | 0.0005 | -0.614 | 0.53926 |
| random | 0.0003 | 0.0004 | 7999.99 | -0.0005 | 0.0011 | 0.789 | 0.43029 |
| relational | -0.0002 | 0.0004 | 8132.40 | -0.0010 | 0.0006 | -0.519 | 0.60390 |
| reward | -0.0003 | 0.0004 | 7727.19 | -0.0011 | 0.0005 | -0.635 | 0.52554 |
| shapes | -0.0005 | 0.0004 | 7988.24 | -0.0013 | 0.0003 | -1.175 | 0.24014 |
| story | 0.0003 | 0.0004 | 8510.75 | -0.0006 | 0.0011 | 0.626 | 0.53124 |
| ToM | 0.0000 | 0.0004 | 7999.99 | -0.0008 | 0.0008 | -0.057 | 0.95491 |

| **condition** | **$\beta_{Extraversion}$** | **SE** | **df** | **$CL_{lower}$** | **$CL_{upper}$** | **t.ratio** | **p.value** |
|---|---|---|---|---|---|---|---|
| 0back | -0.0005 | 0.0004 | 7784.25 | -0.0012 | 0.0003 | -1.162 | 0.24543 |
| 2back | -0.0005 | 0.0004 | 7784.24 | -0.0013 | 0.0003 | -1.253 | 0.21008 |
| faces | -0.0001 | 0.0004 | 7880.49 | -0.0009 | 0.0007 | -0.329 | 0.74185 |
| match | -0.0003 | 0.0004 | 8062.57 | -0.0011 | 0.0005 | -0.637 | 0.52439 |
| math | 0.0005 | 0.0004 | 8341.40 | -0.0003 | 0.0013 | 1.214 | 0.22475 |
| motor | 0.0010 | 0.0004 | 7952.31 | 0.0002 | 0.0018 | 2.422 | 0.01545 |
| punish | 0.0001 | 0.0004 | 7755.99 | -0.0007 | 0.0009 | 0.201 | 0.84080 |
| random | -0.0007 | 0.0004 | 7843.34 | -0.0015 | 0.0001 | -1.757 | 0.07889 |
| relational | -0.0001 | 0.0004 | 8062.57 | -0.0009 | 0.0007 | -0.167 | 0.86749 |
| reward | 0.0005 | 0.0004 | 7755.99 | -0.0003 | 0.0013 | 1.209 | 0.22672 |
| shapes | 0.0003 | 0.0004 | 7880.49 | -0.0004 | 0.0011 | 0.866 | 0.38649 |
| story | 0.0005 | 0.0004 | 8341.40 | -0.0003 | 0.0013 | 1.312 | 0.18956 |
| ToM | -0.0005 | 0.0004 | 7843.34 | -0.0013 | 0.0002 | -1.378 | 0.16811 |

| **condition** | **$\beta_{Agreeableness}$** | **SE** | **df** | **$CL_{lower}$** | **$CL_{upper}$** | **t.ratio** | **p.value** |
|---|---|---|---|---|---|---|---|
| 0back | 0.0008 | 0.0004 | 7410.17 | 0.0000 | 0.0016 | 2.047 | 0.04069 |
| 2back | 0.0009 | 0.0004 | 7410.16 | 0.0001 | 0.0017 | 2.173 | 0.02981 |
| faces | 0.0005 | 0.0004 | 7536.37 | -0.0003 | 0.0013 | 1.211 | 0.22587 |
| match | 0.0001 | 0.0004 | 7656.48 | -0.0007 | 0.0009 | 0.165 | 0.86896 |
| math | -0.0006 | 0.0004 | 8227.56 | -0.0014 | 0.0003 | -1.354 | 0.17572 |
| motor | -0.0014 | 0.0004 | 7563.87 | -0.0022 | -0.0006 | -3.345 | 0.00083 |
| punish | 0.0001 | 0.0004 | 7417.64 | -0.0007 | 0.0009 | 0.301 | 0.76351 |
| random | 0.0005 | 0.0004 | 7459.81 | -0.0003 | 0.0013 | 1.255 | 0.20966 |
| relational | 0.0005 | 0.0004 | 7656.48 | -0.0003 | 0.0013 | 1.119 | 0.26299 |
| reward | 0.0002 | 0.0004 | 7417.64 | -0.0006 | 0.0010 | 0.595 | 0.55188 |
| shapes | 0.0000 | 0.0004 | 7536.37 | -0.0008 | 0.0008 | -0.026 | 0.97905 |
| story | -0.0012 | 0.0004 | 8227.56 | -0.0020 | -0.0003 | -2.713 | 0.00669 |
| ToM | -0.0003 | 0.0004 | 7459.81 | -0.0011 | 0.0005 | -0.796 | 0.42618 |

**SI Table 6.** Estimated marginal trends (slopes) of personality traits for all task conditions along D3 (DMN- FPN)

| **condition** | **$\beta_{Neuroticism}$** | **SE** | **df** | **$CL_{lower}$** | **$CL_{upper}$** | **t.ratio** | **p.value** |
|---|---|---|---|---|---|---|---|
| 0back | 0.0004 | 0.0004 | 8724.24 | -0.0003 | 0.0011 | 1.069 | 0.28532 |
| 2back | -0.0008 | 0.0004 | 8724.23 | -0.0016 | -0.0001 | -2.217 | 0.02663 |
| faces | -0.0003 | 0.0004 | 8897.12 | -0.0010 | 0.0005 | -0.734 | 0.46312 |
| match | -0.0001 | 0.0004 | 9007.30 | -0.0009 | 0.0006 | -0.315 | 0.75295 |
| math | -0.0007 | 0.0004 | 9367.33 | -0.0015 | 0.0000 | -1.847 | 0.06475 |
| motor | -0.0004 | 0.0004 | 8826.35 | -0.0011 | 0.0003 | -1.075 | 0.28227 |
| punish | 0.0003 | 0.0004 | 8727.02 | -0.0004 | 0.0011 | 0.905 | 0.36544 |
| random | -0.0011 | 0.0004 | 8846.86 | -0.0018 | -0.0004 | -2.914 | 0.00357 |
| relational | -0.0004 | 0.0004 | 9007.30 | -0.0012 | 0.0003 | -1.162 | 0.24507 |
| reward | 0.0003 | 0.0004 | 8727.02 | -0.0004 | 0.0011 | 0.875 | 0.38140 |
| shapes | -0.0003 | 0.0004 | 8897.12 | -0.0011 | 0.0004 | -0.844 | 0.39858 |
| story | 0.0003 | 0.0004 | 9367.33 | -0.0005 | 0.0011 | 0.795 | 0.42683 |
| ToM | -0.0001 | 0.0004 | 8846.86 | -0.0009 | 0.0006 | -0.336 | 0.73677 |
| **condition** | **$\beta_{Openness}$** | **SE** | **df** | **$CL_{lower}$** | **$CL_{upper}$** | **t.ratio** | **p.value** |
| 0back | -0.0009 | 0.0004 | 8587.97 | -0.0017 | -0.0002 | -2.361 | 0.01824 |
| 2back | 0.0005 | 0.0004 | 8587.96 | -0.0002 | 0.0013 | 1.380 | 0.16777 |
| faces | -0.0004 | 0.0004 | 8788.00 | -0.0012 | 0.0004 | -1.006 | 0.31437 |
| match | -0.0007 | 0.0004 | 8904.52 | -0.0015 | 0.0001 | -1.834 | 0.06673 |
| math | 0.0003 | 0.0004 | 9315.51 | -0.0006 | 0.0011 | 0.619 | 0.53613 |
| motor | 0.0006 | 0.0004 | 8720.51 | -0.0002 | 0.0014 | 1.491 | 0.13606 |
| punish | 0.0004 | 0.0004 | 8519.61 | -0.0004 | 0.0012 | 1.062 | 0.28829 |
| random | 0.0007 | 0.0004 | 8718.10 | -0.0001 | 0.0014 | 1.626 | 0.10402 |
| relational | -0.0001 | 0.0004 | 8904.52 | -0.0008 | 0.0007 | -0.128 | 0.89781 |
| reward | 0.0008 | 0.0004 | 8519.61 | 0.0001 | 0.0016 | 2.122 | 0.03386 |
| shapes | 0.0004 | 0.0004 | 8788.00 | -0.0004 | 0.0012 | 0.905 | 0.36556 |
| story | -0.0015 | 0.0004 | 9315.51 | -0.0023 | -0.0007 | -3.598 | 0.00032 |
| ToM | -0.0010 | 0.0004 | 8718.10 | -0.0017 | -0.0002 | -2.375 | 0.01759 |
| **condition** | **$\beta_{Conscientiousness}$** | **SE** | **df** | **$CL_{lower}$** | **$CL_{upper}$** | **t.ratio** | **p.value** |
| 0back | 0.0012 | 0.0005 | 8911.74 | 0.0003 | 0.0021 | 2.663 | 0.00777 |
| 2back | -0.0004 | 0.0005 | 8911.73 | -0.0013 | 0.0005 | -0.881 | 0.37817 |
| faces | -0.0008 | 0.0005 | 9074.25 | -0.0017 | 0.0001 | -1.809 | 0.07043 |
| match | 0.0001 | 0.0005 | 9213.58 | -0.0008 | 0.0010 | 0.252 | 0.80075 |
| math | -0.0001 | 0.0005 | 9568.55 | -0.0011 | 0.0008 | -0.271 | 0.78670 |
| motor | -0.0013 | 0.0005 | 9044.66 | -0.0022 | -0.0004 | -2.775 | 0.00553 |

| | | | | | | | |
|---|---|---|---|---|---|---|---|
| punish | 0.0006 | 0.0004 | 8836.47 | -0.0003 | 0.0015 | 1.288 | 0.19770 |
| random | -0.0003 | 0.0005 | 9090.15 | -0.0012 | 0.0006 | -0.620 | 0.53554 |
| relational | -0.0002 | 0.0005 | 9213.58 | -0.0011 | 0.0007 | -0.473 | 0.63597 |
| reward | 0.0010 | 0.0004 | 8836.47 | 0.0001 | 0.0018 | 2.152 | 0.03142 |
| shapes | -0.0010 | 0.0005 | 9074.25 | -0.0019 | -0.0001 | -2.181 | 0.02919 |
| story | 0.0007 | 0.0005 | 9568.55 | -0.0002 | 0.0017 | 1.572 | 0.11592 |
| ToM | 0.0005 | 0.0005 | 9090.15 | -0.0004 | 0.0014 | 1.087 | 0.27693 |

| **condition** | **$\beta_{Extraversion}$** | **SE** | **df** | **$CL_{lower}$** | **$CL_{upper}$** | **t.ratio** | **p.value** |
|---|---|---|---|---|---|---|---|
| 0back | -0.0004 | 0.0004 | 8895.53 | -0.0012 | 0.0005 | -0.820 | 0.41205 |
| 2back | -0.0008 | 0.0004 | 8895.53 | -0.0016 | 0.0001 | -1.773 | 0.07621 |
| faces | -0.0002 | 0.0004 | 8979.95 | -0.0010 | 0.0007 | -0.352 | 0.72459 |
| match | 0.0001 | 0.0004 | 9156.22 | -0.0008 | 0.0009 | 0.158 | 0.87407 |
| math | -0.0005 | 0.0005 | 9418.43 | -0.0014 | 0.0004 | -0.999 | 0.31763 |
| motor | -0.0008 | 0.0004 | 9057.09 | -0.0017 | 0.0001 | -1.811 | 0.07023 |
| punish | 0.0007 | 0.0004 | 8868.54 | -0.0001 | 0.0016 | 1.637 | 0.10169 |
| random | -0.0006 | 0.0004 | 8946.15 | -0.0014 | 0.0003 | -1.246 | 0.21278 |
| relational | -0.0001 | 0.0004 | 9156.22 | -0.0010 | 0.0008 | -0.249 | 0.80353 |
| reward | 0.0005 | 0.0004 | 8868.54 | -0.0004 | 0.0013 | 1.111 | 0.26670 |
| shapes | 0.0007 | 0.0004 | 8979.95 | -0.0002 | 0.0016 | 1.554 | 0.12014 |
| story | -0.0003 | 0.0005 | 9418.43 | -0.0012 | 0.0006 | -0.599 | 0.54897 |
| ToM | 0.0003 | 0.0004 | 8946.15 | -0.0006 | 0.0011 | 0.614 | 0.53893 |

| **condition** | **$\beta_{Agreeableness}$** | **SE** | **df** | **$CL_{lower}$** | **$CL_{upper}$** | **t.ratio** | **p.value** |
|---|---|---|---|---|---|---|---|
| 0back | -0.0007 | 0.0004 | 8507.89 | -0.0016 | 0.0002 | -1.601 | 0.10938 |
| 2back | -0.0003 | 0.0004 | 8507.88 | -0.0012 | 0.0006 | -0.670 | 0.50316 |
| faces | 0.0004 | 0.0005 | 8622.84 | -0.0004 | 0.0013 | 0.963 | 0.33580 |
| match | 0.0001 | 0.0005 | 8740.11 | -0.0008 | 0.0010 | 0.307 | 0.75913 |
| math | -0.0001 | 0.0005 | 9288.07 | -0.0010 | 0.0009 | -0.128 | 0.89810 |
| motor | 0.0008 | 0.0004 | 8660.37 | -0.0001 | 0.0017 | 1.840 | 0.06580 |
| punish | -0.0002 | 0.0004 | 8511.83 | -0.0010 | 0.0007 | -0.374 | 0.70804 |
| random | -0.0005 | 0.0004 | 8548.72 | -0.0014 | 0.0003 | -1.194 | 0.23238 |
| relational | 0.0003 | 0.0005 | 8740.11 | -0.0006 | 0.0012 | 0.743 | 0.45733 |
| reward | -0.0002 | 0.0004 | 8511.83 | -0.0011 | 0.0007 | -0.402 | 0.68801 |
| shapes | 0.0000 | 0.0005 | 8622.84 | -0.0008 | 0.0009 | 0.105 | 0.91665 |
| story | -0.0002 | 0.0005 | 9288.07 | -0.0011 | 0.0007 | -0.462 | 0.64402 |
| ToM | -0.0013 | 0.0004 | 8548.72 | -0.0021 | -0.0004 | -2.840 | 0.00452 |

**SI Table 7.1** Neuroticism* task condition interactions along D1 (primary- association)

| contrast | β | SE | df | $CL_{lower}$ | $CL_{upper}$ | t.ratio | p.value |
|---|---|---|---|---|---|---|---|
| 0back - 2back | 0.0010 | 0.0006 | 11916.92 | -0.0001 | 0.0021 | 1.80 | 0.07253 |
| 0back - faces | 0.0010 | 0.0006 | 11962.92 | -0.0001 | 0.0021 | 1.81 | 0.06980 |
| 0back - match | 0.0002 | 0.0006 | 11965.96 | -0.0009 | 0.0014 | 0.41 | 0.68311 |
| 0back - math | 0.0001 | 0.0006 | 11973.37 | -0.0010 | 0.0013 | 0.25 | 0.80598 |
| 0back - motor | -0.0012 | 0.0006 | 11933.72 | -0.0023 | -0.0001 | -2.18 | 0.02954 |
| 0back - punish | 0.0013 | 0.0006 | 11936.66 | 0.0002 | 0.0024 | 2.31 | 0.02109 |
| 0back - random | -0.0004 | 0.0006 | 11969.37 | -0.0015 | 0.0007 | -0.64 | 0.52261 |
| 0back - relational | 0.0002 | 0.0006 | 11965.96 | -0.0009 | 0.0013 | 0.39 | 0.69392 |
| 0back - reward | 0.0007 | 0.0006 | 11936.66 | -0.0004 | 0.0018 | 1.21 | 0.22574 |
| 0back - shapes | 0.0006 | 0.0006 | 11962.92 | -0.0005 | 0.0017 | 1.04 | 0.29787 |
| 0back - story | 0.0010 | 0.0006 | 11973.37 | -0.0002 | 0.0021 | 1.66 | 0.09702 |
| 0back - ToM | 0.0002 | 0.0006 | 11969.37 | -0.0009 | 0.0013 | 0.43 | 0.67053 |
| 2back - faces | 0.0000 | 0.0006 | 11962.92 | -0.0011 | 0.0011 | 0.04 | 0.97051 |
| 2back - match | -0.0008 | 0.0006 | 11965.96 | -0.0019 | 0.0003 | -1.36 | 0.17410 |
| 2back - math | -0.0009 | 0.0006 | 11973.37 | -0.0020 | 0.0003 | -1.49 | 0.13517 |
| 2back - motor | -0.0022 | 0.0006 | 11933.73 | -0.0033 | -0.0011 | -3.96 | 0.00007 |
| 2back - punish | 0.0003 | 0.0006 | 11936.66 | -0.0008 | 0.0014 | 0.51 | 0.60846 |
| 2back - random | -0.0014 | 0.0006 | 11969.37 | -0.0025 | -0.0003 | -2.42 | 0.01550 |
| 2back - relational | -0.0008 | 0.0006 | 11965.96 | -0.0019 | 0.0003 | -1.37 | 0.16950 |
| 2back - reward | -0.0003 | 0.0006 | 11936.66 | -0.0014 | 0.0008 | -0.58 | 0.55998 |
| 2back - shapes | -0.0004 | 0.0006 | 11962.92 | -0.0015 | 0.0007 | -0.74 | 0.46214 |
| 2back - story | 0.0000 | 0.0006 | 11973.37 | -0.0012 | 0.0011 | -0.08 | 0.93615 |
| 2back - ToM | -0.0008 | 0.0006 | 11969.37 | -0.0019 | 0.0003 | -1.36 | 0.17514 |
| faces - match | -0.0008 | 0.0006 | 11928.41 | -0.0019 | 0.0003 | -1.39 | 0.16551 |
| faces - math | -0.0009 | 0.0006 | 11946.63 | -0.0020 | 0.0003 | -1.52 | 0.12861 |
| faces - motor | -0.0023 | 0.0006 | 11970.62 | -0.0034 | -0.0011 | -3.96 | 0.00008 |
| faces - punish | 0.0003 | 0.0006 | 11961.69 | -0.0008 | 0.0014 | 0.47 | 0.63814 |
| faces - random | -0.0014 | 0.0006 | 11932.32 | -0.0025 | -0.0003 | -2.44 | 0.01465 |
| faces - relational | -0.0008 | 0.0006 | 11928.41 | -0.0019 | 0.0003 | -1.40 | 0.16111 |
| faces - reward | -0.0003 | 0.0006 | 11961.69 | -0.0015 | 0.0008 | -0.61 | 0.53912 |
| faces - shapes | -0.0004 | 0.0006 | 11916.92 | -0.0016 | 0.0007 | -0.77 | 0.44275 |
| faces - story | -0.0001 | 0.0006 | 11946.63 | -0.0012 | 0.0011 | -0.12 | 0.90805 |
| faces - ToM | -0.0008 | 0.0006 | 11932.32 | -0.0019 | 0.0003 | -1.38 | 0.16650 |
| match - math | -0.0001 | 0.0006 | 11940.95 | -0.0012 | 0.0011 | -0.15 | 0.87725 |
| match - motor | -0.0015 | 0.0006 | 11967.67 | -0.0026 | -0.0003 | -2.55 | 0.01077 |
| match - punish | 0.0011 | 0.0006 | 11969.29 | -0.0001 | 0.0022 | 1.86 | 0.06238 |
| match - random | -0.0006 | 0.0006 | 11938.84 | -0.0017 | 0.0005 | -1.04 | 0.29918 |
| match - relational | 0.0000 | 0.0006 | 11916.92 | -0.0011 | 0.0011 | -0.01 | 0.98841 |
| match - reward | 0.0004 | 0.0006 | 11969.29 | -0.0007 | 0.0016 | 0.79 | 0.43243 |

| | | | | | | | |
|---|---|---|---|---|---|---|---|
| match - shapes | 0.0004 | 0.0006 | 11928.41 | -0.0008 | 0.0015 | 0.62 | 0.53299 |
| match - story | 0.0007 | 0.0006 | 11940.95 | -0.0004 | 0.0019 | 1.24 | 0.21364 |
| match - ToM | 0.0000 | 0.0006 | 11938.84 | -0.0011 | 0.0011 | 0.01 | 0.98897 |
| math - motor | -0.0014 | 0.0006 | 11978.19 | -0.0025 | -0.0002 | -2.35 | 0.01855 |
| math - punish | 0.0012 | 0.0006 | 11979.96 | 0.0000 | 0.0023 | 1.99 | 0.04658 |
| math - random | -0.0005 | 0.0006 | 11952.01 | -0.0016 | 0.0006 | -0.87 | 0.38636 |
| math - relational | 0.0001 | 0.0006 | 11940.95 | -0.0011 | 0.0012 | 0.14 | 0.88853 |
| math - reward | 0.0005 | 0.0006 | 11979.96 | -0.0006 | 0.0017 | 0.93 | 0.35305 |
| math - shapes | 0.0004 | 0.0006 | 11946.63 | -0.0007 | 0.0016 | 0.77 | 0.44216 |
| math - story | 0.0008 | 0.0006 | 11916.92 | -0.0003 | 0.0020 | 1.38 | 0.16735 |
| math - ToM | 0.0001 | 0.0006 | 11952.01 | -0.0010 | 0.0012 | 0.17 | 0.86569 |
| motor - punish | 0.0025 | 0.0006 | 11936.29 | 0.0014 | 0.0036 | 4.47 | 0.00001 |
| motor - random | 0.0009 | 0.0006 | 11978.50 | -0.0002 | 0.0020 | 1.52 | 0.12768 |
| motor - relational | 0.0015 | 0.0006 | 11967.67 | 0.0003 | 0.0026 | 2.54 | 0.01123 |
| motor - reward | 0.0019 | 0.0006 | 11936.29 | 0.0008 | 0.0030 | 3.38 | 0.00072 |
| motor - shapes | 0.0018 | 0.0006 | 11970.62 | 0.0007 | 0.0029 | 3.19 | 0.00142 |
| motor - story | 0.0022 | 0.0006 | 11978.19 | 0.0010 | 0.0033 | 3.76 | 0.00017 |
| motor - ToM | 0.0015 | 0.0006 | 11978.50 | 0.0004 | 0.0026 | 2.58 | 0.00980 |
| punish - random | -0.0017 | 0.0006 | 11968.10 | -0.0028 | -0.0005 | -2.93 | 0.00340 |
| punish - relational | -0.0011 | 0.0006 | 11969.29 | -0.0022 | 0.0000 | -1.88 | 0.06034 |
| punish - reward | -0.0006 | 0.0006 | 11916.92 | -0.0017 | 0.0005 | -1.10 | 0.27296 |
| punish - shapes | -0.0007 | 0.0006 | 11961.69 | -0.0018 | 0.0004 | -1.24 | 0.21396 |
| punish - story | -0.0003 | 0.0006 | 11979.96 | -0.0015 | 0.0008 | -0.58 | 0.56419 |
| punish - ToM | -0.0011 | 0.0006 | 11968.10 | -0.0022 | 0.0001 | -1.86 | 0.06223 |
| random - relational | 0.0006 | 0.0006 | 11938.84 | -0.0005 | 0.0017 | 1.02 | 0.30604 |
| random - reward | 0.0010 | 0.0006 | 11968.10 | -0.0001 | 0.0022 | 1.84 | 0.06545 |
| random - shapes | 0.0010 | 0.0006 | 11932.32 | -0.0002 | 0.0021 | 1.67 | 0.09456 |
| random - story | 0.0013 | 0.0006 | 11952.01 | 0.0002 | 0.0025 | 2.27 | 0.02297 |
| random - ToM | 0.0006 | 0.0006 | 11916.92 | -0.0005 | 0.0017 | 1.06 | 0.28856 |
| relational - reward | 0.0005 | 0.0006 | 11969.29 | -0.0007 | 0.0016 | 0.80 | 0.42388 |
| relational - shapes | 0.0004 | 0.0006 | 11928.41 | -0.0008 | 0.0015 | 0.64 | 0.52346 |
| relational - story | 0.0007 | 0.0006 | 11940.95 | -0.0004 | 0.0019 | 1.26 | 0.20843 |
| relational - ToM | 0.0000 | 0.0006 | 11938.84 | -0.0011 | 0.0011 | 0.03 | 0.97731 |
| reward - shapes | -0.0001 | 0.0006 | 11961.69 | -0.0012 | 0.0010 | -0.16 | 0.87419 |
| reward - story | 0.0003 | 0.0006 | 11979.96 | -0.0009 | 0.0014 | 0.48 | 0.62776 |
| reward - ToM | -0.0004 | 0.0006 | 11968.10 | -0.0015 | 0.0007 | -0.78 | 0.43694 |
| shapes - story | 0.0004 | 0.0006 | 11946.63 | -0.0008 | 0.0015 | 0.64 | 0.52504 |
| shapes - ToM | -0.0003 | 0.0006 | 11932.32 | -0.0015 | 0.0008 | -0.61 | 0.53899 |
| story - ToM | -0.0007 | 0.0006 | 11952.01 | -0.0019 | 0.0004 | -1.24 | 0.21546 |

**SI Table 7.2** Openness* task condition interactions along D1 (primary- association)

| contrast | β | SE | df | CL $_{lower}$ | CL $_{upper}$ | t.ratio | p.value |
|---|---|---|---|---|---|---|---|
| 0back - 2back | 0.0004 | 0.0006 | 11916.92 | -0.0007 | 0.0016 | 0.72 | 0.47073 |
| 0back - faces | -0.0008 | 0.0006 | 11958.88 | -0.0020 | 0.0004 | -1.28 | 0.19882 |
| 0back - match | 0.0015 | 0.0006 | 11963.39 | 0.0004 | 0.0027 | 2.56 | 0.01035 |
| 0back - math | -0.0004 | 0.0006 | 11975.61 | -0.0016 | 0.0008 | -0.70 | 0.48675 |
| 0back - motor | 0.0004 | 0.0006 | 11935.74 | -0.0008 | 0.0016 | 0.69 | 0.49206 |
| 0back - punish | 0.0000 | 0.0006 | 11933.93 | -0.0011 | 0.0012 | 0.06 | 0.95443 |
| 0back - random | -0.0005 | 0.0006 | 11960.10 | -0.0017 | 0.0007 | -0.85 | 0.39349 |
| 0back - relational | 0.0009 | 0.0006 | 11963.39 | -0.0002 | 0.0021 | 1.55 | 0.12074 |
| 0back - reward | 0.0006 | 0.0006 | 11933.93 | -0.0006 | 0.0017 | 0.98 | 0.32909 |
| 0back - shapes | -0.0005 | 0.0006 | 11958.88 | -0.0016 | 0.0007 | -0.76 | 0.44594 |
| 0back - story | -0.0015 | 0.0006 | 11975.61 | -0.0027 | -0.0003 | -2.50 | 0.01259 |
| 0back - ToM | 0.0003 | 0.0006 | 11960.10 | -0.0008 | 0.0015 | 0.58 | 0.56428 |
| 2back - faces | -0.0012 | 0.0006 | 11958.89 | -0.0024 | 0.0000 | -2.00 | 0.04570 |
| 2back - match | 0.0011 | 0.0006 | 11963.40 | -0.0001 | 0.0023 | 1.85 | 0.06362 |
| 2back - math | -0.0009 | 0.0006 | 11975.62 | -0.0021 | 0.0003 | -1.39 | 0.16395 |
| 2back - motor | 0.0000 | 0.0006 | 11935.74 | -0.0012 | 0.0012 | -0.03 | 0.97651 |
| 2back - punish | -0.0004 | 0.0006 | 11933.93 | -0.0016 | 0.0008 | -0.67 | 0.50565 |
| 2back - random | -0.0009 | 0.0006 | 11960.11 | -0.0021 | 0.0002 | -1.57 | 0.11670 |
| 2back - relational | 0.0005 | 0.0006 | 11963.40 | -0.0007 | 0.0017 | 0.84 | 0.39955 |
| 2back - reward | 0.0002 | 0.0006 | 11933.93 | -0.0010 | 0.0013 | 0.25 | 0.80012 |
| 2back - shapes | -0.0009 | 0.0006 | 11958.89 | -0.0021 | 0.0003 | -1.48 | 0.14008 |
| 2back - story | -0.0020 | 0.0006 | 11975.62 | -0.0032 | -0.0008 | -3.19 | 0.00142 |
| 2back - ToM | -0.0001 | 0.0006 | 11960.11 | -0.0013 | 0.0011 | -0.14 | 0.88943 |
| faces - match | 0.0023 | 0.0006 | 11934.94 | 0.0011 | 0.0035 | 3.81 | 0.00014 |
| faces - math | 0.0003 | 0.0006 | 11953.48 | -0.0009 | 0.0016 | 0.56 | 0.57882 |
| faces - motor | 0.0012 | 0.0006 | 11968.42 | 0.0000 | 0.0024 | 1.96 | 0.05033 |
| faces - punish | 0.0008 | 0.0006 | 11960.23 | -0.0004 | 0.0020 | 1.35 | 0.17852 |
| faces - random | 0.0003 | 0.0006 | 11926.83 | -0.0009 | 0.0014 | 0.43 | 0.66534 |
| faces - relational | 0.0017 | 0.0006 | 11934.94 | 0.0005 | 0.0029 | 2.81 | 0.00497 |
| faces - reward | 0.0014 | 0.0006 | 11960.23 | 0.0002 | 0.0025 | 2.25 | 0.02415 |
| faces - shapes | 0.0003 | 0.0006 | 11916.92 | -0.0009 | 0.0015 | 0.52 | 0.60346 |
| faces - story | -0.0008 | 0.0006 | 11953.48 | -0.0020 | 0.0005 | -1.23 | 0.21788 |
| faces - ToM | 0.0011 | 0.0006 | 11926.83 | -0.0001 | 0.0023 | 1.85 | 0.06399 |
| match - math | -0.0020 | 0.0006 | 11943.47 | -0.0032 | -0.0008 | -3.18 | 0.00150 |
| match - motor | -0.0011 | 0.0006 | 11965.65 | -0.0023 | 0.0001 | -1.87 | 0.06097 |
| match - punish | -0.0015 | 0.0006 | 11971.11 | -0.0027 | -0.0003 | -2.51 | 0.01193 |
| match - random | -0.0021 | 0.0006 | 11939.24 | -0.0033 | -0.0009 | -3.40 | 0.00069 |
| match - relational | -0.0006 | 0.0006 | 11916.92 | -0.0018 | 0.0006 | -1.00 | 0.31674 |
| match - reward | -0.0010 | 0.0006 | 11971.11 | -0.0022 | 0.0002 | -1.61 | 0.10728 |

| | | | | | | | |
|---|---|---|---|---|---|---|---|
| match - shapes | -0.0020 | 0.0006 | 11934.94 | -0.0032 | -0.0008 | -3.30 | 0.00097 |
| match - story | -0.0031 | 0.0006 | 11943.47 | -0.0043 | -0.0019 | -4.96 | 0.00000 |
| match - ToM | -0.0012 | 0.0006 | 11939.24 | -0.0024 | 0.0000 | -1.98 | 0.04726 |
| math - motor | 0.0008 | 0.0006 | 11975.05 | -0.0004 | 0.0021 | 1.36 | 0.17498 |
| math - punish | 0.0005 | 0.0006 | 11982.14 | -0.0007 | 0.0017 | 0.75 | 0.45179 |
| math - random | -0.0001 | 0.0006 | 11952.04 | -0.0013 | 0.0011 | -0.13 | 0.89299 |
| math - relational | 0.0014 | 0.0006 | 11943.47 | 0.0001 | 0.0026 | 2.19 | 0.02829 |
| math - reward | 0.0010 | 0.0006 | 11982.14 | -0.0002 | 0.0022 | 1.64 | 0.10102 |
| math - shapes | 0.0000 | 0.0006 | 11953.48 | -0.0012 | 0.0012 | -0.05 | 0.96144 |
| math - story | -0.0011 | 0.0006 | 11916.92 | -0.0023 | 0.0001 | -1.75 | 0.07939 |
| math - ToM | 0.0008 | 0.0006 | 11952.04 | -0.0004 | 0.0020 | 1.25 | 0.21080 |
| motor - punish | -0.0004 | 0.0006 | 11936.28 | -0.0015 | 0.0008 | -0.63 | 0.52724 |
| motor - random | -0.0009 | 0.0006 | 11967.63 | -0.0021 | 0.0003 | -1.53 | 0.12594 |
| motor - relational | 0.0005 | 0.0006 | 11965.65 | -0.0007 | 0.0017 | 0.87 | 0.38603 |
| motor - reward | 0.0002 | 0.0006 | 11936.28 | -0.0010 | 0.0013 | 0.28 | 0.77852 |
| motor - shapes | -0.0009 | 0.0006 | 11968.42 | -0.0021 | 0.0003 | -1.44 | 0.15054 |
| motor - story | -0.0019 | 0.0006 | 11975.05 | -0.0032 | -0.0007 | -3.15 | 0.00165 |
| motor - ToM | -0.0001 | 0.0006 | 11967.63 | -0.0012 | 0.0011 | -0.11 | 0.91322 |
| punish - random | -0.0005 | 0.0006 | 11955.72 | -0.0017 | 0.0006 | -0.91 | 0.36137 |
| punish - relational | 0.0009 | 0.0006 | 11971.11 | -0.0003 | 0.0021 | 1.50 | 0.13375 |
| punish - reward | 0.0005 | 0.0006 | 11916.92 | -0.0006 | 0.0017 | 0.92 | 0.35636 |
| punish - shapes | -0.0005 | 0.0006 | 11960.23 | -0.0017 | 0.0007 | -0.82 | 0.41162 |
| punish - story | -0.0016 | 0.0006 | 11982.14 | -0.0028 | -0.0004 | -2.56 | 0.01057 |
| punish - ToM | 0.0003 | 0.0006 | 11955.72 | -0.0009 | 0.0015 | 0.52 | 0.60196 |
| random - relational | 0.0014 | 0.0006 | 11939.24 | 0.0003 | 0.0026 | 2.39 | 0.01699 |
| random - reward | 0.0011 | 0.0006 | 11955.72 | -0.0001 | 0.0023 | 1.83 | 0.06798 |
| random - shapes | 0.0001 | 0.0006 | 11926.83 | -0.0011 | 0.0012 | 0.09 | 0.92977 |
| random - story | -0.0010 | 0.0006 | 11952.04 | -0.0022 | 0.0002 | -1.66 | 0.09731 |
| random - ToM | 0.0009 | 0.0006 | 11916.92 | -0.0003 | 0.0020 | 1.42 | 0.15423 |
| relational - reward | -0.0004 | 0.0006 | 11971.11 | -0.0015 | 0.0008 | -0.60 | 0.55151 |
| relational - shapes | -0.0014 | 0.0006 | 11934.94 | -0.0026 | -0.0002 | -2.29 | 0.02184 |
| relational - story | -0.0025 | 0.0006 | 11943.47 | -0.0037 | -0.0013 | -3.97 | 0.00007 |
| relational - ToM | -0.0006 | 0.0006 | 11939.24 | -0.0018 | 0.0006 | -0.98 | 0.32908 |
| reward - shapes | -0.0010 | 0.0006 | 11960.23 | -0.0022 | 0.0001 | -1.73 | 0.08354 |
| reward - story | -0.0021 | 0.0006 | 11982.14 | -0.0033 | -0.0009 | -3.44 | 0.00057 |
| reward - ToM | -0.0002 | 0.0006 | 11955.72 | -0.0014 | 0.0009 | -0.39 | 0.69585 |
| shapes - story | -0.0011 | 0.0006 | 11953.48 | -0.0023 | 0.0001 | -1.74 | 0.08205 |
| shapes - ToM | 0.0008 | 0.0006 | 11926.83 | -0.0004 | 0.0020 | 1.33 | 0.18299 |
| story - ToM | 0.0019 | 0.0006 | 11952.04 | 0.0007 | 0.0031 | 3.04 | 0.00234 |

**SI Table 7.3** Conscientiousness* task condition interactions along D1 (primary- association)

| **contrast** | **β** | **SE** | **df** | **$CL_{lower}$** | **$CL_{upper}$** | **t.ratio** | **p.value** |
|---|---|---|---|---|---|---|---|
| 0back - 2back | -0.0004 | 0.0007 | 11916.91 | -0.0018 | 0.0009 | -0.62 | 0.53701 |
| 0back - faces | 0.0002 | 0.0007 | 11970.61 | -0.0011 | 0.0016 | 0.36 | 0.72202 |
| 0back - match | -0.0013 | 0.0007 | 11974.23 | -0.0026 | 0.0001 | -1.85 | 0.06435 |
| 0back - math | -0.0009 | 0.0007 | 11988.65 | -0.0023 | 0.0005 | -1.31 | 0.19116 |
| 0back - motor | -0.0001 | 0.0007 | 11929.53 | -0.0015 | 0.0012 | -0.20 | 0.83911 |
| 0back - punish | 0.0005 | 0.0007 | 11936.98 | -0.0008 | 0.0019 | 0.78 | 0.43434 |
| 0back - random | -0.0010 | 0.0007 | 11977.49 | -0.0023 | 0.0004 | -1.43 | 0.15146 |
| 0back - relational | -0.0013 | 0.0007 | 11974.23 | -0.0027 | 0.0000 | -1.94 | 0.05223 |
| 0back - reward | 0.0006 | 0.0007 | 11936.98 | -0.0007 | 0.0019 | 0.87 | 0.38264 |
| 0back - shapes | -0.0003 | 0.0007 | 11970.61 | -0.0017 | 0.0010 | -0.50 | 0.61930 |
| 0back - story | -0.0003 | 0.0007 | 11988.65 | -0.0017 | 0.0011 | -0.44 | 0.66008 |
| 0back - ToM | -0.0006 | 0.0007 | 11977.49 | -0.0020 | 0.0007 | -0.90 | 0.36884 |
| 2back - faces | 0.0007 | 0.0007 | 11970.62 | -0.0007 | 0.0020 | 0.97 | 0.33387 |
| 2back - match | -0.0009 | 0.0007 | 11974.23 | -0.0022 | 0.0005 | -1.24 | 0.21395 |
| 2back - math | -0.0005 | 0.0007 | 11988.66 | -0.0019 | 0.0009 | -0.71 | 0.47772 |
| 2back - motor | 0.0003 | 0.0007 | 11929.54 | -0.0011 | 0.0016 | 0.41 | 0.68139 |
| 2back - punish | 0.0010 | 0.0007 | 11936.99 | -0.0004 | 0.0023 | 1.40 | 0.16141 |
| 2back - random | -0.0006 | 0.0007 | 11977.49 | -0.0019 | 0.0008 | -0.82 | 0.40990 |
| 2back - relational | -0.0009 | 0.0007 | 11974.23 | -0.0023 | 0.0004 | -1.33 | 0.18213 |
| 2back - reward | 0.0010 | 0.0007 | 11936.99 | -0.0003 | 0.0023 | 1.49 | 0.13580 |
| 2back - shapes | 0.0001 | 0.0007 | 11970.62 | -0.0013 | 0.0014 | 0.11 | 0.90942 |
| 2back - story | 0.0001 | 0.0007 | 11988.66 | -0.0013 | 0.0015 | 0.16 | 0.87491 |
| 2back - ToM | -0.0002 | 0.0007 | 11977.49 | -0.0015 | 0.0012 | -0.29 | 0.77313 |
| faces - match | -0.0015 | 0.0007 | 11927.96 | -0.0029 | -0.0002 | -2.19 | 0.02846 |
| faces - math | -0.0012 | 0.0007 | 11948.56 | -0.0026 | 0.0002 | -1.65 | 0.09984 |
| faces - motor | -0.0004 | 0.0007 | 11976.07 | -0.0017 | 0.0010 | -0.55 | 0.57907 |
| faces - punish | 0.0003 | 0.0007 | 11966.23 | -0.0011 | 0.0016 | 0.42 | 0.67646 |
| faces - random | -0.0012 | 0.0007 | 11932.89 | -0.0026 | 0.0001 | -1.78 | 0.07516 |
| faces - relational | -0.0016 | 0.0007 | 11927.96 | -0.0030 | -0.0002 | -2.28 | 0.02250 |
| faces - reward | 0.0003 | 0.0007 | 11966.23 | -0.0010 | 0.0017 | 0.51 | 0.61169 |
| faces - shapes | -0.0006 | 0.0007 | 11916.92 | -0.0019 | 0.0008 | -0.85 | 0.39643 |
| faces - story | -0.0006 | 0.0007 | 11948.56 | -0.0019 | 0.0008 | -0.78 | 0.43348 |
| faces - ToM | -0.0009 | 0.0007 | 11932.89 | -0.0022 | 0.0005 | -1.25 | 0.21243 |
| match - math | 0.0004 | 0.0007 | 11946.87 | -0.0010 | 0.0018 | 0.51 | 0.61206 |
| match - motor | 0.0011 | 0.0007 | 11972.99 | -0.0002 | 0.0025 | 1.64 | 0.10087 |
| match - punish | 0.0018 | 0.0007 | 11972.05 | 0.0005 | 0.0032 | 2.63 | 0.00867 |
| match - random | 0.0003 | 0.0007 | 11940.67 | -0.0011 | 0.0017 | 0.42 | 0.67411 |
| match - relational | -0.0001 | 0.0007 | 11916.92 | -0.0014 | 0.0013 | -0.09 | 0.92789 |
| match - reward | 0.0019 | 0.0007 | 11972.05 | 0.0005 | 0.0032 | 2.72 | 0.00663 |

| | | | | | | | |
|---|---|---|---|---|---|---|---|
| match - shapes | 0.0009 | 0.0007 | 11927.96 | -0.0004 | 0.0023 | 1.35 | 0.17759 |
| match - story | 0.0010 | 0.0007 | 11946.87 | -0.0004 | 0.0024 | 1.37 | 0.17221 |
| match - ToM | 0.0007 | 0.0007 | 11940.67 | -0.0007 | 0.0020 | 0.95 | 0.34210 |
| math - motor | 0.0008 | 0.0007 | 11981.54 | -0.0006 | 0.0022 | 1.10 | 0.26928 |
| math - punish | 0.0015 | 0.0007 | 11985.32 | 0.0001 | 0.0028 | 2.07 | 0.03862 |
| math - random | -0.0001 | 0.0007 | 11956.75 | -0.0015 | 0.0013 | -0.10 | 0.92373 |
| math - relational | -0.0004 | 0.0007 | 11946.87 | -0.0018 | 0.0010 | -0.60 | 0.55110 |
| math - reward | 0.0015 | 0.0007 | 11985.32 | 0.0001 | 0.0029 | 2.16 | 0.03104 |
| math - shapes | 0.0006 | 0.0007 | 11948.56 | -0.0008 | 0.0020 | 0.82 | 0.41419 |
| math - story | 0.0006 | 0.0007 | 11916.92 | -0.0008 | 0.0020 | 0.85 | 0.39668 |
| math - ToM | 0.0003 | 0.0007 | 11956.75 | -0.0011 | 0.0017 | 0.43 | 0.67065 |
| motor - punish | 0.0007 | 0.0007 | 11941.61 | -0.0007 | 0.0020 | 0.98 | 0.32660 |
| motor - random | -0.0008 | 0.0007 | 11982.76 | -0.0022 | 0.0005 | -1.23 | 0.22031 |
| motor - relational | -0.0012 | 0.0007 | 11972.99 | -0.0026 | 0.0002 | -1.73 | 0.08334 |
| motor - reward | 0.0007 | 0.0007 | 11941.61 | -0.0006 | 0.0021 | 1.07 | 0.28385 |
| motor - shapes | -0.0002 | 0.0007 | 11976.07 | -0.0016 | 0.0012 | -0.29 | 0.76938 |
| motor - story | -0.0002 | 0.0007 | 11981.54 | -0.0016 | 0.0012 | -0.24 | 0.80945 |
| motor - ToM | -0.0005 | 0.0007 | 11982.76 | -0.0018 | 0.0009 | -0.69 | 0.48838 |
| punish - random | -0.0015 | 0.0007 | 11966.70 | -0.0029 | -0.0002 | -2.21 | 0.02687 |
| punish - relational | -0.0019 | 0.0007 | 11972.05 | -0.0032 | -0.0005 | -2.72 | 0.00660 |
| punish - reward | 0.0001 | 0.0007 | 11916.92 | -0.0013 | 0.0014 | 0.09 | 0.92699 |
| punish - shapes | -0.0009 | 0.0007 | 11966.23 | -0.0022 | 0.0005 | -1.27 | 0.20313 |
| punish - story | -0.0008 | 0.0007 | 11985.32 | -0.0022 | 0.0005 | -1.20 | 0.23081 |
| punish - ToM | -0.0011 | 0.0007 | 11966.70 | -0.0025 | 0.0002 | -1.68 | 0.09376 |
| random - relational | -0.0004 | 0.0007 | 11940.67 | -0.0017 | 0.0010 | -0.51 | 0.60909 |
| random - reward | 0.0016 | 0.0007 | 11966.70 | 0.0002 | 0.0029 | 2.30 | 0.02124 |
| random - shapes | 0.0006 | 0.0007 | 11932.89 | -0.0007 | 0.0020 | 0.93 | 0.35118 |
| random - story | 0.0007 | 0.0007 | 11956.75 | -0.0007 | 0.0021 | 0.96 | 0.33819 |
| random - ToM | 0.0004 | 0.0007 | 11916.92 | -0.0010 | 0.0017 | 0.53 | 0.59397 |
| relational - reward | 0.0019 | 0.0007 | 11972.05 | 0.0006 | 0.0033 | 2.81 | 0.00501 |
| relational - shapes | 0.0010 | 0.0007 | 11927.96 | -0.0004 | 0.0024 | 1.44 | 0.15011 |
| relational - story | 0.0010 | 0.0007 | 11946.87 | -0.0004 | 0.0024 | 1.45 | 0.14592 |
| relational - ToM | 0.0007 | 0.0007 | 11940.67 | -0.0006 | 0.0021 | 1.04 | 0.29794 |
| reward - shapes | -0.0009 | 0.0007 | 11966.23 | -0.0023 | 0.0004 | -1.36 | 0.17287 |
| reward - story | -0.0009 | 0.0007 | 11985.32 | -0.0023 | 0.0005 | -1.29 | 0.19822 |
| reward - ToM | -0.0012 | 0.0007 | 11966.70 | -0.0026 | 0.0001 | -1.77 | 0.07736 |
| shapes - story | 0.0000 | 0.0007 | 11948.56 | -0.0014 | 0.0014 | 0.05 | 0.96340 |
| shapes - ToM | -0.0003 | 0.0007 | 11932.89 | -0.0016 | 0.0011 | -0.40 | 0.68941 |
| story - ToM | -0.0003 | 0.0007 | 11956.75 | -0.0017 | 0.0011 | -0.44 | 0.66228 |

**SI Table 7.4** Extraversion* task condition interactions along D1 (primary- association)

| contrast | β | SE | df | CL lower | CL upper | t.ratio | p.value |
|---|---|---|---|---|---|---|---|
| 0back - 2back | -0.0006 | 0.0007 | 11916.92 | -0.0019 | 0.0007 | -0.92 | 0.36009 |
| 0back - faces | -0.0011 | 0.0007 | 11953.73 | -0.0024 | 0.0002 | -1.59 | 0.11079 |
| 0back - match | -0.0003 | 0.0007 | 11961.95 | -0.0016 | 0.0010 | -0.40 | 0.68690 |
| 0back - math | -0.0003 | 0.0007 | 11974.25 | -0.0016 | 0.0010 | -0.44 | 0.66266 |
| 0back - motor | -0.0015 | 0.0007 | 11935.74 | -0.0028 | -0.0002 | -2.28 | 0.02253 |
| 0back - punish | -0.0004 | 0.0007 | 11927.35 | -0.0017 | 0.0009 | -0.60 | 0.54930 |
| 0back - random | 0.0005 | 0.0007 | 11955.91 | -0.0008 | 0.0018 | 0.74 | 0.46103 |
| 0back - relational | -0.0007 | 0.0007 | 11961.95 | -0.0020 | 0.0006 | -1.08 | 0.28231 |
| 0back - reward | -0.0009 | 0.0007 | 11927.35 | -0.0022 | 0.0004 | -1.37 | 0.17188 |
| 0back - shapes | -0.0011 | 0.0007 | 11953.73 | -0.0024 | 0.0002 | -1.70 | 0.08879 |
| 0back - story | 0.0003 | 0.0007 | 11974.25 | -0.0011 | 0.0016 | 0.41 | 0.67967 |
| 0back - ToM | -0.0001 | 0.0007 | 11955.91 | -0.0014 | 0.0012 | -0.10 | 0.92390 |
| 2back - faces | -0.0005 | 0.0007 | 11953.73 | -0.0018 | 0.0008 | -0.69 | 0.49309 |
| 2back - match | 0.0003 | 0.0007 | 11961.95 | -0.0010 | 0.0017 | 0.50 | 0.61747 |
| 2back - math | 0.0003 | 0.0007 | 11974.25 | -0.0010 | 0.0016 | 0.46 | 0.64879 |
| 2back - motor | -0.0009 | 0.0007 | 11935.74 | -0.0022 | 0.0004 | -1.37 | 0.16966 |
| 2back - punish | 0.0002 | 0.0007 | 11927.35 | -0.0011 | 0.0015 | 0.32 | 0.75138 |
| 2back - random | 0.0011 | 0.0007 | 11955.91 | -0.0002 | 0.0024 | 1.65 | 0.09936 |
| 2back - relational | -0.0001 | 0.0007 | 11961.95 | -0.0014 | 0.0012 | -0.17 | 0.86291 |
| 2back - reward | -0.0003 | 0.0007 | 11927.35 | -0.0016 | 0.0010 | -0.45 | 0.65227 |
| 2back - shapes | -0.0005 | 0.0007 | 11953.73 | -0.0018 | 0.0008 | -0.79 | 0.42802 |
| 2back - story | 0.0009 | 0.0007 | 11974.25 | -0.0004 | 0.0022 | 1.30 | 0.19204 |
| 2back - ToM | 0.0005 | 0.0007 | 11955.91 | -0.0008 | 0.0018 | 0.82 | 0.41485 |
| faces - match | 0.0008 | 0.0007 | 11931.56 | -0.0005 | 0.0021 | 1.18 | 0.23936 |
| faces - math | 0.0008 | 0.0007 | 11950.03 | -0.0006 | 0.0021 | 1.12 | 0.26091 |
| faces - motor | -0.0005 | 0.0007 | 11960.01 | -0.0018 | 0.0009 | -0.69 | 0.49335 |
| faces - punish | 0.0007 | 0.0007 | 11951.19 | -0.0006 | 0.0020 | 1.00 | 0.31665 |
| faces - random | 0.0016 | 0.0007 | 11931.81 | 0.0002 | 0.0029 | 2.33 | 0.01996 |
| faces - relational | 0.0003 | 0.0007 | 11931.56 | -0.0010 | 0.0017 | 0.51 | 0.61266 |
| faces - reward | 0.0002 | 0.0007 | 11951.19 | -0.0011 | 0.0015 | 0.24 | 0.81169 |
| faces - shapes | -0.0001 | 0.0007 | 11916.92 | -0.0014 | 0.0012 | -0.11 | 0.91485 |
| faces - story | 0.0013 | 0.0007 | 11950.03 | 0.0000 | 0.0027 | 1.97 | 0.04875 |
| faces - ToM | 0.0010 | 0.0007 | 11931.81 | -0.0003 | 0.0023 | 1.50 | 0.13433 |
| match - math | 0.0000 | 0.0007 | 11940.00 | -0.0014 | 0.0013 | -0.04 | 0.96995 |
| match - motor | -0.0013 | 0.0007 | 11954.75 | -0.0026 | 0.0001 | -1.85 | 0.06386 |
| match - punish | -0.0001 | 0.0007 | 11959.18 | -0.0014 | 0.0012 | -0.19 | 0.85134 |
| match - random | 0.0008 | 0.0007 | 11939.45 | -0.0006 | 0.0021 | 1.13 | 0.25769 |
| match - relational | -0.0005 | 0.0007 | 11916.92 | -0.0018 | 0.0009 | -0.67 | 0.50516 |
| match - reward | -0.0006 | 0.0007 | 11959.18 | -0.0019 | 0.0007 | -0.94 | 0.34477 |

| | | | | | | | |
|---|---|---|---|---|---|---|---|
| match - shapes | -0.0009 | 0.0007 | 11931.56 | -0.0022 | 0.0005 | -1.28 | 0.19961 |
| match - story | 0.0006 | 0.0007 | 11940.00 | -0.0008 | 0.0019 | 0.80 | 0.42148 |
| match - ToM | 0.0002 | 0.0007 | 11939.45 | -0.0011 | 0.0015 | 0.31 | 0.75807 |
| math - motor | -0.0012 | 0.0007 | 11969.74 | -0.0026 | 0.0001 | -1.79 | 0.07295 |
| math - punish | -0.0001 | 0.0007 | 11971.87 | -0.0014 | 0.0012 | -0.15 | 0.88306 |
| math - random | 0.0008 | 0.0007 | 11947.55 | -0.0005 | 0.0021 | 1.16 | 0.24745 |
| math - relational | -0.0004 | 0.0007 | 11940.00 | -0.0018 | 0.0009 | -0.62 | 0.53499 |
| math - reward | -0.0006 | 0.0007 | 11971.87 | -0.0019 | 0.0007 | -0.90 | 0.37058 |
| math - shapes | -0.0008 | 0.0007 | 11950.03 | -0.0022 | 0.0005 | -1.23 | 0.21906 |
| math - story | 0.0006 | 0.0007 | 11916.92 | -0.0008 | 0.0019 | 0.83 | 0.40437 |
| math - ToM | 0.0002 | 0.0007 | 11947.55 | -0.0011 | 0.0016 | 0.34 | 0.73207 |
| motor - punish | 0.0011 | 0.0007 | 11931.94 | -0.0002 | 0.0024 | 1.69 | 0.09107 |
| motor - random | 0.0020 | 0.0007 | 11965.26 | 0.0007 | 0.0033 | 3.00 | 0.00267 |
| motor - relational | 0.0008 | 0.0007 | 11954.75 | -0.0005 | 0.0021 | 1.19 | 0.23598 |
| motor - reward | 0.0006 | 0.0007 | 11931.94 | -0.0007 | 0.0019 | 0.93 | 0.35357 |
| motor - shapes | 0.0004 | 0.0007 | 11960.01 | -0.0009 | 0.0017 | 0.58 | 0.56289 |
| motor - story | 0.0018 | 0.0007 | 11969.74 | 0.0005 | 0.0031 | 2.64 | 0.00836 |
| motor - ToM | 0.0015 | 0.0007 | 11965.26 | 0.0001 | 0.0028 | 2.18 | 0.02946 |
| punish - random | 0.0009 | 0.0007 | 11955.97 | -0.0004 | 0.0022 | 1.33 | 0.18207 |
| punish - relational | -0.0003 | 0.0007 | 11959.18 | -0.0016 | 0.0010 | -0.49 | 0.62727 |
| punish - reward | -0.0005 | 0.0007 | 11916.92 | -0.0018 | 0.0008 | -0.77 | 0.44202 |
| punish - shapes | -0.0007 | 0.0007 | 11951.19 | -0.0020 | 0.0006 | -1.11 | 0.26757 |
| punish - story | 0.0007 | 0.0007 | 11971.87 | -0.0007 | 0.0020 | 1.00 | 0.31861 |
| punish - ToM | 0.0003 | 0.0007 | 11955.97 | -0.0010 | 0.0016 | 0.50 | 0.61652 |
| random - relational | -0.0012 | 0.0007 | 11939.45 | -0.0025 | 0.0001 | -1.80 | 0.07139 |
| random - reward | -0.0014 | 0.0007 | 11955.97 | -0.0027 | -0.0001 | -2.10 | 0.03585 |
| random - shapes | -0.0016 | 0.0007 | 11931.81 | -0.0029 | -0.0003 | -2.43 | 0.01493 |
| random - story | -0.0002 | 0.0007 | 11947.55 | -0.0015 | 0.0011 | -0.31 | 0.75782 |
| random - ToM | -0.0006 | 0.0007 | 11916.92 | -0.0019 | 0.0008 | -0.83 | 0.40539 |
| relational - reward | -0.0002 | 0.0007 | 11959.18 | -0.0015 | 0.0011 | -0.27 | 0.78576 |
| relational - shapes | -0.0004 | 0.0007 | 11931.56 | -0.0017 | 0.0009 | -0.61 | 0.54029 |
| relational - story | 0.0010 | 0.0007 | 11940.00 | -0.0003 | 0.0023 | 1.46 | 0.14377 |
| relational - ToM | 0.0007 | 0.0007 | 11939.45 | -0.0007 | 0.0020 | 0.98 | 0.32748 |
| reward - shapes | -0.0002 | 0.0007 | 11951.19 | -0.0015 | 0.0011 | -0.35 | 0.72967 |
| reward - story | 0.0012 | 0.0007 | 11971.87 | -0.0001 | 0.0025 | 1.75 | 0.08089 |
| reward - ToM | 0.0008 | 0.0007 | 11955.97 | -0.0005 | 0.0021 | 1.27 | 0.20583 |
| shapes - story | 0.0014 | 0.0007 | 11950.03 | 0.0001 | 0.0028 | 2.08 | 0.03793 |
| shapes - ToM | 0.0011 | 0.0007 | 11931.81 | -0.0002 | 0.0024 | 1.60 | 0.10866 |
| story - ToM | -0.0003 | 0.0007 | 11947.55 | -0.0017 | 0.0010 | -0.51 | 0.61295 |

**SI Table 7.4** Agreeableness* task condition interactions along D1 (primary- association)

| contrast | β | SE | df | $CL_{lower}$ | $CL_{upper}$ | t.ratio | p.value |
|---|---|---|---|---|---|---|---|
| 0back - 2back | 0.0004 | 0.0007 | 11916.92 | -0.0009 | 0.0017 | 0.56 | 0.57366 |
| 0back - faces | 0.0009 | 0.0007 | 11948.10 | -0.0004 | 0.0023 | 1.39 | 0.16415 |
| 0back - match | 0.0012 | 0.0007 | 11953.49 | -0.0001 | 0.0025 | 1.78 | 0.07558 |
| 0back - math | -0.0007 | 0.0007 | 11978.23 | -0.0020 | 0.0007 | -0.98 | 0.32575 |
| 0back - motor | 0.0003 | 0.0007 | 11930.91 | -0.0010 | 0.0016 | 0.44 | 0.66224 |
| 0back - punish | 0.0006 | 0.0007 | 11929.93 | -0.0007 | 0.0019 | 0.88 | 0.37864 |
| 0back - random | -0.0008 | 0.0007 | 11960.92 | -0.0022 | 0.0005 | -1.26 | 0.20636 |
| 0back - relational | 0.0012 | 0.0007 | 11953.49 | -0.0001 | 0.0026 | 1.82 | 0.06929 |
| 0back - reward | 0.0002 | 0.0007 | 11929.93 | -0.0011 | 0.0015 | 0.32 | 0.75102 |
| 0back - shapes | 0.0011 | 0.0007 | 11948.10 | -0.0002 | 0.0025 | 1.71 | 0.08771 |
| 0back - story | -0.0007 | 0.0007 | 11978.23 | -0.0021 | 0.0006 | -1.04 | 0.29816 |
| 0back - ToM | 0.0000 | 0.0007 | 11960.92 | -0.0013 | 0.0013 | 0.03 | 0.97449 |
| 2back - faces | 0.0006 | 0.0007 | 11948.10 | -0.0008 | 0.0019 | 0.83 | 0.40494 |
| 2back - match | 0.0008 | 0.0007 | 11953.49 | -0.0005 | 0.0022 | 1.22 | 0.22194 |
| 2back - math | -0.0011 | 0.0007 | 11978.23 | -0.0024 | 0.0003 | -1.52 | 0.12732 |
| 2back - motor | -0.0001 | 0.0007 | 11930.91 | -0.0014 | 0.0012 | -0.12 | 0.90276 |
| 2back - punish | 0.0002 | 0.0007 | 11929.93 | -0.0011 | 0.0015 | 0.32 | 0.75020 |
| 2back - random | -0.0012 | 0.0007 | 11960.92 | -0.0025 | 0.0001 | -1.82 | 0.06818 |
| 2back - relational | 0.0009 | 0.0007 | 11953.49 | -0.0005 | 0.0022 | 1.26 | 0.20731 |
| 2back - reward | -0.0002 | 0.0007 | 11929.93 | -0.0015 | 0.0011 | -0.24 | 0.80665 |
| 2back - shapes | 0.0008 | 0.0007 | 11948.10 | -0.0005 | 0.0021 | 1.15 | 0.25048 |
| 2back - story | -0.0011 | 0.0007 | 11978.23 | -0.0025 | 0.0003 | -1.58 | 0.11355 |
| 2back - ToM | -0.0004 | 0.0007 | 11960.92 | -0.0017 | 0.0010 | -0.53 | 0.59732 |
| faces - match | 0.0003 | 0.0007 | 11928.40 | -0.0011 | 0.0016 | 0.39 | 0.69574 |
| faces - math | -0.0016 | 0.0007 | 11957.50 | -0.0030 | -0.0003 | -2.32 | 0.02021 |
| faces - motor | -0.0006 | 0.0007 | 11954.54 | -0.0020 | 0.0007 | -0.95 | 0.34255 |
| faces - punish | -0.0003 | 0.0007 | 11945.55 | -0.0017 | 0.0010 | -0.52 | 0.60560 |
| faces - random | -0.0018 | 0.0007 | 11941.96 | -0.0031 | -0.0005 | -2.65 | 0.00815 |
| faces - relational | 0.0003 | 0.0007 | 11928.40 | -0.0010 | 0.0016 | 0.43 | 0.66681 |
| faces - reward | -0.0007 | 0.0007 | 11945.55 | -0.0020 | 0.0006 | -1.08 | 0.28205 |
| faces - shapes | 0.0002 | 0.0007 | 11916.92 | -0.0011 | 0.0015 | 0.32 | 0.75272 |
| faces - story | -0.0017 | 0.0007 | 11957.50 | -0.0030 | -0.0003 | -2.38 | 0.01732 |
| faces - ToM | -0.0009 | 0.0007 | 11941.96 | -0.0022 | 0.0004 | -1.36 | 0.17479 |
| match - math | -0.0019 | 0.0007 | 11953.39 | -0.0033 | -0.0005 | -2.69 | 0.00710 |
| match - motor | -0.0009 | 0.0007 | 11949.67 | -0.0022 | 0.0004 | -1.34 | 0.18170 |
| match - punish | -0.0006 | 0.0007 | 11951.47 | -0.0019 | 0.0007 | -0.91 | 0.36476 |
| match - random | -0.0020 | 0.0007 | 11950.08 | -0.0034 | -0.0007 | -3.02 | 0.00250 |
| match - relational | 0.0000 | 0.0007 | 11916.92 | -0.0013 | 0.0014 | 0.04 | 0.96866 |
| match - reward | -0.0010 | 0.0007 | 11951.47 | -0.0023 | 0.0003 | -1.46 | 0.14354 |

| | | | | | | | |
|---|---|---|---|---|---|---|---|
| match - shapes | -0.0001 | 0.0007 | 11928.40 | -0.0014 | 0.0013 | -0.08 | 0.93804 |
| match - story | -0.0019 | 0.0007 | 11953.39 | -0.0033 | -0.0006 | -2.75 | 0.00597 |
| match - ToM | -0.0012 | 0.0007 | 11950.08 | -0.0025 | 0.0001 | -1.74 | 0.08156 |
| math - motor | 0.0010 | 0.0007 | 11975.70 | -0.0004 | 0.0023 | 1.40 | 0.16182 |
| math - punish | 0.0013 | 0.0007 | 11973.24 | -0.0001 | 0.0026 | 1.83 | 0.06703 |
| math - random | -0.0002 | 0.0007 | 11965.38 | -0.0015 | 0.0012 | -0.24 | 0.81059 |
| math - relational | 0.0019 | 0.0007 | 11953.39 | 0.0005 | 0.0033 | 2.73 | 0.00632 |
| math - reward | 0.0009 | 0.0007 | 11973.24 | -0.0005 | 0.0023 | 1.29 | 0.19756 |
| math - shapes | 0.0018 | 0.0007 | 11957.50 | 0.0005 | 0.0032 | 2.63 | 0.00859 |
| math - story | 0.0000 | 0.0007 | 11916.92 | -0.0014 | 0.0014 | -0.06 | 0.95526 |
| math - ToM | 0.0007 | 0.0007 | 11965.38 | -0.0007 | 0.0021 | 1.01 | 0.31141 |
| motor - punish | 0.0003 | 0.0007 | 11933.07 | -0.0010 | 0.0016 | 0.44 | 0.66095 |
| motor - random | -0.0011 | 0.0007 | 11970.77 | -0.0025 | 0.0002 | -1.69 | 0.09090 |
| motor - relational | 0.0009 | 0.0007 | 11949.67 | -0.0004 | 0.0023 | 1.38 | 0.16914 |
| motor - reward | -0.0001 | 0.0007 | 11933.07 | -0.0014 | 0.0012 | -0.12 | 0.90354 |
| motor - shapes | 0.0009 | 0.0007 | 11954.54 | -0.0005 | 0.0022 | 1.26 | 0.20638 |
| motor - story | -0.0010 | 0.0007 | 11975.70 | -0.0024 | 0.0004 | -1.46 | 0.14529 |
| motor - ToM | -0.0003 | 0.0007 | 11970.77 | -0.0016 | 0.0011 | -0.40 | 0.68676 |
| punish - random | -0.0014 | 0.0007 | 11959.73 | -0.0028 | -0.0001 | -2.14 | 0.03233 |
| punish - relational | 0.0006 | 0.0007 | 11951.47 | -0.0007 | 0.0020 | 0.95 | 0.34418 |
| punish - reward | -0.0004 | 0.0007 | 11916.92 | -0.0017 | 0.0009 | -0.56 | 0.57325 |
| punish - shapes | 0.0006 | 0.0007 | 11945.55 | -0.0008 | 0.0019 | 0.83 | 0.40504 |
| punish - story | -0.0013 | 0.0007 | 11973.24 | -0.0027 | 0.0000 | -1.89 | 0.05887 |
| punish - ToM | -0.0006 | 0.0007 | 11959.73 | -0.0019 | 0.0007 | -0.85 | 0.39795 |
| random - relational | 0.0021 | 0.0007 | 11950.08 | 0.0007 | 0.0034 | 3.06 | 0.00219 |
| random - reward | 0.0011 | 0.0007 | 11959.73 | -0.0003 | 0.0024 | 1.58 | 0.11427 |
| random - shapes | 0.0020 | 0.0007 | 11941.96 | 0.0007 | 0.0033 | 2.96 | 0.00306 |
| random - story | 0.0001 | 0.0007 | 11965.38 | -0.0012 | 0.0015 | 0.18 | 0.85553 |
| random - ToM | 0.0009 | 0.0007 | 11916.92 | -0.0004 | 0.0022 | 1.30 | 0.19527 |
| relational - reward | -0.0010 | 0.0007 | 11951.47 | -0.0023 | 0.0003 | -1.50 | 0.13301 |
| relational - shapes | -0.0001 | 0.0007 | 11928.40 | -0.0014 | 0.0013 | -0.12 | 0.90672 |
| relational - story | -0.0020 | 0.0007 | 11953.39 | -0.0033 | -0.0006 | -2.79 | 0.00531 |
| relational - ToM | -0.0012 | 0.0007 | 11950.08 | -0.0025 | 0.0001 | -1.78 | 0.07487 |
| reward - shapes | 0.0009 | 0.0007 | 11945.55 | -0.0004 | 0.0023 | 1.39 | 0.16392 |
| reward - story | -0.0009 | 0.0007 | 11973.24 | -0.0023 | 0.0004 | -1.35 | 0.17824 |
| reward - ToM | -0.0002 | 0.0007 | 11959.73 | -0.0015 | 0.0011 | -0.28 | 0.77627 |
| shapes - story | -0.0019 | 0.0007 | 11957.50 | -0.0032 | -0.0005 | -2.69 | 0.00724 |
| shapes - ToM | -0.0011 | 0.0007 | 11941.96 | -0.0024 | 0.0002 | -1.67 | 0.09437 |
| story - ToM | 0.0007 | 0.0007 | 11965.38 | -0.0006 | 0.0021 | 1.07 | 0.28468 |

**SI Table 8.1** Neuroticism* task condition interactions along D2 (motor- visual)

| **contrast** | **β** | **SE** | **df** | **$CL_{lower}$** | **$CL_{upper}$** | **t.ratio** | **p.value** |
|---|---|---|---|---|---|---|---|
| 0back - 2back | 0.0009 | 0.0004 | 11922.26 | 0.0001 | 0.0017 | 2.22 | 0.02632 |
| 0back - faces | 0.0003 | 0.0004 | 11971.88 | -0.0005 | 0.0012 | 0.82 | 0.41306 |
| 0back - match | 0.0000 | 0.0004 | 11975.31 | -0.0008 | 0.0008 | -0.03 | 0.97897 |
| 0back - math | -0.0006 | 0.0004 | 11983.92 | -0.0015 | 0.0002 | -1.47 | 0.14239 |
| 0back - motor | -0.0010 | 0.0004 | 11940.41 | -0.0018 | -0.0002 | -2.34 | 0.01921 |
| 0back - punish | -0.0001 | 0.0004 | 11943.31 | -0.0009 | 0.0007 | -0.15 | 0.87940 |
| 0back - random | -0.0004 | 0.0004 | 11978.05 | -0.0012 | 0.0004 | -0.97 | 0.33209 |
| 0back - relational | 0.0001 | 0.0004 | 11975.31 | -0.0007 | 0.0010 | 0.33 | 0.73810 |
| 0back - reward | -0.0001 | 0.0004 | 11943.31 | -0.0009 | 0.0007 | -0.17 | 0.86735 |
| 0back - shapes | 0.0001 | 0.0004 | 11971.88 | -0.0007 | 0.0009 | 0.19 | 0.85175 |
| 0back - story | -0.0008 | 0.0004 | 11983.92 | -0.0016 | 0.0000 | -1.85 | 0.06467 |
| 0back - ToM | -0.0002 | 0.0004 | 11978.05 | -0.0010 | 0.0006 | -0.55 | 0.57908 |
| 2back - faces | -0.0006 | 0.0004 | 11971.89 | -0.0014 | 0.0002 | -1.38 | 0.16781 |
| 2back - match | -0.0009 | 0.0004 | 11975.32 | -0.0018 | -0.0001 | -2.21 | 0.02690 |
| 2back - math | -0.0015 | 0.0004 | 11983.93 | -0.0024 | -0.0007 | -3.62 | 0.00030 |
| 2back - motor | -0.0019 | 0.0004 | 11940.42 | -0.0027 | -0.0011 | -4.55 | 0.00001 |
| 2back - punish | -0.0010 | 0.0004 | 11943.32 | -0.0018 | -0.0002 | -2.37 | 0.01772 |
| 2back - random | -0.0013 | 0.0004 | 11978.06 | -0.0021 | -0.0005 | -3.17 | 0.00151 |
| 2back - relational | -0.0008 | 0.0004 | 11975.32 | -0.0016 | 0.0000 | -1.85 | 0.06398 |
| 2back - reward | -0.0010 | 0.0004 | 11943.32 | -0.0018 | -0.0002 | -2.39 | 0.01700 |
| 2back - shapes | -0.0008 | 0.0004 | 11971.89 | -0.0017 | 0.0000 | -2.01 | 0.04435 |
| 2back - story | -0.0017 | 0.0004 | 11983.93 | -0.0025 | -0.0009 | -4.00 | 0.00006 |
| 2back - ToM | -0.0011 | 0.0004 | 11978.06 | -0.0020 | -0.0003 | -2.76 | 0.00581 |
| faces - match | -0.0004 | 0.0004 | 11934.81 | -0.0012 | 0.0005 | -0.84 | 0.40362 |
| faces - math | -0.0010 | 0.0004 | 11955.45 | -0.0018 | -0.0001 | -2.25 | 0.02429 |
| faces - motor | -0.0013 | 0.0004 | 11979.99 | -0.0021 | -0.0005 | -3.13 | 0.00174 |
| faces - punish | -0.0004 | 0.0004 | 11970.40 | -0.0012 | 0.0004 | -0.97 | 0.33256 |
| faces - random | -0.0007 | 0.0004 | 11938.33 | -0.0016 | 0.0001 | -1.78 | 0.07534 |
| faces - relational | -0.0002 | 0.0004 | 11934.81 | -0.0010 | 0.0006 | -0.48 | 0.63347 |
| faces - reward | -0.0004 | 0.0004 | 11970.40 | -0.0012 | 0.0004 | -0.98 | 0.32506 |
| faces - shapes | -0.0003 | 0.0004 | 11922.26 | -0.0011 | 0.0006 | -0.63 | 0.53021 |
| faces - story | -0.0011 | 0.0004 | 11955.45 | -0.0020 | -0.0003 | -2.63 | 0.00853 |
| faces - ToM | -0.0006 | 0.0004 | 11938.33 | -0.0014 | 0.0002 | -1.37 | 0.17191 |
| match - math | -0.0006 | 0.0004 | 11949.32 | -0.0015 | 0.0002 | -1.42 | 0.15425 |
| match - motor | -0.0010 | 0.0004 | 11976.97 | -0.0018 | -0.0001 | -2.28 | 0.02258 |
| match - punish | -0.0001 | 0.0004 | 11978.71 | -0.0009 | 0.0008 | -0.12 | 0.90202 |
| match - random | -0.0004 | 0.0004 | 11945.62 | -0.0012 | 0.0004 | -0.93 | 0.35134 |
| match - relational | 0.0002 | 0.0004 | 11922.26 | -0.0007 | 0.0010 | 0.36 | 0.72119 |
| match - reward | -0.0001 | 0.0004 | 11978.71 | -0.0009 | 0.0008 | -0.14 | 0.89011 |

| | | | | | | | |
|---|---|---|---|---|---|---|---|
| match - shapes | 0.0001 | 0.0004 | 11934.81 | -0.0007 | 0.0009 | 0.21 | 0.83297 |
| match - story | -0.0008 | 0.0004 | 11949.32 | -0.0016 | 0.0001 | -1.80 | 0.07170 |
| match - ToM | -0.0002 | 0.0004 | 11945.62 | -0.0010 | 0.0006 | -0.52 | 0.60181 |
| math - motor | -0.0003 | 0.0004 | 11988.96 | -0.0012 | 0.0005 | -0.81 | 0.41796 |
| math - punish | 0.0006 | 0.0004 | 11991.05 | -0.0003 | 0.0014 | 1.32 | 0.18700 |
| math - random | 0.0002 | 0.0004 | 11960.60 | -0.0006 | 0.0011 | 0.52 | 0.60483 |
| math - relational | 0.0008 | 0.0004 | 11949.32 | -0.0001 | 0.0016 | 1.78 | 0.07578 |
| math - reward | 0.0006 | 0.0004 | 11991.05 | -0.0003 | 0.0014 | 1.30 | 0.19200 |
| math - shapes | 0.0007 | 0.0004 | 11955.45 | -0.0001 | 0.0015 | 1.64 | 0.10134 |
| math - story | -0.0002 | 0.0004 | 11922.26 | -0.0010 | 0.0007 | -0.37 | 0.71011 |
| math - ToM | 0.0004 | 0.0004 | 11960.60 | -0.0004 | 0.0012 | 0.92 | 0.35696 |
| motor - punish | 0.0009 | 0.0004 | 11943.13 | 0.0001 | 0.0017 | 2.19 | 0.02850 |
| motor - random | 0.0006 | 0.0004 | 11987.87 | -0.0003 | 0.0014 | 1.36 | 0.17427 |
| motor - relational | 0.0011 | 0.0004 | 11976.97 | 0.0003 | 0.0019 | 2.64 | 0.00830 |
| motor - reward | 0.0009 | 0.0004 | 11943.13 | 0.0001 | 0.0017 | 2.18 | 0.02962 |
| motor - shapes | 0.0011 | 0.0004 | 11979.99 | 0.0002 | 0.0019 | 2.50 | 0.01230 |
| motor - story | 0.0002 | 0.0004 | 11988.96 | -0.0007 | 0.0010 | 0.43 | 0.66654 |
| motor - ToM | 0.0007 | 0.0004 | 11987.87 | -0.0001 | 0.0016 | 1.77 | 0.07643 |
| punish - random | -0.0003 | 0.0004 | 11976.70 | -0.0012 | 0.0005 | -0.82 | 0.41256 |
| punish - relational | 0.0002 | 0.0004 | 11978.71 | -0.0006 | 0.0010 | 0.48 | 0.62851 |
| punish - reward | 0.0000 | 0.0004 | 11922.26 | -0.0008 | 0.0008 | -0.02 | 0.98779 |
| punish - shapes | 0.0001 | 0.0004 | 11970.40 | -0.0007 | 0.0010 | 0.34 | 0.73597 |
| punish - story | -0.0007 | 0.0004 | 11991.05 | -0.0016 | 0.0001 | -1.70 | 0.08912 |
| punish - ToM | -0.0002 | 0.0004 | 11976.70 | -0.0010 | 0.0006 | -0.40 | 0.68610 |
| random - relational | 0.0005 | 0.0004 | 11945.62 | -0.0003 | 0.0014 | 1.29 | 0.19665 |
| random - reward | 0.0003 | 0.0004 | 11976.70 | -0.0005 | 0.0012 | 0.80 | 0.42127 |
| random - shapes | 0.0005 | 0.0004 | 11938.33 | -0.0003 | 0.0013 | 1.15 | 0.25040 |
| random - story | -0.0004 | 0.0004 | 11960.60 | -0.0012 | 0.0005 | -0.90 | 0.37000 |
| random - ToM | 0.0002 | 0.0004 | 11922.26 | -0.0006 | 0.0010 | 0.41 | 0.67903 |
| relational - reward | -0.0002 | 0.0004 | 11978.71 | -0.0010 | 0.0006 | -0.50 | 0.61786 |
| relational - shapes | -0.0001 | 0.0004 | 11934.81 | -0.0009 | 0.0008 | -0.15 | 0.88280 |
| relational - story | -0.0009 | 0.0004 | 11949.32 | -0.0018 | -0.0001 | -2.15 | 0.03140 |
| relational - ToM | -0.0004 | 0.0004 | 11945.62 | -0.0012 | 0.0005 | -0.88 | 0.37833 |
| reward - shapes | 0.0001 | 0.0004 | 11970.40 | -0.0007 | 0.0010 | 0.35 | 0.72458 |
| reward - story | -0.0007 | 0.0004 | 11991.05 | -0.0016 | 0.0001 | -1.69 | 0.09195 |
| reward - ToM | -0.0002 | 0.0004 | 11976.70 | -0.0010 | 0.0007 | -0.39 | 0.69730 |
| shapes - story | -0.0009 | 0.0004 | 11955.45 | -0.0017 | 0.0000 | -2.02 | 0.04377 |
| shapes - ToM | -0.0003 | 0.0004 | 11938.33 | -0.0011 | 0.0005 | -0.74 | 0.46109 |
| story - ToM | 0.0006 | 0.0004 | 11960.60 | -0.0003 | 0.0014 | 1.30 | 0.19355 |

**SI Table 8.2** Openness* task condition interactions along D2 (motor- visual)

| contrast | β | SE | df | CL lower | CL upper | t.ratio | p.value |
|---|---|---|---|---|---|---|---|
| 0back - 2back | -0.0005 | 0.0004 | 11922.26 | -0.0014 | 0.0003 | -1.20 | 0.23013 |
| 0back - faces | -0.0005 | 0.0004 | 11968.02 | -0.0014 | 0.0004 | -1.11 | 0.26817 |
| 0back - match | 0.0006 | 0.0004 | 11972.91 | -0.0003 | 0.0014 | 1.24 | 0.21323 |
| 0back - math | -0.0011 | 0.0005 | 11987.09 | -0.0019 | -0.0002 | -2.32 | 0.02041 |
| 0back - motor | 0.0000 | 0.0004 | 11942.86 | -0.0009 | 0.0008 | -0.09 | 0.92739 |
| 0back - punish | 0.0006 | 0.0004 | 11940.58 | -0.0003 | 0.0014 | 1.35 | 0.17768 |
| 0back - random | -0.0006 | 0.0004 | 11969.08 | -0.0014 | 0.0003 | -1.28 | 0.19983 |
| 0back - relational | 0.0000 | 0.0004 | 11972.91 | -0.0009 | 0.0008 | -0.11 | 0.91392 |
| 0back - reward | 0.0004 | 0.0004 | 11940.58 | -0.0005 | 0.0012 | 0.88 | 0.38049 |
| 0back - shapes | 0.0003 | 0.0004 | 11968.02 | -0.0006 | 0.0012 | 0.69 | 0.49277 |
| 0back - story | -0.0006 | 0.0005 | 11987.09 | -0.0015 | 0.0003 | -1.35 | 0.17814 |
| 0back - ToM | -0.0001 | 0.0004 | 11969.08 | -0.0009 | 0.0008 | -0.12 | 0.90412 |
| 2back - faces | 0.0000 | 0.0004 | 11968.02 | -0.0008 | 0.0009 | 0.08 | 0.93651 |
| 2back - match | 0.0011 | 0.0004 | 11972.92 | 0.0002 | 0.0020 | 2.43 | 0.01532 |
| 2back - math | -0.0005 | 0.0005 | 11987.09 | -0.0014 | 0.0004 | -1.16 | 0.24605 |
| 2back - motor | 0.0005 | 0.0004 | 11942.87 | -0.0004 | 0.0013 | 1.10 | 0.27091 |
| 2back - punish | 0.0011 | 0.0004 | 11940.59 | 0.0003 | 0.0020 | 2.55 | 0.01077 |
| 2back - random | 0.0000 | 0.0004 | 11969.08 | -0.0009 | 0.0008 | -0.09 | 0.92713 |
| 2back - relational | 0.0005 | 0.0004 | 11972.92 | -0.0004 | 0.0014 | 1.07 | 0.28364 |
| 2back - reward | 0.0009 | 0.0004 | 11940.59 | 0.0001 | 0.0018 | 2.08 | 0.03758 |
| 2back - shapes | 0.0008 | 0.0004 | 11968.02 | 0.0000 | 0.0017 | 1.87 | 0.06110 |
| 2back - story | -0.0001 | 0.0005 | 11987.09 | -0.0010 | 0.0008 | -0.19 | 0.85120 |
| 2back - ToM | 0.0005 | 0.0004 | 11969.08 | -0.0004 | 0.0013 | 1.07 | 0.28456 |
| faces - match | 0.0010 | 0.0004 | 11942.02 | 0.0002 | 0.0019 | 2.33 | 0.01987 |
| faces - math | -0.0006 | 0.0005 | 11963.12 | -0.0015 | 0.0003 | -1.23 | 0.21911 |
| faces - motor | 0.0005 | 0.0004 | 11978.21 | -0.0004 | 0.0013 | 1.01 | 0.31222 |
| faces - punish | 0.0011 | 0.0004 | 11968.99 | 0.0002 | 0.0019 | 2.45 | 0.01449 |
| faces - random | -0.0001 | 0.0004 | 11933.03 | -0.0009 | 0.0008 | -0.17 | 0.86490 |
| faces - relational | 0.0004 | 0.0004 | 11942.02 | -0.0004 | 0.0013 | 0.99 | 0.32419 |
| faces - reward | 0.0009 | 0.0004 | 11968.99 | 0.0000 | 0.0017 | 1.98 | 0.04785 |
| faces - shapes | 0.0008 | 0.0004 | 11922.26 | -0.0001 | 0.0017 | 1.78 | 0.07483 |
| faces - story | -0.0001 | 0.0005 | 11963.12 | -0.0010 | 0.0008 | -0.26 | 0.79220 |
| faces - ToM | 0.0004 | 0.0004 | 11933.03 | -0.0004 | 0.0013 | 0.98 | 0.32550 |
| match - math | -0.0016 | 0.0005 | 11952.08 | -0.0025 | -0.0007 | -3.50 | 0.00047 |
| match - motor | -0.0006 | 0.0004 | 11975.19 | -0.0015 | 0.0003 | -1.33 | 0.18428 |
| match - punish | 0.0000 | 0.0004 | 11980.90 | -0.0008 | 0.0009 | 0.08 | 0.93746 |
| match - random | -0.0011 | 0.0004 | 11946.66 | -0.0020 | -0.0002 | -2.50 | 0.01227 |
| match - relational | -0.0006 | 0.0005 | 11922.26 | -0.0015 | 0.0003 | -1.34 | 0.18106 |
| match - reward | -0.0002 | 0.0004 | 11980.90 | -0.0010 | 0.0007 | -0.39 | 0.70024 |

| | | | | | | | |
|---|---|---|---|---|---|---|---|
| match - shapes | -0.0003 | 0.0004 | 11942.02 | -0.0011 | 0.0006 | -0.56 | 0.57638 |
| match - story | -0.0012 | 0.0005 | 11952.08 | -0.0021 | -0.0003 | -2.54 | 0.01113 |
| match - ToM | -0.0006 | 0.0004 | 11946.66 | -0.0015 | 0.0003 | -1.36 | 0.17444 |
| math - motor | 0.0010 | 0.0005 | 11986.21 | 0.0001 | 0.0019 | 2.22 | 0.02650 |
| math - punish | 0.0016 | 0.0005 | 11993.86 | 0.0008 | 0.0025 | 3.63 | 0.00029 |
| math - random | 0.0005 | 0.0005 | 11961.42 | -0.0004 | 0.0014 | 1.07 | 0.28622 |
| math - relational | 0.0010 | 0.0005 | 11952.08 | 0.0001 | 0.0019 | 2.19 | 0.02875 |
| math - reward | 0.0014 | 0.0005 | 11993.86 | 0.0005 | 0.0023 | 3.17 | 0.00152 |
| math - shapes | 0.0014 | 0.0005 | 11963.12 | 0.0005 | 0.0023 | 2.97 | 0.00301 |
| math - story | 0.0004 | 0.0005 | 11922.26 | -0.0005 | 0.0014 | 0.95 | 0.34337 |
| math - ToM | 0.0010 | 0.0005 | 11961.42 | 0.0001 | 0.0019 | 2.19 | 0.02836 |
| motor - punish | 0.0006 | 0.0004 | 11943.46 | -0.0002 | 0.0015 | 1.43 | 0.15229 |
| motor - random | -0.0005 | 0.0004 | 11977.18 | -0.0014 | 0.0003 | -1.18 | 0.23640 |
| motor - relational | 0.0000 | 0.0004 | 11975.19 | -0.0009 | 0.0009 | -0.02 | 0.98582 |
| motor - reward | 0.0004 | 0.0004 | 11943.46 | -0.0004 | 0.0013 | 0.96 | 0.33541 |
| motor - shapes | 0.0003 | 0.0004 | 11978.21 | -0.0005 | 0.0012 | 0.77 | 0.44008 |
| motor - story | -0.0006 | 0.0005 | 11986.21 | -0.0015 | 0.0003 | -1.25 | 0.21074 |
| motor - ToM | 0.0000 | 0.0004 | 11977.18 | -0.0009 | 0.0009 | -0.03 | 0.97665 |
| punish - random | -0.0012 | 0.0004 | 11964.15 | -0.0020 | -0.0003 | -2.63 | 0.00868 |
| punish - relational | -0.0006 | 0.0004 | 11980.90 | -0.0015 | 0.0002 | -1.43 | 0.15135 |
| punish - reward | -0.0002 | 0.0004 | 11922.26 | -0.0011 | 0.0006 | -0.47 | 0.63639 |
| punish - shapes | -0.0003 | 0.0004 | 11968.99 | -0.0012 | 0.0006 | -0.65 | 0.51794 |
| punish - story | -0.0012 | 0.0005 | 11993.86 | -0.0021 | -0.0003 | -2.65 | 0.00800 |
| punish - ToM | -0.0006 | 0.0004 | 11964.15 | -0.0015 | 0.0002 | -1.46 | 0.14440 |
| random - relational | 0.0005 | 0.0004 | 11946.66 | -0.0004 | 0.0014 | 1.16 | 0.24700 |
| random - reward | 0.0009 | 0.0004 | 11964.15 | 0.0001 | 0.0018 | 2.16 | 0.03101 |
| random - shapes | 0.0009 | 0.0004 | 11933.03 | 0.0000 | 0.0017 | 1.96 | 0.05047 |
| random - story | 0.0000 | 0.0005 | 11961.42 | -0.0009 | 0.0008 | -0.10 | 0.92181 |
| random - ToM | 0.0005 | 0.0004 | 11922.26 | -0.0004 | 0.0014 | 1.16 | 0.24712 |
| relational - reward | 0.0004 | 0.0004 | 11980.90 | -0.0004 | 0.0013 | 0.97 | 0.33136 |
| relational - shapes | 0.0004 | 0.0004 | 11942.02 | -0.0005 | 0.0012 | 0.78 | 0.43275 |
| relational - story | -0.0006 | 0.0005 | 11952.08 | -0.0015 | 0.0003 | -1.23 | 0.22023 |
| relational - ToM | 0.0000 | 0.0004 | 11946.66 | -0.0009 | 0.0009 | -0.01 | 0.99101 |
| reward - shapes | -0.0001 | 0.0004 | 11968.99 | -0.0009 | 0.0008 | -0.18 | 0.85695 |
| reward - story | -0.0010 | 0.0005 | 11993.86 | -0.0019 | -0.0001 | -2.20 | 0.02802 |
| reward - ToM | -0.0004 | 0.0004 | 11964.15 | -0.0013 | 0.0004 | -0.99 | 0.32126 |
| shapes - story | -0.0009 | 0.0005 | 11963.12 | -0.0018 | 0.0000 | -2.00 | 0.04532 |
| shapes - ToM | -0.0004 | 0.0004 | 11933.03 | -0.0012 | 0.0005 | -0.80 | 0.42214 |
| story - ToM | 0.0006 | 0.0005 | 11961.42 | -0.0003 | 0.0015 | 1.22 | 0.22093 |

**SI Table 8.3** Conscientiousness* task condition interactions along D2 (motor- visual)

| contrast | β | SE | df | CL lower | CL upper | t.ratio | p.value |
|---|---|---|---|---|---|---|---|
| 0back - 2back | 0.0006 | 0.0005 | 11922.26 | -0.0004 | 0.0015 | 1.13 | 0.25828 |
| 0back - faces | 0.0002 | 0.0005 | 11980.41 | -0.0007 | 0.0012 | 0.49 | 0.62476 |
| 0back - match | -0.0007 | 0.0005 | 11984.50 | -0.0017 | 0.0003 | -1.35 | 0.17580 |
| 0back - math | -0.0001 | 0.0005 | 12000.76 | -0.0011 | 0.0009 | -0.16 | 0.87072 |
| 0back - motor | -0.0004 | 0.0005 | 11936.32 | -0.0014 | 0.0006 | -0.78 | 0.43314 |
| 0back - punish | -0.0002 | 0.0005 | 11943.83 | -0.0012 | 0.0007 | -0.46 | 0.64431 |
| 0back - random | -0.0008 | 0.0005 | 11987.75 | -0.0018 | 0.0002 | -1.59 | 0.11190 |
| 0back - relational | -0.0003 | 0.0005 | 11984.50 | -0.0013 | 0.0007 | -0.52 | 0.60520 |
| 0back - reward | -0.0002 | 0.0005 | 11943.83 | -0.0012 | 0.0008 | -0.44 | 0.65651 |
| 0back - shapes | 0.0000 | 0.0005 | 11980.41 | -0.0010 | 0.0010 | 0.01 | 0.99086 |
| 0back - story | -0.0007 | 0.0005 | 12000.76 | -0.0018 | 0.0003 | -1.44 | 0.14909 |
| 0back - ToM | -0.0005 | 0.0005 | 11987.75 | -0.0015 | 0.0005 | -0.90 | 0.36782 |
| 2back - faces | -0.0003 | 0.0005 | 11980.42 | -0.0013 | 0.0007 | -0.63 | 0.52922 |
| 2back - match | -0.0013 | 0.0005 | 11984.51 | -0.0023 | -0.0003 | -2.47 | 0.01369 |
| 2back - math | -0.0007 | 0.0005 | 12000.77 | -0.0017 | 0.0004 | -1.26 | 0.20890 |
| 2back - motor | -0.0010 | 0.0005 | 11936.32 | -0.0020 | 0.0000 | -1.91 | 0.05647 |
| 2back - punish | -0.0008 | 0.0005 | 11943.84 | -0.0018 | 0.0002 | -1.59 | 0.11083 |
| 2back - random | -0.0014 | 0.0005 | 11987.76 | -0.0024 | -0.0004 | -2.71 | 0.00678 |
| 2back - relational | -0.0008 | 0.0005 | 11984.51 | -0.0018 | 0.0002 | -1.63 | 0.10337 |
| 2back - reward | -0.0008 | 0.0005 | 11943.84 | -0.0018 | 0.0002 | -1.58 | 0.11468 |
| 2back - shapes | -0.0006 | 0.0005 | 11980.42 | -0.0016 | 0.0004 | -1.11 | 0.26836 |
| 2back - story | -0.0013 | 0.0005 | 12000.77 | -0.0023 | -0.0003 | -2.54 | 0.01120 |
| 2back - ToM | -0.0010 | 0.0005 | 11987.76 | -0.0020 | 0.0000 | -2.02 | 0.04356 |
| faces - match | -0.0009 | 0.0005 | 11934.36 | -0.0019 | 0.0001 | -1.83 | 0.06734 |
| faces - math | -0.0003 | 0.0005 | 11957.43 | -0.0014 | 0.0007 | -0.64 | 0.52385 |
| faces - motor | -0.0006 | 0.0005 | 11986.40 | -0.0016 | 0.0004 | -1.26 | 0.20686 |
| faces - punish | -0.0005 | 0.0005 | 11975.53 | -0.0015 | 0.0005 | -0.95 | 0.34314 |
| faces - random | -0.0011 | 0.0005 | 11939.47 | -0.0021 | -0.0001 | -2.07 | 0.03881 |
| faces - relational | -0.0005 | 0.0005 | 11934.36 | -0.0015 | 0.0005 | -1.00 | 0.31856 |
| faces - reward | -0.0005 | 0.0005 | 11975.53 | -0.0015 | 0.0005 | -0.93 | 0.35175 |
| faces - shapes | -0.0002 | 0.0005 | 11922.26 | -0.0012 | 0.0008 | -0.48 | 0.63477 |
| faces - story | -0.0010 | 0.0005 | 11957.43 | -0.0020 | 0.0000 | -1.91 | 0.05615 |
| faces - ToM | -0.0007 | 0.0005 | 11939.47 | -0.0017 | 0.0003 | -1.38 | 0.16723 |
| match - math | 0.0006 | 0.0005 | 11955.54 | -0.0004 | 0.0016 | 1.16 | 0.24752 |
| match - motor | 0.0003 | 0.0005 | 11983.22 | -0.0007 | 0.0013 | 0.58 | 0.56508 |
| match - punish | 0.0005 | 0.0005 | 11982.11 | -0.0005 | 0.0015 | 0.90 | 0.36626 |
| match - random | -0.0001 | 0.0005 | 11948.16 | -0.0011 | 0.0009 | -0.23 | 0.82127 |
| match - relational | 0.0004 | 0.0005 | 11922.26 | -0.0006 | 0.0014 | 0.83 | 0.40759 |
| match - reward | 0.0005 | 0.0005 | 11982.11 | -0.0005 | 0.0015 | 0.92 | 0.35748 |

| | | | | | | | |
|---|---|---|---|---|---|---|---|
| match - shapes | 0.0007 | 0.0005 | 11934.36 | -0.0003 | 0.0017 | 1.36 | 0.17469 |
| match - story | -0.0001 | 0.0005 | 11955.54 | -0.0011 | 0.0010 | -0.11 | 0.91265 |
| match - ToM | 0.0002 | 0.0005 | 11948.16 | -0.0008 | 0.0012 | 0.46 | 0.64899 |
| math - motor | -0.0003 | 0.0005 | 11992.67 | -0.0013 | 0.0007 | -0.60 | 0.55006 |
| math - punish | -0.0001 | 0.0005 | 11997.02 | -0.0012 | 0.0009 | -0.28 | 0.77646 |
| math - random | -0.0007 | 0.0005 | 11966.26 | -0.0017 | 0.0003 | -1.38 | 0.16635 |
| math - relational | -0.0002 | 0.0005 | 11955.54 | -0.0012 | 0.0008 | -0.34 | 0.73232 |
| math - reward | -0.0001 | 0.0005 | 11997.02 | -0.0012 | 0.0009 | -0.27 | 0.78907 |
| math - shapes | 0.0001 | 0.0005 | 11957.43 | -0.0009 | 0.0011 | 0.17 | 0.86270 |
| math - story | -0.0007 | 0.0005 | 11922.26 | -0.0017 | 0.0004 | -1.25 | 0.21112 |
| math - ToM | -0.0004 | 0.0005 | 11966.26 | -0.0014 | 0.0006 | -0.71 | 0.47527 |
| motor - punish | 0.0002 | 0.0005 | 11949.01 | -0.0008 | 0.0012 | 0.33 | 0.74387 |
| motor - random | -0.0004 | 0.0005 | 11993.60 | -0.0014 | 0.0006 | -0.81 | 0.42050 |
| motor - relational | 0.0001 | 0.0005 | 11983.22 | -0.0009 | 0.0011 | 0.26 | 0.79689 |
| motor - reward | 0.0002 | 0.0005 | 11949.01 | -0.0008 | 0.0012 | 0.34 | 0.73116 |
| motor - shapes | 0.0004 | 0.0005 | 11986.40 | -0.0006 | 0.0014 | 0.79 | 0.43115 |
| motor - story | -0.0004 | 0.0005 | 11992.67 | -0.0014 | 0.0007 | -0.68 | 0.49854 |
| motor - ToM | -0.0001 | 0.0005 | 11993.60 | -0.0011 | 0.0009 | -0.12 | 0.90440 |
| punish - random | -0.0006 | 0.0005 | 11976.28 | -0.0016 | 0.0004 | -1.14 | 0.25506 |
| punish - relational | 0.0000 | 0.0005 | 11982.11 | -0.0010 | 0.0010 | -0.06 | 0.94897 |
| punish - reward | 0.0000 | 0.0005 | 11922.26 | -0.0010 | 0.0010 | 0.02 | 0.98643 |
| punish - shapes | 0.0002 | 0.0005 | 11975.53 | -0.0008 | 0.0012 | 0.47 | 0.63925 |
| punish - story | -0.0005 | 0.0005 | 11997.02 | -0.0015 | 0.0005 | -1.00 | 0.31726 |
| punish - ToM | -0.0002 | 0.0005 | 11976.28 | -0.0012 | 0.0008 | -0.45 | 0.65521 |
| random - relational | 0.0005 | 0.0005 | 11948.16 | -0.0005 | 0.0015 | 1.06 | 0.29033 |
| random - reward | 0.0006 | 0.0005 | 11976.28 | -0.0004 | 0.0016 | 1.15 | 0.24812 |
| random - shapes | 0.0008 | 0.0005 | 11939.47 | -0.0002 | 0.0018 | 1.59 | 0.11147 |
| random - story | 0.0001 | 0.0005 | 11966.26 | -0.0010 | 0.0011 | 0.11 | 0.91076 |
| random - ToM | 0.0003 | 0.0005 | 11922.26 | -0.0006 | 0.0013 | 0.69 | 0.49294 |
| relational - reward | 0.0000 | 0.0005 | 11982.11 | -0.0010 | 0.0010 | 0.08 | 0.93570 |
| relational - shapes | 0.0003 | 0.0005 | 11934.36 | -0.0007 | 0.0013 | 0.53 | 0.59940 |
| relational - story | -0.0005 | 0.0005 | 11955.54 | -0.0015 | 0.0005 | -0.92 | 0.35546 |
| relational - ToM | -0.0002 | 0.0005 | 11948.16 | -0.0012 | 0.0008 | -0.38 | 0.70666 |
| reward - shapes | 0.0002 | 0.0005 | 11975.53 | -0.0008 | 0.0012 | 0.45 | 0.65129 |
| reward - story | -0.0005 | 0.0005 | 11997.02 | -0.0015 | 0.0005 | -1.02 | 0.30939 |
| reward - ToM | -0.0002 | 0.0005 | 11976.28 | -0.0012 | 0.0008 | -0.46 | 0.64314 |
| shapes - story | -0.0008 | 0.0005 | 11957.43 | -0.0018 | 0.0003 | -1.45 | 0.14833 |
| shapes - ToM | -0.0005 | 0.0005 | 11939.47 | -0.0015 | 0.0005 | -0.91 | 0.36464 |
| story - ToM | 0.0003 | 0.0005 | 11966.26 | -0.0007 | 0.0013 | 0.56 | 0.57680 |

81 **SI Table 8.4** Extraversion* task condition interactions along D2 (motor- visual)

| contrast | β | SE | df | $CL_{lower}$ | $CL_{upper}$ | t.ratio | p.value |
|---|---|---|---|---|---|---|---|
| 0back - 2back | 0.0000 | 0.0005 | 11922.26 | -0.0009 | 0.0010 | 0.07 | 0.94046 |
| 0back - faces | -0.0003 | 0.0005 | 11962.48 | -0.0013 | 0.0006 | -0.67 | 0.50264 |
| 0back - match | -0.0002 | 0.0005 | 11971.66 | -0.0012 | 0.0008 | -0.41 | 0.68107 |
| 0back - math | -0.0010 | 0.0005 | 11985.18 | -0.0019 | 0.0000 | -1.92 | 0.05503 |
| 0back - motor | -0.0014 | 0.0005 | 11943.04 | -0.0024 | -0.0005 | -2.91 | 0.00359 |
| 0back - punish | -0.0005 | 0.0005 | 11933.78 | -0.0015 | 0.0004 | -1.11 | 0.26782 |
| 0back - random | 0.0002 | 0.0005 | 11964.27 | -0.0007 | 0.0012 | 0.49 | 0.62455 |
| 0back - relational | -0.0004 | 0.0005 | 11971.66 | -0.0014 | 0.0006 | -0.80 | 0.42657 |
| 0back - reward | -0.0009 | 0.0005 | 11933.78 | -0.0019 | 0.0000 | -1.93 | 0.05401 |
| 0back - shapes | -0.0008 | 0.0005 | 11962.48 | -0.0018 | 0.0002 | -1.64 | 0.10030 |
| 0back - story | -0.0010 | 0.0005 | 11985.18 | -0.0020 | 0.0000 | -2.00 | 0.04560 |
| 0back - ToM | 0.0001 | 0.0005 | 11964.27 | -0.0009 | 0.0010 | 0.18 | 0.85626 |
| 2back - faces | -0.0004 | 0.0005 | 11962.48 | -0.0013 | 0.0006 | -0.74 | 0.45654 |
| 2back - match | -0.0002 | 0.0005 | 11971.66 | -0.0012 | 0.0007 | -0.48 | 0.62792 |
| 2back - math | -0.0010 | 0.0005 | 11985.18 | -0.0020 | 0.0000 | -1.99 | 0.04644 |
| 2back - motor | -0.0015 | 0.0005 | 11943.05 | -0.0024 | -0.0005 | -2.99 | 0.00282 |
| 2back - punish | -0.0006 | 0.0005 | 11933.79 | -0.0015 | 0.0004 | -1.18 | 0.23688 |
| 2back - random | 0.0002 | 0.0005 | 11964.27 | -0.0008 | 0.0012 | 0.42 | 0.67809 |
| 2back - relational | -0.0004 | 0.0005 | 11971.66 | -0.0014 | 0.0005 | -0.87 | 0.38499 |
| 2back - reward | -0.0010 | 0.0005 | 11933.79 | -0.0019 | 0.0000 | -2.00 | 0.04534 |
| 2back - shapes | -0.0008 | 0.0005 | 11962.48 | -0.0018 | 0.0001 | -1.72 | 0.08587 |
| 2back - story | -0.0010 | 0.0005 | 11985.18 | -0.0020 | -0.0001 | -2.07 | 0.03828 |
| 2back - ToM | 0.0001 | 0.0005 | 11964.27 | -0.0009 | 0.0010 | 0.11 | 0.91496 |
| faces - match | 0.0001 | 0.0005 | 11938.46 | -0.0008 | 0.0011 | 0.25 | 0.79976 |
| faces - math | -0.0006 | 0.0005 | 11958.76 | -0.0016 | 0.0004 | -1.26 | 0.20857 |
| faces - motor | -0.0011 | 0.0005 | 11969.31 | -0.0021 | -0.0001 | -2.23 | 0.02568 |
| faces - punish | -0.0002 | 0.0005 | 11959.56 | -0.0012 | 0.0007 | -0.43 | 0.66665 |
| faces - random | 0.0006 | 0.0005 | 11938.04 | -0.0004 | 0.0015 | 1.16 | 0.24720 |
| faces - relational | -0.0001 | 0.0005 | 11938.46 | -0.0010 | 0.0009 | -0.13 | 0.89705 |
| faces - reward | -0.0006 | 0.0005 | 11959.56 | -0.0016 | 0.0004 | -1.25 | 0.21315 |
| faces - shapes | -0.0005 | 0.0005 | 11922.26 | -0.0014 | 0.0005 | -0.97 | 0.33169 |
| faces - story | -0.0007 | 0.0005 | 11958.76 | -0.0017 | 0.0003 | -1.34 | 0.18097 |
| faces - ToM | 0.0004 | 0.0005 | 11938.04 | -0.0005 | 0.0014 | 0.85 | 0.39537 |
| match - math | -0.0008 | 0.0005 | 11947.83 | -0.0017 | 0.0002 | -1.50 | 0.13391 |
| match - motor | -0.0012 | 0.0005 | 11963.58 | -0.0022 | -0.0003 | -2.47 | 0.01354 |
| match - punish | -0.0003 | 0.0005 | 11968.54 | -0.0013 | 0.0006 | -0.68 | 0.49511 |
| match - random | 0.0004 | 0.0005 | 11946.69 | -0.0005 | 0.0014 | 0.89 | 0.37099 |
| match - relational | -0.0002 | 0.0005 | 11922.26 | -0.0012 | 0.0008 | -0.38 | 0.70337 |
| match - reward | -0.0007 | 0.0005 | 11968.54 | -0.0017 | 0.0002 | -1.49 | 0.13615 |

| | | | | | | | |
|---|---|---|---|---|---|---|---|
| match - shapes | -0.0006 | 0.0005 | 11938.46 | -0.0016 | 0.0004 | -1.22 | 0.22366 |
| match - story | -0.0008 | 0.0005 | 11947.83 | -0.0018 | 0.0002 | -1.58 | 0.11442 |
| match - ToM | 0.0003 | 0.0005 | 11946.69 | -0.0007 | 0.0013 | 0.59 | 0.55544 |
| math - motor | -0.0005 | 0.0005 | 11979.98 | -0.0015 | 0.0005 | -0.94 | 0.34917 |
| math - punish | 0.0004 | 0.0005 | 11982.45 | -0.0006 | 0.0014 | 0.84 | 0.40061 |
| math - random | 0.0012 | 0.0005 | 11955.72 | 0.0002 | 0.0022 | 2.40 | 0.01663 |
| math - relational | 0.0006 | 0.0005 | 11947.83 | -0.0004 | 0.0016 | 1.12 | 0.26151 |
| math - reward | 0.0000 | 0.0005 | 11982.45 | -0.0010 | 0.0010 | 0.04 | 0.96645 |
| math - shapes | 0.0002 | 0.0005 | 11958.76 | -0.0008 | 0.0011 | 0.31 | 0.75954 |
| math - story | 0.0000 | 0.0005 | 11922.26 | -0.0010 | 0.0010 | -0.08 | 0.93699 |
| math - ToM | 0.0010 | 0.0005 | 11955.72 | 0.0001 | 0.0020 | 2.09 | 0.03631 |
| motor - punish | 0.0009 | 0.0005 | 11938.83 | -0.0001 | 0.0019 | 1.82 | 0.06934 |
| motor - random | 0.0017 | 0.0005 | 11974.44 | 0.0007 | 0.0026 | 3.39 | 0.00071 |
| motor - relational | 0.0010 | 0.0005 | 11963.58 | 0.0001 | 0.0020 | 2.09 | 0.03685 |
| motor - reward | 0.0005 | 0.0005 | 11938.83 | -0.0005 | 0.0015 | 1.00 | 0.31579 |
| motor - shapes | 0.0006 | 0.0005 | 11969.31 | -0.0003 | 0.0016 | 1.26 | 0.20598 |
| motor - story | 0.0004 | 0.0005 | 11979.98 | -0.0006 | 0.0014 | 0.86 | 0.39190 |
| motor - ToM | 0.0015 | 0.0005 | 11974.44 | 0.0006 | 0.0025 | 3.08 | 0.00206 |
| punish - random | 0.0008 | 0.0005 | 11964.28 | -0.0002 | 0.0017 | 1.59 | 0.11101 |
| punish - relational | 0.0001 | 0.0005 | 11968.54 | -0.0008 | 0.0011 | 0.30 | 0.76599 |
| punish - reward | -0.0004 | 0.0005 | 11922.26 | -0.0014 | 0.0006 | -0.82 | 0.41211 |
| punish - shapes | -0.0003 | 0.0005 | 11959.56 | -0.0012 | 0.0007 | -0.54 | 0.58668 |
| punish - story | -0.0005 | 0.0005 | 11982.45 | -0.0014 | 0.0005 | -0.92 | 0.35698 |
| punish - ToM | 0.0006 | 0.0005 | 11964.28 | -0.0003 | 0.0016 | 1.29 | 0.19876 |
| random - relational | -0.0006 | 0.0005 | 11946.69 | -0.0016 | 0.0003 | -1.28 | 0.20120 |
| random - reward | -0.0012 | 0.0005 | 11964.28 | -0.0021 | -0.0002 | -2.41 | 0.01599 |
| random - shapes | -0.0010 | 0.0005 | 11938.04 | -0.0020 | -0.0001 | -2.13 | 0.03328 |
| random - story | -0.0012 | 0.0005 | 11955.72 | -0.0022 | -0.0003 | -2.48 | 0.01332 |
| random - ToM | -0.0002 | 0.0005 | 11922.26 | -0.0011 | 0.0008 | -0.31 | 0.75807 |
| relational - reward | -0.0005 | 0.0005 | 11968.54 | -0.0015 | 0.0004 | -1.11 | 0.26884 |
| relational - shapes | -0.0004 | 0.0005 | 11938.46 | -0.0014 | 0.0006 | -0.83 | 0.40437 |
| relational - story | -0.0006 | 0.0005 | 11947.83 | -0.0016 | 0.0004 | -1.20 | 0.22913 |
| relational - ToM | 0.0005 | 0.0005 | 11946.69 | -0.0005 | 0.0015 | 0.97 | 0.33048 |
| reward - shapes | 0.0001 | 0.0005 | 11959.56 | -0.0008 | 0.0011 | 0.27 | 0.78671 |
| reward - story | -0.0001 | 0.0005 | 11982.45 | -0.0010 | 0.0009 | -0.12 | 0.90236 |
| reward - ToM | 0.0010 | 0.0005 | 11964.28 | 0.0001 | 0.0020 | 2.10 | 0.03568 |
| shapes - story | -0.0002 | 0.0005 | 11958.76 | -0.0012 | 0.0008 | -0.39 | 0.69923 |
| shapes - ToM | 0.0009 | 0.0005 | 11938.04 | -0.0001 | 0.0019 | 1.82 | 0.06853 |
| story - ToM | 0.0011 | 0.0005 | 11955.72 | 0.0001 | 0.0021 | 2.17 | 0.02972 |

**SI Table 8.5** Agreeableness* task condition interactions along D2 (motor- visual)

| contrast | β | SE | df | CL $_{lower}$ | CL $_{upper}$ | t.ratio | p.value |
|---|---|---|---|---|---|---|---|
| 0back - 2back | -0.0001 | 0.0005 | 11922.26 | -0.0010 | 0.0009 | -0.10 | 0.91781 |
| 0back - faces | 0.0003 | 0.0005 | 11956.14 | -0.0006 | 0.0013 | 0.67 | 0.50312 |
| 0back - match | 0.0008 | 0.0005 | 11961.98 | -0.0002 | 0.0017 | 1.52 | 0.12859 |
| 0back - math | 0.0014 | 0.0005 | 11989.53 | 0.0004 | 0.0024 | 2.74 | 0.00611 |
| 0back - motor | 0.0022 | 0.0005 | 11937.59 | 0.0012 | 0.0032 | 4.41 | 0.00001 |
| 0back - punish | 0.0007 | 0.0005 | 11936.29 | -0.0003 | 0.0017 | 1.43 | 0.15328 |
| 0back - random | 0.0003 | 0.0005 | 11968.64 | -0.0007 | 0.0013 | 0.64 | 0.52122 |
| 0back - relational | 0.0004 | 0.0005 | 11961.98 | -0.0006 | 0.0013 | 0.73 | 0.46271 |
| 0back - reward | 0.0006 | 0.0005 | 11936.29 | -0.0004 | 0.0016 | 1.19 | 0.23517 |
| 0back - shapes | 0.0008 | 0.0005 | 11956.14 | -0.0001 | 0.0018 | 1.69 | 0.09190 |
| 0back - story | 0.0020 | 0.0005 | 11989.53 | 0.0010 | 0.0030 | 3.87 | 0.00011 |
| 0back - ToM | 0.0011 | 0.0005 | 11968.64 | 0.0002 | 0.0021 | 2.32 | 0.02028 |
| 2back - faces | 0.0004 | 0.0005 | 11956.14 | -0.0006 | 0.0014 | 0.77 | 0.44010 |
| 2back - match | 0.0008 | 0.0005 | 11961.98 | -0.0002 | 0.0018 | 1.62 | 0.10489 |
| 2back - math | 0.0015 | 0.0005 | 11989.53 | 0.0005 | 0.0025 | 2.84 | 0.00449 |
| 2back - motor | 0.0022 | 0.0005 | 11937.59 | 0.0013 | 0.0032 | 4.51 | 0.00001 |
| 2back - punish | 0.0008 | 0.0005 | 11936.30 | -0.0002 | 0.0017 | 1.53 | 0.12574 |
| 2back - random | 0.0004 | 0.0005 | 11968.65 | -0.0006 | 0.0013 | 0.74 | 0.45674 |
| 2back - relational | 0.0004 | 0.0005 | 11961.98 | -0.0006 | 0.0014 | 0.84 | 0.40299 |
| 2back - reward | 0.0006 | 0.0005 | 11936.30 | -0.0003 | 0.0016 | 1.29 | 0.19697 |
| 2back - shapes | 0.0009 | 0.0005 | 11956.14 | -0.0001 | 0.0019 | 1.79 | 0.07380 |
| 2back - story | 0.0020 | 0.0005 | 11989.53 | 0.0010 | 0.0030 | 3.97 | 0.00007 |
| 2back - ToM | 0.0012 | 0.0005 | 11968.65 | 0.0002 | 0.0022 | 2.42 | 0.01536 |
| faces - match | 0.0004 | 0.0005 | 11934.83 | -0.0006 | 0.0014 | 0.85 | 0.39552 |
| faces - math | 0.0011 | 0.0005 | 11967.26 | 0.0001 | 0.0021 | 2.08 | 0.03727 |
| faces - motor | 0.0019 | 0.0005 | 11963.11 | 0.0009 | 0.0028 | 3.72 | 0.00020 |
| faces - punish | 0.0004 | 0.0005 | 11953.27 | -0.0006 | 0.0013 | 0.75 | 0.45369 |
| faces - random | 0.0000 | 0.0005 | 11948.29 | -0.0010 | 0.0010 | -0.03 | 0.97595 |
| faces - relational | 0.0000 | 0.0005 | 11934.83 | -0.0009 | 0.0010 | 0.07 | 0.94586 |
| faces - reward | 0.0003 | 0.0005 | 11953.27 | -0.0007 | 0.0012 | 0.51 | 0.61009 |
| faces - shapes | 0.0005 | 0.0005 | 11922.26 | -0.0005 | 0.0015 | 1.01 | 0.31173 |
| faces - story | 0.0016 | 0.0005 | 11967.26 | 0.0006 | 0.0027 | 3.21 | 0.00134 |
| faces - ToM | 0.0008 | 0.0005 | 11948.29 | -0.0002 | 0.0018 | 1.64 | 0.10074 |
| match - math | 0.0006 | 0.0005 | 11962.78 | -0.0004 | 0.0017 | 1.25 | 0.21189 |
| match - motor | 0.0014 | 0.0005 | 11957.91 | 0.0004 | 0.0024 | 2.85 | 0.00437 |
| match - punish | -0.0001 | 0.0005 | 11959.86 | -0.0010 | 0.0009 | -0.11 | 0.91409 |
| match - random | -0.0004 | 0.0005 | 11957.22 | -0.0014 | 0.0005 | -0.88 | 0.37787 |
| match - relational | -0.0004 | 0.0005 | 11922.26 | -0.0014 | 0.0006 | -0.78 | 0.43624 |
| match - reward | -0.0002 | 0.0005 | 11959.86 | -0.0011 | 0.0008 | -0.35 | 0.72937 |

| | | | | | | | |
|---|---|---|---|---|---|---|---|
| match - shapes | 0.0001 | 0.0005 | 11934.83 | -0.0009 | 0.0011 | 0.16 | 0.87565 |
| match - story | 0.0012 | 0.0005 | 11962.78 | 0.0002 | 0.0022 | 2.37 | 0.01784 |
| match - ToM | 0.0004 | 0.0005 | 11957.22 | -0.0006 | 0.0014 | 0.78 | 0.43495 |
| math - motor | 0.0008 | 0.0005 | 11986.69 | -0.0002 | 0.0018 | 1.53 | 0.12630 |
| math - punish | -0.0007 | 0.0005 | 11984.16 | -0.0017 | 0.0003 | -1.36 | 0.17237 |
| math - random | -0.0011 | 0.0005 | 11974.75 | -0.0021 | -0.0001 | -2.12 | 0.03412 |
| math - relational | -0.0010 | 0.0005 | 11962.78 | -0.0020 | 0.0000 | -2.01 | 0.04467 |
| math - reward | -0.0008 | 0.0005 | 11984.16 | -0.0018 | 0.0002 | -1.60 | 0.11027 |
| math - shapes | -0.0006 | 0.0005 | 11967.26 | -0.0016 | 0.0004 | -1.10 | 0.27083 |
| math - story | 0.0006 | 0.0005 | 11922.26 | -0.0005 | 0.0016 | 1.10 | 0.27148 |
| math - ToM | -0.0003 | 0.0005 | 11974.75 | -0.0013 | 0.0007 | -0.50 | 0.62022 |
| motor - punish | -0.0015 | 0.0005 | 11939.70 | -0.0025 | -0.0005 | -2.99 | 0.00279 |
| motor - random | -0.0019 | 0.0005 | 11979.36 | -0.0028 | -0.0009 | -3.76 | 0.00017 |
| motor - relational | -0.0018 | 0.0005 | 11957.91 | -0.0028 | -0.0008 | -3.63 | 0.00028 |
| motor - reward | -0.0016 | 0.0005 | 11939.70 | -0.0026 | -0.0006 | -3.23 | 0.00124 |
| motor - shapes | -0.0013 | 0.0005 | 11963.11 | -0.0023 | -0.0004 | -2.71 | 0.00681 |
| motor - story | -0.0002 | 0.0005 | 11986.69 | -0.0012 | 0.0008 | -0.40 | 0.68618 |
| motor - ToM | -0.0010 | 0.0005 | 11979.36 | -0.0020 | -0.0001 | -2.09 | 0.03681 |
| punish - random | -0.0004 | 0.0005 | 11967.54 | -0.0014 | 0.0006 | -0.78 | 0.43428 |
| punish - relational | -0.0003 | 0.0005 | 11959.86 | -0.0013 | 0.0006 | -0.68 | 0.49830 |
| punish - reward | -0.0001 | 0.0005 | 11922.26 | -0.0011 | 0.0008 | -0.24 | 0.80957 |
| punish - shapes | 0.0001 | 0.0005 | 11953.27 | -0.0008 | 0.0011 | 0.27 | 0.78992 |
| punish - story | 0.0013 | 0.0005 | 11984.16 | 0.0003 | 0.0023 | 2.50 | 0.01260 |
| punish - ToM | 0.0004 | 0.0005 | 11967.54 | -0.0005 | 0.0014 | 0.90 | 0.36952 |
| random - relational | 0.0000 | 0.0005 | 11957.22 | -0.0009 | 0.0010 | 0.10 | 0.92189 |
| random - reward | 0.0003 | 0.0005 | 11967.54 | -0.0007 | 0.0012 | 0.54 | 0.58795 |
| random - shapes | 0.0005 | 0.0005 | 11948.29 | -0.0005 | 0.0015 | 1.04 | 0.29636 |
| random - story | 0.0017 | 0.0005 | 11974.75 | 0.0007 | 0.0027 | 3.25 | 0.00117 |
| random - ToM | 0.0008 | 0.0005 | 11922.26 | -0.0001 | 0.0018 | 1.68 | 0.09316 |
| relational - reward | 0.0002 | 0.0005 | 11959.86 | -0.0008 | 0.0012 | 0.44 | 0.66060 |
| relational - shapes | 0.0005 | 0.0005 | 11934.83 | -0.0005 | 0.0015 | 0.94 | 0.34812 |
| relational - story | 0.0016 | 0.0005 | 11962.78 | 0.0006 | 0.0026 | 3.13 | 0.00176 |
| relational - ToM | 0.0008 | 0.0005 | 11957.22 | -0.0002 | 0.0018 | 1.56 | 0.11771 |
| reward - shapes | 0.0003 | 0.0005 | 11953.27 | -0.0007 | 0.0012 | 0.51 | 0.61303 |
| reward - story | 0.0014 | 0.0005 | 11984.16 | 0.0004 | 0.0024 | 2.73 | 0.00639 |
| reward - ToM | 0.0006 | 0.0005 | 11967.54 | -0.0004 | 0.0015 | 1.14 | 0.25535 |
| shapes - story | 0.0011 | 0.0005 | 11967.26 | 0.0001 | 0.0021 | 2.23 | 0.02598 |
| shapes - ToM | 0.0003 | 0.0005 | 11948.29 | -0.0007 | 0.0013 | 0.63 | 0.53052 |
| story - ToM | -0.0008 | 0.0005 | 11974.75 | -0.0018 | 0.0002 | -1.62 | 0.10424 |

**SI Table 9.1** Neuroticism* task condition interactions along D3 (DMN- FPN)

| contrast | β | SE | df | $CL_{lower}$ | $CL_{upper}$ | t.ratio | p.value |
|---|---|---|---|---|---|---|---|
| 0back - 2back | 0.0012 | 0.0005 | 11937.12 | 0.0003 | 0.0021 | 2.61 | 0.00903 |
| 0back - faces | 0.0007 | 0.0005 | 11991.40 | -0.0003 | 0.0016 | 1.43 | 0.15386 |
| 0back - match | 0.0005 | 0.0005 | 11995.39 | -0.0004 | 0.0015 | 1.09 | 0.27660 |
| 0back - math | 0.0011 | 0.0005 | 12006.01 | 0.0002 | 0.0021 | 2.32 | 0.02061 |
| 0back - motor | 0.0008 | 0.0005 | 11957.22 | -0.0001 | 0.0017 | 1.70 | 0.08895 |
| 0back - punish | 0.0001 | 0.0005 | 11959.99 | -0.0009 | 0.0010 | 0.13 | 0.89661 |
| 0back - random | 0.0015 | 0.0005 | 11997.00 | 0.0006 | 0.0024 | 3.16 | 0.00156 |
| 0back - relational | 0.0008 | 0.0005 | 11995.39 | -0.0001 | 0.0018 | 1.77 | 0.07730 |
| 0back - reward | 0.0001 | 0.0005 | 11959.99 | -0.0008 | 0.0010 | 0.15 | 0.87796 |
| 0back - shapes | 0.0007 | 0.0005 | 11991.40 | -0.0002 | 0.0016 | 1.51 | 0.12998 |
| 0back - story | 0.0001 | 0.0005 | 12006.01 | -0.0009 | 0.0010 | 0.18 | 0.85639 |
| 0back - ToM | 0.0005 | 0.0005 | 11997.00 | -0.0004 | 0.0015 | 1.11 | 0.26684 |
| 2back - faces | -0.0005 | 0.0005 | 11991.41 | -0.0015 | 0.0004 | -1.16 | 0.24695 |
| 2back - match | -0.0007 | 0.0005 | 11995.39 | -0.0016 | 0.0002 | -1.48 | 0.13812 |
| 2back - math | -0.0001 | 0.0005 | 12006.01 | -0.0011 | 0.0008 | -0.22 | 0.82942 |
| 2back - motor | -0.0004 | 0.0005 | 11957.23 | -0.0013 | 0.0005 | -0.90 | 0.36961 |
| 2back - punish | -0.0012 | 0.0005 | 11959.99 | -0.0021 | -0.0002 | -2.48 | 0.01317 |
| 2back - random | 0.0003 | 0.0005 | 11997.01 | -0.0007 | 0.0012 | 0.57 | 0.56663 |
| 2back - relational | -0.0004 | 0.0005 | 11995.39 | -0.0013 | 0.0006 | -0.80 | 0.42127 |
| 2back - reward | -0.0012 | 0.0005 | 11959.99 | -0.0021 | -0.0002 | -2.46 | 0.01406 |
| 2back - shapes | -0.0005 | 0.0005 | 11991.41 | -0.0014 | 0.0004 | -1.07 | 0.28479 |
| 2back - story | -0.0011 | 0.0005 | 12006.01 | -0.0021 | -0.0002 | -2.35 | 0.01880 |
| 2back - ToM | -0.0007 | 0.0005 | 11997.01 | -0.0016 | 0.0002 | -1.48 | 0.13867 |
| faces - match | -0.0002 | 0.0005 | 11951.24 | -0.0011 | 0.0008 | -0.33 | 0.74243 |
| faces - math | 0.0004 | 0.0005 | 11975.56 | -0.0005 | 0.0014 | 0.91 | 0.36177 |
| faces - motor | 0.0001 | 0.0005 | 12000.06 | -0.0008 | 0.0011 | 0.26 | 0.79149 |
| faces - punish | -0.0006 | 0.0005 | 11989.64 | -0.0015 | 0.0003 | -1.30 | 0.19440 |
| faces - random | 0.0008 | 0.0005 | 11954.03 | -0.0001 | 0.0018 | 1.72 | 0.08510 |
| faces - relational | 0.0002 | 0.0005 | 11951.24 | -0.0008 | 0.0011 | 0.35 | 0.72979 |
| faces - reward | -0.0006 | 0.0005 | 11989.64 | -0.0015 | 0.0003 | -1.27 | 0.20256 |
| faces - shapes | 0.0000 | 0.0005 | 11937.12 | -0.0009 | 0.0010 | 0.09 | 0.93019 |
| faces - story | -0.0006 | 0.0005 | 11975.56 | -0.0015 | 0.0004 | -1.21 | 0.22740 |
| faces - ToM | -0.0002 | 0.0005 | 11954.03 | -0.0011 | 0.0008 | -0.32 | 0.75088 |
| match - math | 0.0006 | 0.0005 | 11968.70 | -0.0004 | 0.0016 | 1.23 | 0.21870 |
| match - motor | 0.0003 | 0.0005 | 11996.99 | -0.0007 | 0.0012 | 0.59 | 0.55340 |
| match - punish | -0.0005 | 0.0005 | 11999.00 | -0.0014 | 0.0005 | -0.96 | 0.33707 |
| match - random | 0.0010 | 0.0005 | 11962.35 | 0.0000 | 0.0019 | 2.04 | 0.04112 |
| match - relational | 0.0003 | 0.0005 | 11937.12 | -0.0006 | 0.0013 | 0.67 | 0.50205 |
| match - reward | -0.0004 | 0.0005 | 11999.00 | -0.0014 | 0.0005 | -0.94 | 0.34890 |

| | | | | | | | |
|---|---|---|---|---|---|---|---|
| match - shapes | 0.0002 | 0.0005 | 11951.24 | -0.0007 | 0.0011 | 0.42 | 0.67757 |
| match - story | -0.0004 | 0.0005 | 11968.70 | -0.0014 | 0.0005 | -0.88 | 0.37895 |
| match - ToM | 0.0000 | 0.0005 | 11962.35 | -0.0009 | 0.0009 | 0.01 | 0.98913 |
| math - motor | -0.0003 | 0.0005 | 12011.20 | -0.0013 | 0.0006 | -0.66 | 0.51203 |
| math - punish | -0.0011 | 0.0005 | 12013.91 | -0.0020 | -0.0001 | -2.19 | 0.02863 |
| math - random | 0.0004 | 0.0005 | 11980.23 | -0.0006 | 0.0013 | 0.77 | 0.44034 |
| math - relational | -0.0003 | 0.0005 | 11968.70 | -0.0012 | 0.0007 | -0.57 | 0.56894 |
| math - reward | -0.0011 | 0.0005 | 12013.91 | -0.0020 | -0.0001 | -2.17 | 0.03033 |
| math - shapes | -0.0004 | 0.0005 | 11975.56 | -0.0014 | 0.0006 | -0.83 | 0.40866 |
| math - story | -0.0010 | 0.0005 | 11937.12 | -0.0020 | -0.0001 | -2.08 | 0.03723 |
| math - ToM | -0.0006 | 0.0005 | 11980.23 | -0.0016 | 0.0004 | -1.23 | 0.22047 |
| motor - punish | -0.0007 | 0.0005 | 11960.17 | -0.0017 | 0.0002 | -1.57 | 0.11605 |
| motor - random | 0.0007 | 0.0005 | 12007.73 | -0.0002 | 0.0016 | 1.46 | 0.14404 |
| motor - relational | 0.0000 | 0.0005 | 11996.99 | -0.0009 | 0.0010 | 0.08 | 0.93370 |
| motor - reward | -0.0007 | 0.0005 | 11960.17 | -0.0017 | 0.0002 | -1.55 | 0.12161 |
| motor - shapes | -0.0001 | 0.0005 | 12000.06 | -0.0010 | 0.0009 | -0.18 | 0.85982 |
| motor - story | -0.0007 | 0.0005 | 12011.20 | -0.0017 | 0.0002 | -1.47 | 0.14166 |
| motor - ToM | -0.0003 | 0.0005 | 12007.73 | -0.0012 | 0.0007 | -0.58 | 0.55967 |
| punish - random | 0.0014 | 0.0005 | 11995.58 | 0.0005 | 0.0024 | 3.04 | 0.00240 |
| punish - relational | 0.0008 | 0.0005 | 11999.00 | -0.0002 | 0.0017 | 1.64 | 0.10130 |
| punish - reward | 0.0000 | 0.0005 | 11937.12 | -0.0009 | 0.0009 | 0.02 | 0.98115 |
| punish - shapes | 0.0007 | 0.0005 | 11989.64 | -0.0003 | 0.0016 | 1.39 | 0.16579 |
| punish - story | 0.0000 | 0.0005 | 12013.91 | -0.0009 | 0.0010 | 0.05 | 0.95619 |
| punish - ToM | 0.0005 | 0.0005 | 11995.58 | -0.0005 | 0.0014 | 0.98 | 0.32635 |
| random - relational | -0.0007 | 0.0005 | 11962.35 | -0.0016 | 0.0003 | -1.37 | 0.17172 |
| random - reward | -0.0014 | 0.0005 | 11995.58 | -0.0024 | -0.0005 | -3.01 | 0.00260 |
| random - shapes | -0.0008 | 0.0005 | 11954.03 | -0.0017 | 0.0002 | -1.63 | 0.10225 |
| random - story | -0.0014 | 0.0005 | 11980.23 | -0.0024 | -0.0005 | -2.90 | 0.00378 |
| random - ToM | -0.0010 | 0.0005 | 11937.12 | -0.0019 | 0.0000 | -2.05 | 0.04073 |
| relational - reward | -0.0008 | 0.0005 | 11999.00 | -0.0017 | 0.0002 | -1.62 | 0.10624 |
| relational - shapes | -0.0001 | 0.0005 | 11951.24 | -0.0011 | 0.0008 | -0.26 | 0.79620 |
| relational - story | -0.0008 | 0.0005 | 11968.70 | -0.0017 | 0.0002 | -1.54 | 0.12351 |
| relational - ToM | -0.0003 | 0.0005 | 11962.35 | -0.0013 | 0.0006 | -0.66 | 0.50792 |
| reward - shapes | 0.0006 | 0.0005 | 11989.64 | -0.0003 | 0.0016 | 1.36 | 0.17305 |
| reward - story | 0.0000 | 0.0005 | 12013.91 | -0.0009 | 0.0010 | 0.03 | 0.97444 |
| reward - ToM | 0.0005 | 0.0005 | 11995.58 | -0.0005 | 0.0014 | 0.96 | 0.33804 |
| shapes - story | -0.0006 | 0.0005 | 11975.56 | -0.0016 | 0.0003 | -1.29 | 0.19607 |
| shapes - ToM | -0.0002 | 0.0005 | 11954.03 | -0.0011 | 0.0007 | -0.41 | 0.68527 |
| story - ToM | 0.0004 | 0.0005 | 11980.23 | -0.0005 | 0.0014 | 0.90 | 0.36841 |

**SI Table 9.2** Openness* task condition interactions along D3 (DMN- FPN)

| contrast | β | SE | df | CL lower | CL upper | t.ratio | p.value |
|---|---|---|---|---|---|---|---|
| 0back - 2back | -0.0015 | 0.0005 | 11937.12 | -0.0025 | -0.0005 | -2.99 | 0.00284 |
| 0back - faces | -0.0005 | 0.0005 | 11987.74 | -0.0015 | 0.0005 | -1.06 | 0.29013 |
| 0back - match | -0.0002 | 0.0005 | 11993.24 | -0.0012 | 0.0008 | -0.38 | 0.70488 |
| 0back - math | -0.0012 | 0.0005 | 12010.45 | -0.0022 | -0.0002 | -2.32 | 0.02022 |
| 0back - motor | -0.0015 | 0.0005 | 11960.29 | -0.0025 | -0.0006 | -3.06 | 0.00218 |
| 0back - punish | -0.0014 | 0.0005 | 11957.25 | -0.0023 | -0.0004 | -2.73 | 0.00627 |
| 0back - random | -0.0016 | 0.0005 | 11988.51 | -0.0026 | -0.0006 | -3.17 | 0.00153 |
| 0back - relational | -0.0009 | 0.0005 | 11993.24 | -0.0019 | 0.0001 | -1.75 | 0.08007 |
| 0back - reward | -0.0018 | 0.0005 | 11957.25 | -0.0027 | -0.0008 | -3.58 | 0.00035 |
| 0back - shapes | -0.0013 | 0.0005 | 11987.74 | -0.0023 | -0.0003 | -2.59 | 0.00963 |
| 0back - story | 0.0006 | 0.0005 | 12010.45 | -0.0004 | 0.0016 | 1.10 | 0.27095 |
| 0back - ToM | 0.0000 | 0.0005 | 11988.51 | -0.0010 | 0.0010 | 0.03 | 0.97679 |
| 2back - faces | 0.0010 | 0.0005 | 11987.74 | 0.0000 | 0.0019 | 1.90 | 0.05806 |
| 2back - match | 0.0013 | 0.0005 | 11993.24 | 0.0003 | 0.0023 | 2.56 | 0.01054 |
| 2back - math | 0.0003 | 0.0005 | 12010.45 | -0.0007 | 0.0013 | 0.56 | 0.57465 |
| 2back - motor | 0.0000 | 0.0005 | 11960.29 | -0.0010 | 0.0009 | -0.10 | 0.92103 |
| 2back - punish | 0.0001 | 0.0005 | 11957.25 | -0.0008 | 0.0011 | 0.26 | 0.79641 |
| 2back - random | -0.0001 | 0.0005 | 11988.51 | -0.0011 | 0.0009 | -0.21 | 0.83547 |
| 2back - relational | 0.0006 | 0.0005 | 11993.24 | -0.0004 | 0.0016 | 1.19 | 0.23551 |
| 2back - reward | -0.0003 | 0.0005 | 11957.25 | -0.0013 | 0.0007 | -0.59 | 0.55798 |
| 2back - shapes | 0.0002 | 0.0005 | 11987.74 | -0.0008 | 0.0012 | 0.36 | 0.71588 |
| 2back - story | 0.0021 | 0.0005 | 12010.45 | 0.0010 | 0.0031 | 3.98 | 0.00007 |
| 2back - ToM | 0.0015 | 0.0005 | 11988.51 | 0.0005 | 0.0025 | 2.99 | 0.00278 |
| faces - match | 0.0003 | 0.0005 | 11959.50 | -0.0007 | 0.0013 | 0.67 | 0.50396 |
| faces - math | -0.0007 | 0.0005 | 11984.64 | -0.0017 | 0.0004 | -1.28 | 0.20047 |
| faces - motor | -0.0010 | 0.0005 | 11998.97 | -0.0020 | 0.0000 | -1.98 | 0.04745 |
| faces - punish | -0.0008 | 0.0005 | 11988.13 | -0.0018 | 0.0002 | -1.65 | 0.09985 |
| faces - random | -0.0011 | 0.0005 | 11949.12 | -0.0020 | -0.0001 | -2.09 | 0.03631 |
| faces - relational | -0.0004 | 0.0005 | 11959.50 | -0.0014 | 0.0006 | -0.69 | 0.48807 |
| faces - reward | -0.0012 | 0.0005 | 11988.13 | -0.0022 | -0.0003 | -2.48 | 0.01311 |
| faces - shapes | -0.0008 | 0.0005 | 11937.12 | -0.0018 | 0.0002 | -1.52 | 0.12822 |
| faces - story | 0.0011 | 0.0005 | 11984.64 | 0.0001 | 0.0021 | 2.12 | 0.03416 |
| faces - ToM | 0.0005 | 0.0005 | 11949.12 | -0.0004 | 0.0015 | 1.08 | 0.27913 |
| match - math | -0.0010 | 0.0005 | 11972.06 | -0.0020 | 0.0000 | -1.93 | 0.05392 |
| match - motor | -0.0013 | 0.0005 | 11995.66 | -0.0023 | -0.0003 | -2.64 | 0.00826 |
| match - punish | -0.0012 | 0.0005 | 12001.58 | -0.0022 | -0.0002 | -2.31 | 0.02086 |
| match - random | -0.0014 | 0.0005 | 11964.58 | -0.0024 | -0.0004 | -2.75 | 0.00594 |
| match - relational | -0.0007 | 0.0005 | 11937.12 | -0.0017 | 0.0003 | -1.36 | 0.17516 |
| match - reward | -0.0016 | 0.0005 | 12001.58 | -0.0026 | -0.0006 | -3.14 | 0.00169 |

| | | | | | | | |
|---|---|---|---|---|---|---|---|
| match - shapes | -0.0011 | 0.0005 | 11959.50 | -0.0021 | -0.0001 | -2.18 | 0.02927 |
| match - story | 0.0008 | 0.0005 | 11972.06 | -0.0003 | 0.0018 | 1.46 | 0.14556 |
| match - ToM | 0.0002 | 0.0005 | 11964.58 | -0.0008 | 0.0012 | 0.41 | 0.68494 |
| math - motor | -0.0003 | 0.0005 | 12009.06 | -0.0014 | 0.0007 | -0.65 | 0.51294 |
| math - punish | -0.0002 | 0.0005 | 12017.39 | -0.0012 | 0.0008 | -0.31 | 0.75402 |
| math - random | -0.0004 | 0.0005 | 11982.40 | -0.0014 | 0.0006 | -0.76 | 0.44720 |
| math - relational | 0.0003 | 0.0005 | 11972.06 | -0.0007 | 0.0013 | 0.60 | 0.55078 |
| math - reward | -0.0006 | 0.0005 | 12017.39 | -0.0016 | 0.0004 | -1.13 | 0.25899 |
| math - shapes | -0.0001 | 0.0005 | 11984.64 | -0.0011 | 0.0009 | -0.20 | 0.83811 |
| math - story | 0.0018 | 0.0005 | 11937.12 | 0.0007 | 0.0028 | 3.34 | 0.00086 |
| math - ToM | 0.0012 | 0.0005 | 11982.40 | 0.0002 | 0.0022 | 2.34 | 0.01926 |
| motor - punish | 0.0002 | 0.0005 | 11961.01 | -0.0008 | 0.0012 | 0.36 | 0.72190 |
| motor - random | -0.0001 | 0.0005 | 11997.48 | -0.0010 | 0.0009 | -0.11 | 0.91397 |
| motor - relational | 0.0006 | 0.0005 | 11995.66 | -0.0003 | 0.0016 | 1.28 | 0.20143 |
| motor - reward | -0.0002 | 0.0005 | 11961.01 | -0.0012 | 0.0007 | -0.48 | 0.62908 |
| motor - shapes | 0.0002 | 0.0005 | 11998.97 | -0.0008 | 0.0012 | 0.46 | 0.64553 |
| motor - story | 0.0021 | 0.0005 | 12009.06 | 0.0011 | 0.0031 | 4.06 | 0.00005 |
| motor - ToM | 0.0015 | 0.0005 | 11997.48 | 0.0006 | 0.0025 | 3.07 | 0.00213 |
| punish - random | -0.0002 | 0.0005 | 11982.77 | -0.0012 | 0.0007 | -0.46 | 0.64221 |
| punish - relational | 0.0005 | 0.0005 | 12001.58 | -0.0005 | 0.0015 | 0.94 | 0.34952 |
| punish - reward | -0.0004 | 0.0005 | 11937.12 | -0.0014 | 0.0006 | -0.85 | 0.39702 |
| punish - shapes | 0.0001 | 0.0005 | 11988.13 | -0.0009 | 0.0010 | 0.11 | 0.91270 |
| punish - story | 0.0019 | 0.0005 | 12017.39 | 0.0009 | 0.0029 | 3.75 | 0.00018 |
| punish - ToM | 0.0014 | 0.0005 | 11982.77 | 0.0004 | 0.0024 | 2.74 | 0.00607 |
| random - relational | 0.0007 | 0.0005 | 11964.58 | -0.0003 | 0.0017 | 1.39 | 0.16580 |
| random - reward | -0.0002 | 0.0005 | 11982.77 | -0.0012 | 0.0008 | -0.37 | 0.70872 |
| random - shapes | 0.0003 | 0.0005 | 11949.12 | -0.0007 | 0.0013 | 0.57 | 0.56961 |
| random - story | 0.0022 | 0.0005 | 11982.40 | 0.0011 | 0.0032 | 4.17 | 0.00003 |
| random - ToM | 0.0016 | 0.0005 | 11937.12 | 0.0006 | 0.0026 | 3.19 | 0.00144 |
| relational - reward | -0.0009 | 0.0005 | 12001.58 | -0.0019 | 0.0001 | -1.77 | 0.07741 |
| relational - shapes | -0.0004 | 0.0005 | 11959.50 | -0.0014 | 0.0006 | -0.82 | 0.41315 |
| relational - story | 0.0015 | 0.0005 | 11972.06 | 0.0004 | 0.0025 | 2.79 | 0.00534 |
| relational - ToM | 0.0009 | 0.0005 | 11964.58 | -0.0001 | 0.0019 | 1.77 | 0.07655 |
| reward - shapes | 0.0005 | 0.0005 | 11988.13 | -0.0005 | 0.0015 | 0.95 | 0.34461 |
| reward - story | 0.0023 | 0.0005 | 12017.39 | 0.0013 | 0.0034 | 4.56 | 0.00001 |
| reward - ToM | 0.0018 | 0.0005 | 11982.77 | 0.0008 | 0.0028 | 3.58 | 0.00034 |
| shapes - story | 0.0019 | 0.0005 | 11984.64 | 0.0009 | 0.0029 | 3.60 | 0.00032 |
| shapes - ToM | 0.0013 | 0.0005 | 11949.12 | 0.0003 | 0.0023 | 2.61 | 0.00914 |
| story - ToM | -0.0006 | 0.0005 | 11982.40 | -0.0016 | 0.0005 | -1.07 | 0.28556 |

**SI Table 9.3** Conscientiousness* task condition interactions along D3 (DMN- FPN)

| contrast | β | SE | df | $CL_{lower}$ | $CL_{upper}$ | t.ratio | p.value |
|---|---|---|---|---|---|---|---|
| 0back - 2back | 0.0016 | 0.0006 | 11937.12 | 0.0005 | 0.0027 | 2.81 | 0.00501 |
| 0back - faces | 0.0020 | 0.0006 | 12000.84 | 0.0009 | 0.0032 | 3.53 | 0.00042 |
| 0back - match | 0.0011 | 0.0006 | 12005.49 | -0.0001 | 0.0022 | 1.87 | 0.06105 |
| 0back - math | 0.0013 | 0.0006 | 12024.37 | 0.0002 | 0.0025 | 2.26 | 0.02388 |
| 0back - motor | 0.0025 | 0.0006 | 11953.33 | 0.0013 | 0.0036 | 4.30 | 0.00002 |
| 0back - punish | 0.0006 | 0.0006 | 11960.85 | -0.0005 | 0.0017 | 1.10 | 0.27291 |
| 0back - random | 0.0015 | 0.0006 | 12008.75 | 0.0004 | 0.0026 | 2.58 | 0.00993 |
| 0back - relational | 0.0014 | 0.0006 | 12005.49 | 0.0003 | 0.0026 | 2.45 | 0.01419 |
| 0back - reward | 0.0002 | 0.0006 | 11960.85 | -0.0009 | 0.0014 | 0.41 | 0.67858 |
| 0back - shapes | 0.0022 | 0.0006 | 12000.84 | 0.0011 | 0.0033 | 3.82 | 0.00013 |
| 0back - story | 0.0005 | 0.0006 | 12024.37 | -0.0007 | 0.0016 | 0.77 | 0.43853 |
| 0back - ToM | 0.0007 | 0.0006 | 12008.75 | -0.0004 | 0.0018 | 1.22 | 0.22187 |
| 2back - faces | 0.0004 | 0.0006 | 12000.84 | -0.0007 | 0.0016 | 0.75 | 0.45403 |
| 2back - match | -0.0005 | 0.0006 | 12005.49 | -0.0016 | 0.0006 | -0.89 | 0.37443 |
| 2back - math | -0.0003 | 0.0006 | 12024.37 | -0.0014 | 0.0009 | -0.46 | 0.64722 |
| 2back - motor | 0.0009 | 0.0006 | 11953.33 | -0.0003 | 0.0020 | 1.51 | 0.13072 |
| 2back - punish | -0.0010 | 0.0006 | 11960.85 | -0.0021 | 0.0001 | -1.72 | 0.08605 |
| 2back - random | -0.0001 | 0.0006 | 12008.75 | -0.0012 | 0.0010 | -0.20 | 0.84318 |
| 2back - relational | -0.0002 | 0.0006 | 12005.49 | -0.0013 | 0.0010 | -0.31 | 0.75748 |
| 2back - reward | -0.0014 | 0.0006 | 11960.85 | -0.0025 | -0.0002 | -2.40 | 0.01647 |
| 2back - shapes | 0.0006 | 0.0006 | 12000.84 | -0.0005 | 0.0017 | 1.04 | 0.29624 |
| 2back - story | -0.0011 | 0.0006 | 12024.37 | -0.0023 | 0.0000 | -1.94 | 0.05212 |
| 2back - ToM | -0.0009 | 0.0006 | 12008.75 | -0.0020 | 0.0002 | -1.55 | 0.11998 |
| faces - match | -0.0009 | 0.0006 | 11950.73 | -0.0021 | 0.0002 | -1.62 | 0.10464 |
| faces - math | -0.0007 | 0.0006 | 11977.40 | -0.0019 | 0.0005 | -1.18 | 0.23687 |
| faces - motor | 0.0004 | 0.0006 | 12007.61 | -0.0007 | 0.0016 | 0.75 | 0.45231 |
| faces - punish | -0.0014 | 0.0006 | 11995.27 | -0.0025 | -0.0003 | -2.45 | 0.01423 |
| faces - random | -0.0005 | 0.0006 | 11956.08 | -0.0017 | 0.0006 | -0.94 | 0.34699 |
| faces - relational | -0.0006 | 0.0006 | 11950.73 | -0.0018 | 0.0005 | -1.05 | 0.29516 |
| faces - reward | -0.0018 | 0.0006 | 11995.27 | -0.0029 | -0.0007 | -3.13 | 0.00177 |
| faces - shapes | 0.0002 | 0.0006 | 11937.12 | -0.0010 | 0.0013 | 0.29 | 0.76865 |
| faces - story | -0.0016 | 0.0006 | 11977.40 | -0.0027 | -0.0004 | -2.66 | 0.00786 |
| faces - ToM | -0.0013 | 0.0006 | 11956.08 | -0.0025 | -0.0002 | -2.29 | 0.02208 |
| match - math | 0.0002 | 0.0006 | 11975.22 | -0.0009 | 0.0014 | 0.41 | 0.68061 |
| match - motor | 0.0014 | 0.0006 | 12004.16 | 0.0002 | 0.0025 | 2.37 | 0.01771 |
| match - punish | -0.0005 | 0.0006 | 12002.80 | -0.0016 | 0.0007 | -0.80 | 0.42402 |
| match - random | 0.0004 | 0.0006 | 11966.06 | -0.0007 | 0.0015 | 0.69 | 0.49215 |
| match - relational | 0.0003 | 0.0006 | 11937.12 | -0.0008 | 0.0015 | 0.57 | 0.56650 |
| match - reward | -0.0008 | 0.0006 | 12002.80 | -0.0020 | 0.0003 | -1.47 | 0.14130 |

| | | | | | | | |
|---|---|---|---|---|---|---|---|
| match - shapes | 0.0011 | 0.0006 | 11950.73 | 0.0000 | 0.0023 | 1.92 | 0.05548 |
| match - story | -0.0006 | 0.0006 | 11975.22 | -0.0018 | 0.0005 | -1.06 | 0.29073 |
| match - ToM | -0.0004 | 0.0006 | 11966.06 | -0.0015 | 0.0008 | -0.65 | 0.51314 |
| math - motor | 0.0011 | 0.0006 | 12014.87 | 0.0000 | 0.0023 | 1.92 | 0.05478 |
| math - punish | -0.0007 | 0.0006 | 12020.13 | -0.0019 | 0.0004 | -1.20 | 0.22856 |
| math - random | 0.0002 | 0.0006 | 11987.21 | -0.0010 | 0.0013 | 0.26 | 0.79305 |
| math - relational | 0.0001 | 0.0006 | 11975.22 | -0.0011 | 0.0013 | 0.15 | 0.87909 |
| math - reward | -0.0011 | 0.0006 | 12020.13 | -0.0022 | 0.0001 | -1.86 | 0.06222 |
| math - shapes | 0.0009 | 0.0006 | 11977.40 | -0.0003 | 0.0020 | 1.47 | 0.14143 |
| math - story | -0.0009 | 0.0006 | 11937.12 | -0.0021 | 0.0003 | -1.45 | 0.14710 |
| math - ToM | -0.0006 | 0.0006 | 11987.21 | -0.0018 | 0.0005 | -1.06 | 0.29048 |
| motor - punish | -0.0018 | 0.0006 | 11966.79 | -0.0030 | -0.0007 | -3.22 | 0.00127 |
| motor - random | -0.0010 | 0.0006 | 12015.48 | -0.0021 | 0.0002 | -1.69 | 0.09057 |
| motor - relational | -0.0010 | 0.0006 | 12004.16 | -0.0022 | 0.0001 | -1.80 | 0.07260 |
| motor - reward | -0.0022 | 0.0006 | 11966.79 | -0.0034 | -0.0011 | -3.90 | 0.00010 |
| motor - shapes | -0.0003 | 0.0006 | 12007.61 | -0.0014 | 0.0009 | -0.46 | 0.64743 |
| motor - story | -0.0020 | 0.0006 | 12014.87 | -0.0032 | -0.0009 | -3.40 | 0.00068 |
| motor - ToM | -0.0018 | 0.0006 | 12015.48 | -0.0029 | -0.0006 | -3.04 | 0.00235 |
| punish - random | 0.0009 | 0.0006 | 11996.42 | -0.0003 | 0.0020 | 1.50 | 0.13318 |
| punish - relational | 0.0008 | 0.0006 | 12002.80 | -0.0003 | 0.0019 | 1.38 | 0.16739 |
| punish - reward | -0.0004 | 0.0006 | 11937.12 | -0.0015 | 0.0007 | -0.68 | 0.49354 |
| punish - shapes | 0.0016 | 0.0006 | 11995.27 | 0.0005 | 0.0027 | 2.75 | 0.00600 |
| punish - story | -0.0002 | 0.0006 | 12020.13 | -0.0013 | 0.0010 | -0.29 | 0.77555 |
| punish - ToM | 0.0001 | 0.0006 | 11996.42 | -0.0010 | 0.0012 | 0.14 | 0.88871 |
| random - relational | -0.0001 | 0.0006 | 11966.06 | -0.0012 | 0.0011 | -0.11 | 0.91139 |
| random - reward | -0.0012 | 0.0006 | 11996.42 | -0.0024 | -0.0001 | -2.18 | 0.02948 |
| random - shapes | 0.0007 | 0.0006 | 11956.08 | -0.0004 | 0.0019 | 1.23 | 0.21707 |
| random - story | -0.0010 | 0.0006 | 11987.21 | -0.0022 | 0.0001 | -1.74 | 0.08233 |
| random - ToM | -0.0008 | 0.0006 | 11937.12 | -0.0019 | 0.0004 | -1.35 | 0.17713 |
| relational - reward | -0.0012 | 0.0006 | 12002.80 | -0.0023 | -0.0001 | -2.05 | 0.04016 |
| relational - shapes | 0.0008 | 0.0006 | 11950.73 | -0.0004 | 0.0019 | 1.34 | 0.18050 |
| relational - story | -0.0010 | 0.0006 | 11975.22 | -0.0021 | 0.0002 | -1.62 | 0.10519 |
| relational - ToM | -0.0007 | 0.0006 | 11966.06 | -0.0019 | 0.0004 | -1.23 | 0.21887 |
| reward - shapes | 0.0020 | 0.0006 | 11995.27 | 0.0008 | 0.0031 | 3.42 | 0.00062 |
| reward - story | 0.0002 | 0.0006 | 12020.13 | -0.0009 | 0.0014 | 0.38 | 0.70721 |
| reward - ToM | 0.0005 | 0.0006 | 11996.42 | -0.0007 | 0.0016 | 0.82 | 0.41485 |
| shapes - story | -0.0017 | 0.0006 | 11977.40 | -0.0029 | -0.0006 | -2.95 | 0.00322 |
| shapes - ToM | -0.0015 | 0.0006 | 11956.08 | -0.0026 | -0.0004 | -2.58 | 0.00980 |
| story - ToM | 0.0002 | 0.0006 | 11987.21 | -0.0009 | 0.0014 | 0.42 | 0.67595 |

**SI Table 9.4** Extraversion* task condition interactions along D3 (DMN- FPN)

| contrast | β | SE | df | $CL_{lower}$ | $CL_{upper}$ | t.ratio | p.value |
|---|---|---|---|---|---|---|---|
| 0back - 2back | 0.0004 | 0.0006 | 11937.12 | -0.0007 | 0.0015 | 0.75 | 0.45031 |
| 0back - faces | -0.0002 | 0.0006 | 11981.63 | -0.0013 | 0.0009 | -0.37 | 0.71416 |
| 0back - match | -0.0004 | 0.0006 | 11992.15 | -0.0015 | 0.0007 | -0.77 | 0.44284 |
| 0back - math | 0.0001 | 0.0006 | 12007.53 | -0.0010 | 0.0012 | 0.17 | 0.86594 |
| 0back - motor | 0.0004 | 0.0006 | 11960.83 | -0.0007 | 0.0015 | 0.79 | 0.42668 |
| 0back - punish | -0.0011 | 0.0006 | 11950.21 | -0.0022 | 0.0000 | -1.95 | 0.05174 |
| 0back - random | 0.0002 | 0.0006 | 11982.70 | -0.0009 | 0.0013 | 0.34 | 0.73319 |
| 0back - relational | -0.0002 | 0.0006 | 11992.15 | -0.0014 | 0.0009 | -0.44 | 0.65802 |
| 0back - reward | -0.0008 | 0.0006 | 11950.21 | -0.0019 | 0.0002 | -1.53 | 0.12625 |
| 0back - shapes | -0.0010 | 0.0006 | 11981.63 | -0.0021 | 0.0000 | -1.88 | 0.06026 |
| 0back - story | -0.0001 | 0.0006 | 12007.53 | -0.0012 | 0.0010 | -0.15 | 0.87894 |
| 0back - ToM | -0.0006 | 0.0006 | 11982.70 | -0.0017 | 0.0005 | -1.13 | 0.25676 |
| 2back - faces | -0.0006 | 0.0006 | 11981.63 | -0.0017 | 0.0005 | -1.12 | 0.26421 |
| 2back - match | -0.0008 | 0.0006 | 11992.15 | -0.0020 | 0.0003 | -1.51 | 0.13053 |
| 2back - math | -0.0003 | 0.0006 | 12007.53 | -0.0014 | 0.0008 | -0.57 | 0.57071 |
| 2back - motor | 0.0000 | 0.0006 | 11960.83 | -0.0011 | 0.0011 | 0.05 | 0.96353 |
| 2back - punish | -0.0015 | 0.0006 | 11950.21 | -0.0026 | -0.0004 | -2.70 | 0.00693 |
| 2back - random | -0.0002 | 0.0006 | 11982.70 | -0.0013 | 0.0009 | -0.41 | 0.68130 |
| 2back - relational | -0.0007 | 0.0006 | 11992.15 | -0.0018 | 0.0004 | -1.19 | 0.23511 |
| 2back - reward | -0.0013 | 0.0006 | 11950.21 | -0.0024 | -0.0002 | -2.28 | 0.02236 |
| 2back - shapes | -0.0015 | 0.0006 | 11981.63 | -0.0026 | -0.0004 | -2.63 | 0.00857 |
| 2back - story | -0.0005 | 0.0006 | 12007.53 | -0.0016 | 0.0006 | -0.89 | 0.37447 |
| 2back - ToM | -0.0011 | 0.0006 | 11982.70 | -0.0021 | 0.0000 | -1.89 | 0.05935 |
| faces - match | -0.0002 | 0.0006 | 11955.59 | -0.0013 | 0.0009 | -0.40 | 0.68717 |
| faces - math | 0.0003 | 0.0006 | 11978.58 | -0.0008 | 0.0014 | 0.53 | 0.59860 |
| faces - motor | 0.0006 | 0.0006 | 11989.43 | -0.0005 | 0.0017 | 1.15 | 0.24833 |
| faces - punish | -0.0009 | 0.0006 | 11978.19 | -0.0020 | 0.0002 | -1.57 | 0.11687 |
| faces - random | 0.0004 | 0.0006 | 11954.02 | -0.0007 | 0.0015 | 0.71 | 0.48053 |
| faces - relational | 0.0000 | 0.0006 | 11955.59 | -0.0011 | 0.0011 | -0.08 | 0.93714 |
| faces - reward | -0.0006 | 0.0006 | 11978.19 | -0.0017 | 0.0004 | -1.15 | 0.24849 |
| faces - shapes | -0.0008 | 0.0006 | 11937.12 | -0.0019 | 0.0003 | -1.51 | 0.13137 |
| faces - story | 0.0001 | 0.0006 | 11978.58 | -0.0010 | 0.0012 | 0.21 | 0.83657 |
| faces - ToM | -0.0004 | 0.0006 | 11954.02 | -0.0015 | 0.0007 | -0.76 | 0.44449 |
| match - math | 0.0005 | 0.0006 | 11966.42 | -0.0006 | 0.0017 | 0.92 | 0.35834 |
| match - motor | 0.0009 | 0.0006 | 11982.91 | -0.0002 | 0.0020 | 1.55 | 0.12157 |
| match - punish | -0.0006 | 0.0006 | 11988.62 | -0.0017 | 0.0005 | -1.15 | 0.24930 |
| match - random | 0.0006 | 0.0006 | 11964.16 | -0.0005 | 0.0017 | 1.10 | 0.26982 |
| match - relational | 0.0002 | 0.0006 | 11937.12 | -0.0009 | 0.0013 | 0.32 | 0.74753 |
| match - reward | -0.0004 | 0.0006 | 11988.62 | -0.0015 | 0.0007 | -0.74 | 0.45864 |

| | | | | | | | |
|---|---|---|---|---|---|---|---|
| match - shapes | -0.0006 | 0.0006 | 11955.59 | -0.0017 | 0.0005 | -1.09 | 0.27373 |
| match - story | 0.0003 | 0.0006 | 11966.42 | -0.0008 | 0.0015 | 0.60 | 0.54826 |
| match - ToM | -0.0002 | 0.0006 | 11964.16 | -0.0013 | 0.0009 | -0.36 | 0.72203 |
| math - motor | 0.0003 | 0.0006 | 12001.32 | -0.0008 | 0.0015 | 0.61 | 0.54300 |
| math - punish | -0.0012 | 0.0006 | 12004.38 | -0.0023 | -0.0001 | -2.07 | 0.03878 |
| math - random | 0.0001 | 0.0006 | 11974.73 | -0.0010 | 0.0012 | 0.16 | 0.86917 |
| math - relational | -0.0003 | 0.0006 | 11966.42 | -0.0015 | 0.0008 | -0.60 | 0.54810 |
| math - reward | -0.0009 | 0.0006 | 12004.38 | -0.0021 | 0.0002 | -1.66 | 0.09682 |
| math - shapes | -0.0011 | 0.0006 | 11978.58 | -0.0023 | 0.0000 | -2.01 | 0.04493 |
| math - story | -0.0002 | 0.0006 | 11937.12 | -0.0013 | 0.0010 | -0.32 | 0.75260 |
| math - ToM | -0.0007 | 0.0006 | 11974.73 | -0.0018 | 0.0004 | -1.28 | 0.20143 |
| motor - punish | -0.0015 | 0.0006 | 11956.03 | -0.0026 | -0.0004 | -2.73 | 0.00638 |
| motor - random | -0.0003 | 0.0006 | 11994.21 | -0.0014 | 0.0008 | -0.45 | 0.65027 |
| motor - relational | -0.0007 | 0.0006 | 11982.91 | -0.0018 | 0.0004 | -1.23 | 0.22042 |
| motor - reward | -0.0013 | 0.0006 | 11956.03 | -0.0024 | -0.0002 | -2.31 | 0.02065 |
| motor - shapes | -0.0015 | 0.0006 | 11989.43 | -0.0026 | -0.0004 | -2.66 | 0.00789 |
| motor - story | -0.0005 | 0.0006 | 12001.32 | -0.0017 | 0.0006 | -0.93 | 0.35367 |
| motor - ToM | -0.0011 | 0.0006 | 11994.21 | -0.0022 | 0.0000 | -1.92 | 0.05511 |
| punish - random | 0.0013 | 0.0006 | 11982.70 | 0.0002 | 0.0024 | 2.28 | 0.02266 |
| punish - relational | 0.0008 | 0.0006 | 11988.62 | -0.0003 | 0.0019 | 1.48 | 0.13963 |
| punish - reward | 0.0002 | 0.0006 | 11937.12 | -0.0009 | 0.0013 | 0.42 | 0.67666 |
| punish - shapes | 0.0000 | 0.0006 | 11978.19 | -0.0011 | 0.0011 | 0.05 | 0.95744 |
| punish - story | 0.0010 | 0.0006 | 12004.38 | -0.0001 | 0.0021 | 1.75 | 0.08099 |
| punish - ToM | 0.0004 | 0.0006 | 11982.70 | -0.0006 | 0.0015 | 0.80 | 0.42220 |
| random - relational | -0.0004 | 0.0006 | 11964.16 | -0.0015 | 0.0007 | -0.78 | 0.43585 |
| random - reward | -0.0010 | 0.0006 | 11982.70 | -0.0021 | 0.0001 | -1.86 | 0.06225 |
| random - shapes | -0.0012 | 0.0006 | 11954.02 | -0.0023 | -0.0001 | -2.22 | 0.02671 |
| random - story | -0.0003 | 0.0006 | 11974.73 | -0.0014 | 0.0008 | -0.49 | 0.62737 |
| random - ToM | -0.0008 | 0.0006 | 11937.12 | -0.0019 | 0.0003 | -1.47 | 0.14064 |
| relational - reward | -0.0006 | 0.0006 | 11988.62 | -0.0017 | 0.0005 | -1.07 | 0.28631 |
| relational - shapes | -0.0008 | 0.0006 | 11955.59 | -0.0019 | 0.0003 | -1.42 | 0.15610 |
| relational - story | 0.0002 | 0.0006 | 11966.42 | -0.0010 | 0.0013 | 0.28 | 0.77760 |
| relational - ToM | -0.0004 | 0.0006 | 11964.16 | -0.0015 | 0.0007 | -0.68 | 0.49650 |
| reward - shapes | -0.0002 | 0.0006 | 11978.19 | -0.0013 | 0.0009 | -0.36 | 0.71833 |
| reward - story | 0.0008 | 0.0006 | 12004.38 | -0.0004 | 0.0019 | 1.34 | 0.18058 |
| reward - ToM | 0.0002 | 0.0006 | 11982.70 | -0.0009 | 0.0013 | 0.39 | 0.69811 |
| shapes - story | 0.0010 | 0.0006 | 11978.58 | -0.0002 | 0.0021 | 1.69 | 0.09194 |
| shapes - ToM | 0.0004 | 0.0006 | 11954.02 | -0.0007 | 0.0015 | 0.75 | 0.45579 |
| story - ToM | -0.0005 | 0.0006 | 11974.73 | -0.0017 | 0.0006 | -0.96 | 0.33868 |

**SI Table 9.5** Agreeableness* task condition interactions along D3 (DMN- FPN)

| contrast | β | SE | df | CL lower | CL upper | t.ratio | p.value |
|---|---|---|---|---|---|---|---|
| 0back - 2back | -0.0004 | 0.0006 | 11937.12 | -0.0015 | 0.0007 | -0.74 | 0.45790 |
| 0back - faces | -0.0011 | 0.0006 | 11974.44 | -0.0023 | 0.0000 | -2.04 | 0.04178 |
| 0back - match | -0.0009 | 0.0006 | 11981.03 | -0.0020 | 0.0003 | -1.51 | 0.13212 |
| 0back - math | -0.0007 | 0.0006 | 12012.93 | -0.0018 | 0.0005 | -1.13 | 0.26029 |
| 0back - motor | -0.0015 | 0.0006 | 11954.40 | -0.0026 | -0.0004 | -2.74 | 0.00618 |
| 0back - punish | -0.0005 | 0.0006 | 11952.55 | -0.0016 | 0.0006 | -0.98 | 0.32907 |
| 0back - random | -0.0002 | 0.0006 | 11986.08 | -0.0013 | 0.0009 | -0.32 | 0.75025 |
| 0back - relational | -0.0011 | 0.0006 | 11981.03 | -0.0022 | 0.0001 | -1.86 | 0.06350 |
| 0back - reward | -0.0005 | 0.0006 | 11952.55 | -0.0016 | 0.0006 | -0.95 | 0.33987 |
| 0back - shapes | -0.0008 | 0.0006 | 11974.44 | -0.0019 | 0.0003 | -1.35 | 0.17696 |
| 0back - story | -0.0005 | 0.0006 | 12012.93 | -0.0016 | 0.0006 | -0.85 | 0.39287 |
| 0back - ToM | 0.0006 | 0.0006 | 11986.08 | -0.0005 | 0.0017 | 0.99 | 0.32014 |
| 2back - faces | -0.0007 | 0.0006 | 11974.44 | -0.0018 | 0.0004 | -1.30 | 0.19397 |
| 2back - match | -0.0004 | 0.0006 | 11981.03 | -0.0015 | 0.0007 | -0.77 | 0.43972 |
| 2back - math | -0.0002 | 0.0006 | 12012.93 | -0.0014 | 0.0009 | -0.41 | 0.68164 |
| 2back - motor | -0.0011 | 0.0006 | 11954.40 | -0.0022 | 0.0000 | -2.00 | 0.04541 |
| 2back - punish | -0.0001 | 0.0006 | 11952.56 | -0.0012 | 0.0010 | -0.23 | 0.81464 |
| 2back - random | 0.0002 | 0.0006 | 11986.09 | -0.0009 | 0.0013 | 0.42 | 0.67373 |
| 2back - relational | -0.0006 | 0.0006 | 11981.03 | -0.0017 | 0.0005 | -1.12 | 0.26162 |
| 2back - reward | -0.0001 | 0.0006 | 11952.56 | -0.0012 | 0.0010 | -0.21 | 0.83144 |
| 2back - shapes | -0.0003 | 0.0006 | 11974.44 | -0.0014 | 0.0008 | -0.61 | 0.53969 |
| 2back - story | -0.0001 | 0.0006 | 12012.93 | -0.0012 | 0.0011 | -0.14 | 0.88950 |
| 2back - ToM | 0.0010 | 0.0006 | 11986.09 | -0.0001 | 0.0021 | 1.73 | 0.08302 |
| faces - match | 0.0003 | 0.0006 | 11951.28 | -0.0008 | 0.0014 | 0.52 | 0.60497 |
| faces - math | 0.0005 | 0.0006 | 11988.84 | -0.0006 | 0.0016 | 0.85 | 0.39700 |
| faces - motor | -0.0004 | 0.0006 | 11982.16 | -0.0015 | 0.0007 | -0.70 | 0.48634 |
| faces - punish | 0.0006 | 0.0006 | 11971.14 | -0.0005 | 0.0017 | 1.07 | 0.28656 |
| faces - random | 0.0010 | 0.0006 | 11964.19 | -0.0001 | 0.0021 | 1.72 | 0.08627 |
| faces - relational | 0.0001 | 0.0006 | 11951.28 | -0.0010 | 0.0012 | 0.17 | 0.86580 |
| faces - reward | 0.0006 | 0.0006 | 11971.14 | -0.0005 | 0.0017 | 1.09 | 0.27698 |
| faces - shapes | 0.0004 | 0.0006 | 11937.12 | -0.0007 | 0.0015 | 0.68 | 0.49482 |
| faces - story | 0.0007 | 0.0006 | 11988.84 | -0.0005 | 0.0018 | 1.12 | 0.26399 |
| faces - ToM | 0.0017 | 0.0006 | 11964.19 | 0.0006 | 0.0028 | 3.02 | 0.00252 |
| match - math | 0.0002 | 0.0006 | 11983.72 | -0.0009 | 0.0013 | 0.34 | 0.73332 |
| match - motor | -0.0007 | 0.0006 | 11976.57 | -0.0018 | 0.0004 | -1.21 | 0.22634 |
| match - punish | 0.0003 | 0.0006 | 11978.78 | -0.0008 | 0.0014 | 0.54 | 0.58874 |
| match - random | 0.0007 | 0.0006 | 11974.26 | -0.0004 | 0.0018 | 1.19 | 0.23493 |
| match - relational | -0.0002 | 0.0006 | 11937.12 | -0.0013 | 0.0009 | -0.35 | 0.72871 |
| match - reward | 0.0003 | 0.0006 | 11978.78 | -0.0008 | 0.0014 | 0.56 | 0.57412 |

| | | | | | | | |
|---|---|---|---|---|---|---|---|
| match - shapes | 0.0001 | 0.0006 | 11951.28 | -0.0010 | 0.0012 | 0.16 | 0.87151 |
| match - story | 0.0004 | 0.0006 | 11983.72 | -0.0008 | 0.0015 | 0.61 | 0.54215 |
| match - ToM | 0.0014 | 0.0006 | 11974.26 | 0.0003 | 0.0025 | 2.49 | 0.01290 |
| math - motor | -0.0009 | 0.0006 | 12009.56 | -0.0020 | 0.0003 | -1.52 | 0.12779 |
| math - punish | 0.0001 | 0.0006 | 12007.02 | -0.0010 | 0.0012 | 0.18 | 0.85402 |
| math - random | 0.0005 | 0.0006 | 11995.59 | -0.0007 | 0.0016 | 0.82 | 0.41423 |
| math - relational | -0.0004 | 0.0006 | 11983.72 | -0.0015 | 0.0007 | -0.68 | 0.49707 |
| math - reward | 0.0001 | 0.0006 | 12007.02 | -0.0010 | 0.0013 | 0.20 | 0.83772 |
| math - shapes | -0.0001 | 0.0006 | 11988.84 | -0.0013 | 0.0010 | -0.18 | 0.85374 |
| math - story | 0.0002 | 0.0006 | 11937.12 | -0.0010 | 0.0013 | 0.26 | 0.79197 |
| math - ToM | 0.0012 | 0.0006 | 11995.59 | 0.0001 | 0.0024 | 2.08 | 0.03711 |
| motor - punish | 0.0010 | 0.0006 | 11956.36 | -0.0001 | 0.0021 | 1.77 | 0.07726 |
| motor - random | 0.0014 | 0.0006 | 11998.01 | 0.0003 | 0.0025 | 2.41 | 0.01589 |
| motor - relational | 0.0005 | 0.0006 | 11976.57 | -0.0006 | 0.0016 | 0.86 | 0.38878 |
| motor - reward | 0.0010 | 0.0006 | 11956.36 | -0.0001 | 0.0021 | 1.79 | 0.07373 |
| motor - shapes | 0.0008 | 0.0006 | 11982.16 | -0.0003 | 0.0019 | 1.38 | 0.16830 |
| motor - story | 0.0010 | 0.0006 | 12009.56 | -0.0001 | 0.0022 | 1.79 | 0.07302 |
| motor - ToM | 0.0021 | 0.0006 | 11998.01 | 0.0010 | 0.0032 | 3.72 | 0.00020 |
| punish - random | 0.0004 | 0.0006 | 11985.17 | -0.0007 | 0.0015 | 0.65 | 0.51277 |
| punish - relational | -0.0005 | 0.0006 | 11978.78 | -0.0016 | 0.0006 | -0.89 | 0.37323 |
| punish - reward | 0.0000 | 0.0006 | 11937.12 | -0.0011 | 0.0011 | 0.02 | 0.98278 |
| punish - shapes | -0.0002 | 0.0006 | 11971.14 | -0.0013 | 0.0009 | -0.38 | 0.70379 |
| punish - story | 0.0001 | 0.0006 | 12007.02 | -0.0011 | 0.0012 | 0.09 | 0.93047 |
| punish - ToM | 0.0011 | 0.0006 | 11985.17 | 0.0000 | 0.0022 | 1.97 | 0.04925 |
| random - relational | -0.0009 | 0.0006 | 11974.26 | -0.0020 | 0.0002 | -1.54 | 0.12431 |
| random - reward | -0.0004 | 0.0006 | 11985.17 | -0.0015 | 0.0007 | -0.63 | 0.52672 |
| random - shapes | -0.0006 | 0.0006 | 11964.19 | -0.0017 | 0.0005 | -1.03 | 0.30251 |
| random - story | -0.0003 | 0.0006 | 11995.59 | -0.0015 | 0.0008 | -0.55 | 0.58535 |
| random - ToM | 0.0007 | 0.0006 | 11937.12 | -0.0004 | 0.0018 | 1.31 | 0.18967 |
| relational - reward | 0.0005 | 0.0006 | 11978.78 | -0.0006 | 0.0016 | 0.91 | 0.36189 |
| relational - shapes | 0.0003 | 0.0006 | 11951.28 | -0.0008 | 0.0014 | 0.51 | 0.61004 |
| relational - story | 0.0006 | 0.0006 | 11983.72 | -0.0006 | 0.0017 | 0.95 | 0.34315 |
| relational - ToM | 0.0016 | 0.0006 | 11974.26 | 0.0005 | 0.0027 | 2.84 | 0.00458 |
| reward - shapes | -0.0002 | 0.0006 | 11971.14 | -0.0013 | 0.0009 | -0.40 | 0.68794 |
| reward - story | 0.0000 | 0.0006 | 12007.02 | -0.0011 | 0.0012 | 0.07 | 0.94704 |
| reward - ToM | 0.0011 | 0.0006 | 11985.17 | 0.0000 | 0.0022 | 1.95 | 0.05179 |
| shapes - story | 0.0003 | 0.0006 | 11988.84 | -0.0009 | 0.0014 | 0.45 | 0.64955 |
| shapes - ToM | 0.0013 | 0.0006 | 11964.19 | 0.0002 | 0.0024 | 2.34 | 0.01946 |
| story - ToM | 0.0011 | 0.0006 | 11995.59 | -0.0001 | 0.0022 | 1.81 | 0.06972 |
